# Conservation of gene co-expression networks and neuropsychiatric-neurological disease gene programs between mouse and human frontal cortex at single-cell resolution

**DOI:** 10.64898/2026.10.05.756860

**Authors:** Qian-Quan Sun, Tianyu Cao, Chunzhao Zhang

## Abstract

Mouse models are widely used in translational neuroscience, but whether a gene has the same partners in mouse and human prefrontal cortex (PFC) is unknown. The genes whose expression rises and falls with a given gene across samples (its co-expression partners) indicate the molecular program to which it belongs. We compared the co-expression of 13,214 genes between mouse medial PFC and human dorsolateral PFC in 246,211 single-cell and single-nucleus RNA-sequencing profiles. These two regions are not homologous, so part of any measured difference may reflect region and not species. The partners of single genes were only modestly similar in the two species, whereas groups of co-expressed genes (modules) were strongly preserved, and the higher conservation of disease genes was largely accounted for by general gene properties. We report five findings. First, the median similarity of partners for single genes was 0.169 (0, unrelated; 1, identical). This median was below that of every comparison among human datasets but within the range of comparisons among mouse datasets, and correction for measurement noise raised it to 0.349 at most. Second, all four mouse modules and four of five human modules were strongly preserved in the other species. Third, neuropsychiatric disease genes were modestly more conserved than other genes, but no more than control genes matched for mutational constraint and expression. None of the 14 disease genes had fewer cross-species partners than connectivity-matched comparison genes. Fifth, in an independent human dataset, four of five human modules defined on the 5,000 most variable genes recurred. We also identified two sources of error: analyzing only the most variable genes excludes most disease genes, and networks built from few samples, as for glial cells, underestimate the similarity between species. Scores and code are released for cross-species studies.

## Introduction

The prefrontal cortex (PFC) supports working memory, decision making and behavioral flexibility, functions that are disrupted in many neuropsychiatric disorders. Mouse models are widely used in translational studies of these disorders. However, the dorsolateral PFC with a distinct layer 4 (granular dorsolateral PFC) is a specialization of primates [1], rodents lack counterparts of some prefrontal regions of primates [2, 3], and it is unknown which features of human brain disorders can be reproduced in mouse models [4]. Atlases of cortical cell types based on single-cell and single-nucleus RNA sequencing (snRNA-seq) exist for mouse [5, 6] and for human and other primates [7, 8], and direct comparisons show that the major cell classes are shared between mouse and human, whereas marker genes, cell proportions and laminar distributions differ [7, 9, 10]. These comparisons examine mainly the expression of single genes. A gene, however, does not act alone. Genes that take part in the same molecular program rise and fall together in expression across many samples (they are co-expressed), and the genes that follow a given gene most closely (its co-expression partners) indicate the program in which the gene takes part. The partners of a gene can differ between two species even when the gene is expressed in both [11, 12]. Studies of bulk tissue found that human and mouse co-expression networks are similar in overall structure but differ for many individual genes [12, 13], and that groups of co-expressed genes (modules) are conserved between human and chimpanzee brain [11]. Comparative analyses of co-expression now cover up to 37 species [14, 15], and methods for comparing regulatory networks between species have been developed [16]. For the brain, a comparison across 116 datasets from human, mouse and non-human primates found the largest differences in cerebral cortex and more divergence in glial cells than in neurons [4], and a study of five primate species found that fewer than 1% of the genes compared have co-expression that is conserved in non-human animals and strongly divergent in human [17].

For the PFC at the resolution of cell classes, three aspects remain unknown: how similar the partners of an individual gene are in mouse and human, how this similarity varies between cell classes, and whether it is related to a gene’s tolerance of mutations and to its role in disease. A further question is how the measured similarity of co-expression partners between two species should be interpreted. A similarity score obtained between species reflects the species difference, but it also reflects differences between datasets (donors or animals, tissue handling, sequencing chemistry and the procedure used to remove differences between datasets) and measurement noise. It therefore cannot be interpreted without the score obtained when the same analysis compares two datasets of the same species. We introduced such same-species comparisons, together with a correction for measurement noise that is based on the reproducibility of each gene’s score. The regions compared are a further reason for caution. We compared mouse medial PFC (mPFC) with human dorsolateral PFC (DLPFC), the comparison most often made, but the two regions are not homologous [3]. Every score that we report between the species is therefore the score of an mPFC-DLPFC comparison, in which a difference of region is added to the difference of species. This mismatch is expected to lower the similarity scores and not to raise them.

These questions are important for translational research that uses mouse models of neuropsychiatric disorders. Genes that carry risk for neuropsychiatric disorders tolerate damaging mutations poorly (they are under strong mutational constraint) [18, 19, 20, 21], and they are concentrated in particular cell types and co-expression modules of the cortex [22] and of the brain as a whole [23]. Dozens of risk genes for neuropsychiatric and neurodegenerative disorders have been reported to have highly divergent co-expression between mouse and human [4]. If a disease gene has different partners in mouse and human, manipulating it in mouse may engage a different molecular program than in human. Two recent advances in psychiatric genetics are relevant to the choice of disease gene sets for such a comparison. The cross-disorder analysis of the Psychiatric Genomics Consortium studied 14 psychiatric disorders and identified five groups of disorders with shared genetic risk (genomic factors) [24], so gene sets defined by diagnosis mix genes specific to one disorder with genes shared across disorders. In addition, whole-exome sequencing identified 36 genes with high-confidence evidence of risk for obsessive-compulsive disorder (OCD) and chronic tic disorders (CTD) [25], a gene set discovered entirely through rare variants.

In this study we compared the co-expression networks of mouse mPFC and human DLPFC in 246,211 cells and nuclei from seven datasets, one obtained by single-cell RNA sequencing and six by snRNA-seq. We first measured how similar the partners of single genes are in the two species, and how much of the apparent difference comes from measurement noise and from differences between datasets and not from evolution. We then tested whether gene modules are preserved where single genes differ, whether disease genes are more conserved than other genes once general gene properties are taken into account, and whether any disease gene has different partners in the two species when it is compared with genes of similar network connectivity. We also tested the human network in an independent human dataset. During the analysis we found that two common choices, the selection of highly variable genes and the use of small numbers of samples, can lead to false conclusions, and we describe both as general cautions.

## Results

### Data and the two scores of conservation

We combined data from mouse mPFC (107,471 neurons and 14,024 glial cells, 121,495 profiles in total, from one in-house dataset obtained by single-cell RNA sequencing and three public snRNA-seq datasets) and from human DLPFC (93,433 neurons and 31,283 glial cells, 124,716 profiles in total, from three public snRNA-seq datasets; Fig. 1; Table 1). Each profile is one whole cell in the in-house dataset and one nucleus in the public datasets. Co-expression is measured as the correlation between the expression of two genes across samples. Cells or nuclei from the same individual cannot serve as separate samples for this purpose, because they are not independent of one another. We therefore summed the counts of all cells or nuclei of one cell subtype in one sample into a single expression profile (a pseudobulk sample), which gave 180 neuronal pseudobulk samples in mouse and 320 in human. All analyses used 13,214 genes that have a single counterpart in the other species (one-to-one orthologs) and are expressed in both species; we refer to these genes as the gene panel. Results are reported for three broad classes: all neurons, excitatory neurons and inhibitory neurons. Results for single subtypes are not reported, because the assignment of mouse subtypes to human subtypes depends on the method: an assignment by correlation of expression and an assignment by shared marker genes agreed for only 10 of the 21 mouse subtypes (6 of 11 inhibitory subtypes and 4 of 9 excitatory subtypes; the one remaining subtype, which the source study assigned to neither class, did not agree; Fig. S1; Supplementary Note 1).

**Figure 1.**
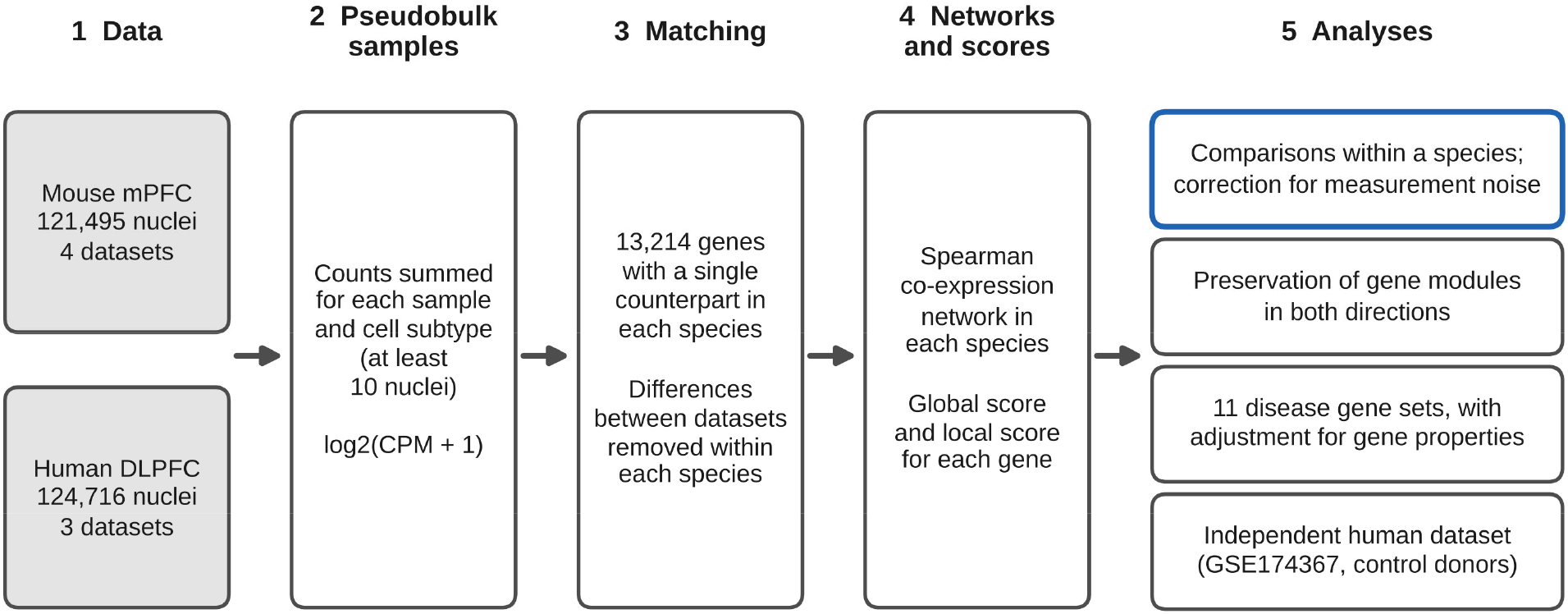
Design of the study. The diagram shows the data (mouse medial prefrontal cortex, mPFC, 121,495 cells and nuclei from four datasets; human dorsolateral prefrontal cortex, DLPFC, 124,716 nuclei from three datasets) and the steps of the analysis: summing of counts into pseudobulk samples for each sample and cell subtype; matching of mouse and human genes through one-to-one orthologs and removal of differences between datasets; a Spearman co-expression network for each species, from which the global score and the local score of each gene are computed; and four analyses (comparisons within a species and correction for measurement noise; the test of module preservation in both directions; the tests of 11 disease gene sets with adjustment for gene properties; and the test of the human network in the control donors of an independent dataset, GSE174367). The first analysis is outlined in blue because it gives the reference values with which the scores between species are interpreted. CPM, counts per million.

**Table 1.** Datasets.

| Data | Source datasets | Cells or nuclei | Cell types | Pseudobulk samples |
| --- | --- | --- | --- | --- |
| Mouse mPFC, neurons | In-house dataset (control mice of the TRAP2 line), GSE248200, GSE291951 and GSE233763 | 107,471 | 21 subtypes | 180 |
| Mouse mPFC, glial cells | In-house dataset | 14,024 | 5 types | 74 |
| Human DLPFC, neurons | GSE243292, GSE207334 and GSE261983 | 93,433 | 18 subtypes | 320 |
| Human DLPFC, glial cells | GSE243292 (control donors) | 31,283 | 5 types | 34 |
| Independent human PFC dataset | GSE174367 (control donors only) | 3,450 (excitatory and inhibitory neurons) | 2 classes | 7 (one per donor) |

A pseudobulk sample is the sum of the counts of all cells or nuclei of one cell subtype in one sample of a source dataset. The in-house mouse dataset was obtained by single-cell RNA sequencing of whole cells, and the public datasets by single-nucleus RNA sequencing. mPFC, medial prefrontal cortex; DLPFC, dorsolateral prefrontal cortex.

Within each species, we computed the correlation of every gene with every other gene across the pseudobulk samples; the genes with which a gene is most strongly correlated are its co-expression partners. We used two scores to describe how similar the partners of a gene are in mouse and human. The global score takes the correlations of the gene with all other genes in mouse and compares them with the same correlations in human, as the Spearman correlation between the two sets of values. It is 0 when the two sets are unrelated and 1 when they are identical. We used the Spearman correlation for this comparison because it depends only on the order of the gene’s correlations with the other genes and is therefore not dominated by a few very strong correlations. The local score considers only the 50 genes most strongly correlated with the gene in each species (its 50 strongest partners) and measures how many of them are shared, as the Jaccard index (the number of partners found in both species divided by the number found in either species). The two scores were correlated across genes (0.51), but one cannot replace the other. In the tests of gene sets described below, we used the average of the two scores of each gene, which we call the conservation score (Methods).

For single genes, the similarity of partners in this mPFC-DLPFC comparison was modest (Fig. 2A; Table 2). The median global score was 0.169 in all neurons, 0.108 in excitatory neurons and 0.057 in inhibitory neurons. The median local score was 0.000 in each of these analyses, because most genes shared few or none of their 50 strongest partners between the species. The scores varied widely among genes: a minority of genes had nearly the same partners in both species, whereas most genes showed weak correspondence. The observed medians nevertheless exceeded the values expected by chance, which we obtained by comparing each gene in mouse with a randomly chosen gene in human (Fig. S2; Supplementary Note 1). The median global score is therefore low, but it is higher than expected by chance.

**Figure 2.**
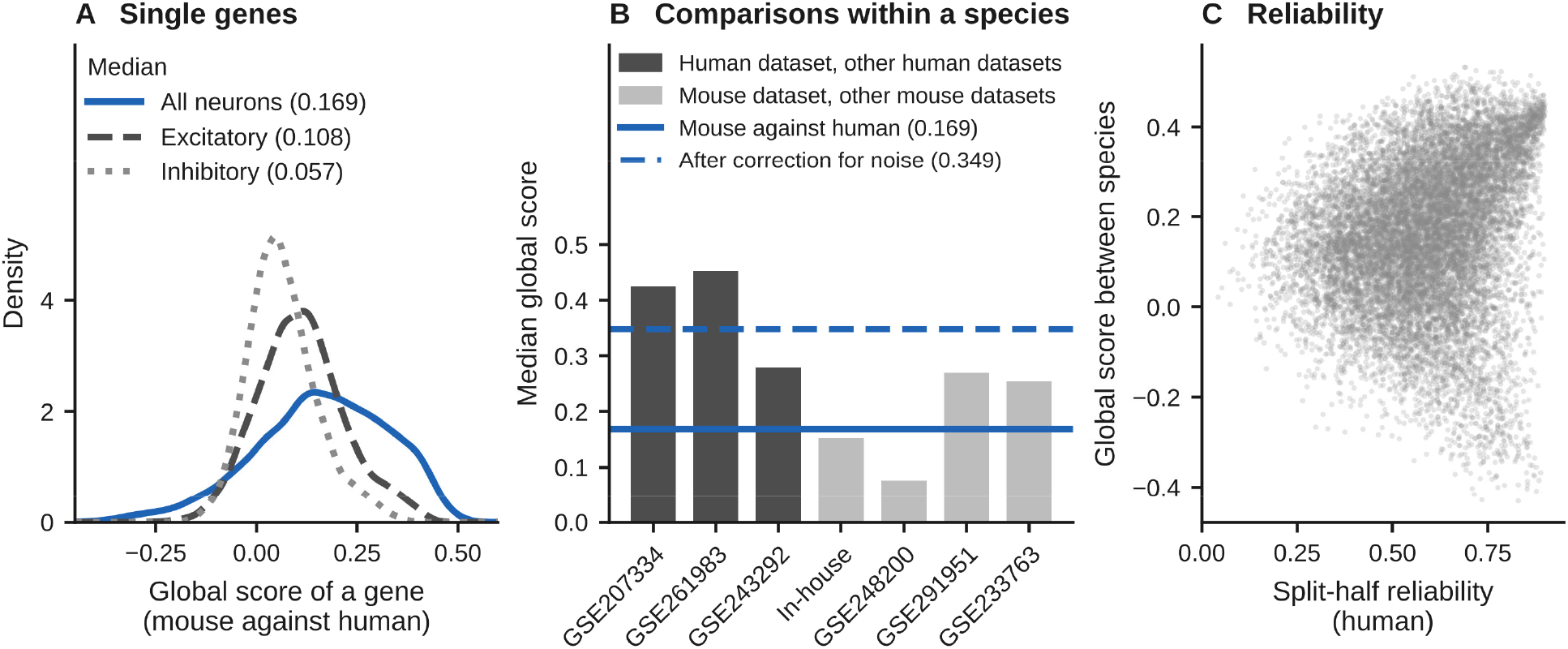
Similarity of co-expression partners between mouse and human, with reference values from comparisons within a species. (A) Distribution of the global score of single genes in all neurons (blue), in excitatory neurons (dashed) and in inhibitory neurons (dotted); medians in parentheses. (B) Median global score when one dataset is compared with the other datasets of the same species (dark grey bars, human, 0.279 to 0.453; light grey bars, mouse, 0.076 to 0.270), with the median between species (blue line, 0.169) and the median after correction for measurement noise (dashed blue line, 0.349). (C) Global score between species of each gene plotted against the split-half reliability of the gene in human; each point is one gene. The median reliabilities are given in Table 2.

**Table 2.**
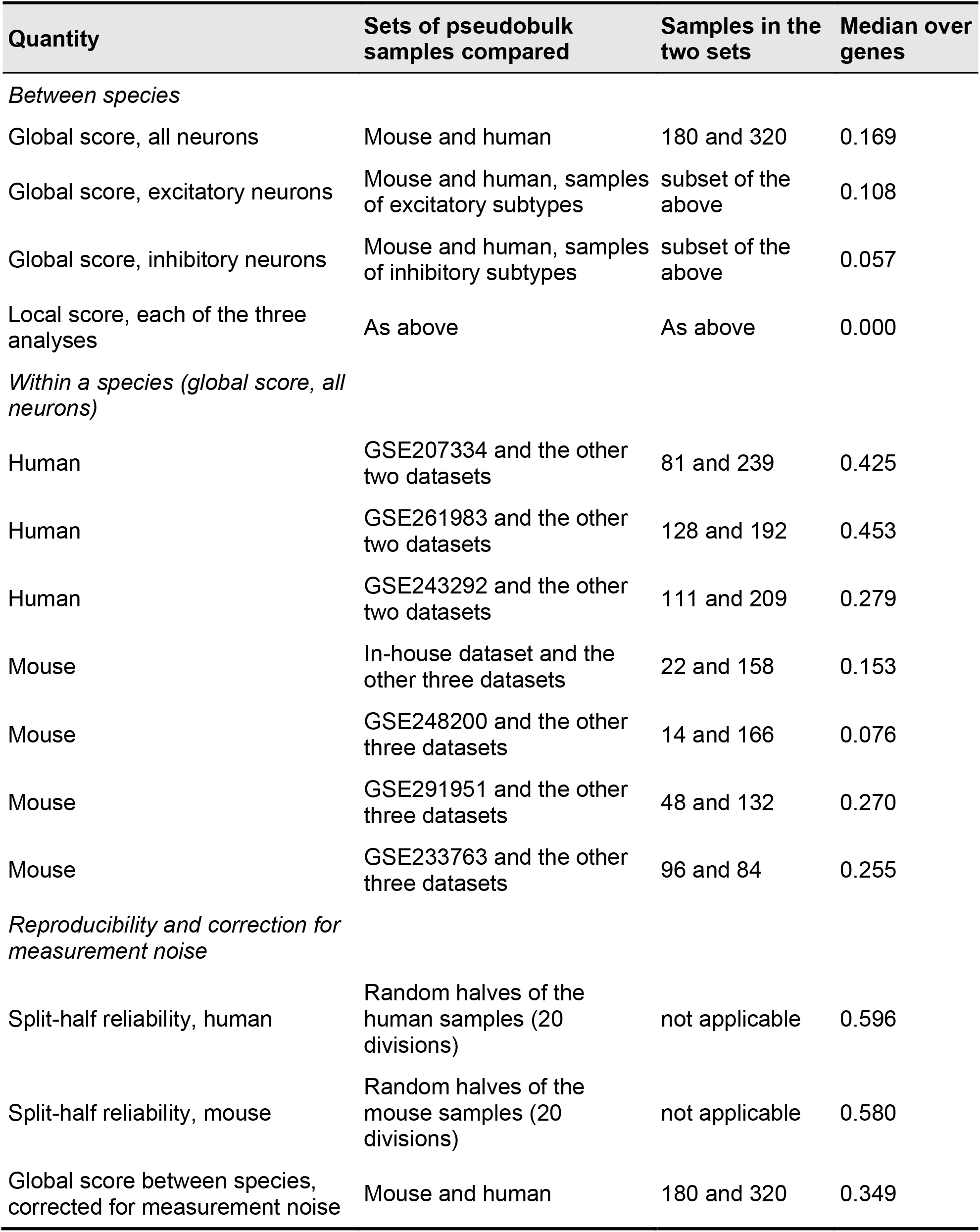
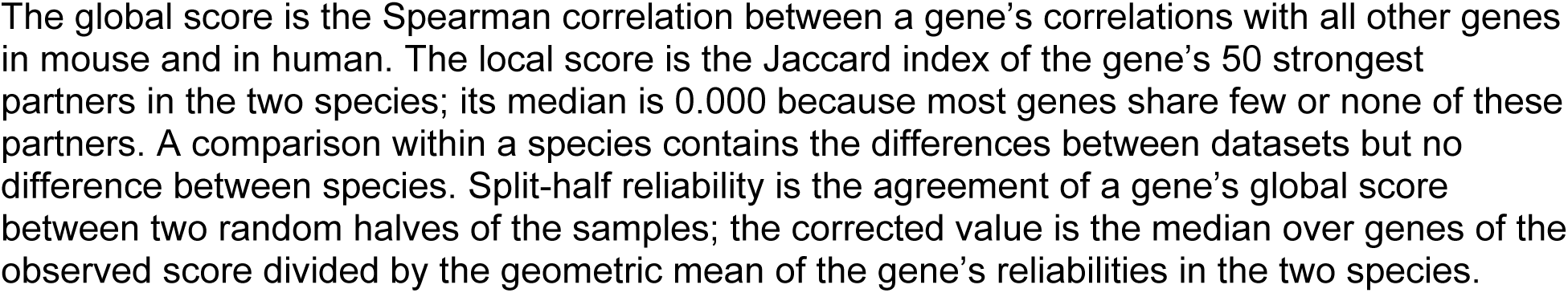
Similarity of co-expression partners of single genes between mouse and human, with reference values from within a species (13,214 genes).

### Reference values for the global score: comparisons within a species and measurement noise

A low score between species could result in part from differences between datasets. To measure this effect, we computed the global score within each species, comparing each dataset with the remaining datasets of the same species (Fig. 2B; Table 2). These same-species comparisons contain every difference between datasets (donors or animals, tissue handling, sequencing chemistry and the procedure used to remove dataset differences) but no species difference. The mouse comparisons, but not the human comparisons, also contain the difference between whole cells and nuclei, because only the in-house mouse dataset consists of whole cells. The median global score was 0.279 to 0.453 in the three human comparisons and 0.076 to 0.270 in the four mouse comparisons. The median between species (0.169) was lower than in every human comparison, as expected if the species differ in addition to the datasets. It was, however, within the range of the mouse comparisons, in which two sets of mouse data gave a median as low as 0.076. The distance between 0.169 and a perfect score of 1 therefore cannot be interpreted as evolutionary divergence.

We next measured how reproducible the score of each gene is. The pseudobulk samples of each species were divided at random into two halves, a network was built in each half, and the global score of each gene was computed between the two halves (split-half reliability; Fig. 2C). The median reliability was 0.596 in human and 0.580 in mouse. Dividing the observed score by the reliability is a standard correction for measurement noise. When we applied this correction to each gene, the median global score rose to 0.349 (the dashed line in Fig. 2B); when we applied it once to the median, the result was 0.287. Measurement noise therefore lowers the observed score to roughly half of its noise-corrected value. Three points limit how these reference values can be used. The same-species comparisons are lowered by the same noise and were not corrected, so the corrected median must not be compared with them; the finding that the median between species lies below the human comparisons is based on uncorrected scores. In some mouse comparisons, one of the two sets of pseudobulk samples is small (14 samples in the smallest), which lowers the score of these comparisons. The correction also assumes that the noise in the two halves is unrelated, and an influence shared by both halves would make the corrected value too high. We therefore take 0.349 as the upper bound of the noise-corrected median and 0.169 as its lower bound.

### Gene modules are strongly preserved

A gene module is a group of genes that are all strongly co-expressed with one another, and it is taken to represent one molecular program. We defined modules in each species with weighted gene co-expression network analysis (WGCNA; Methods). In the network of all neurons, this analysis gave four modules in mouse (3,374, 2,447, 1,866 and 260 genes; the remaining 5,267 genes were not assigned to a module) and five modules in human (1,650, 1,469, 382, 215 and 149 genes; 9,349 genes not assigned). We then tested whether the genes of a module defined in one species are also co-expressed with one another in the other species. The test statistic (Zsummary) compares the module with random groups of genes, and values above 10 are conventionally taken as strong evidence that a module is preserved. All four mouse modules were strongly preserved in human (Zsummary 13.2 to 23.7), and four of the five human modules were strongly preserved in mouse (Zsummary 15.6 to 38.5; Fig. 3A). The exception was the smallest human module (149 genes; Zsummary −0.1). Each mouse module shared more genes than expected by chance with one or two human modules (Fig. 3C); for example, the mouse module of 2,447 genes and the human module of 1,469 genes shared 768 genes. Gene modules are therefore preserved between the species although the partners of single genes are only modestly similar.

**Figure 3.**
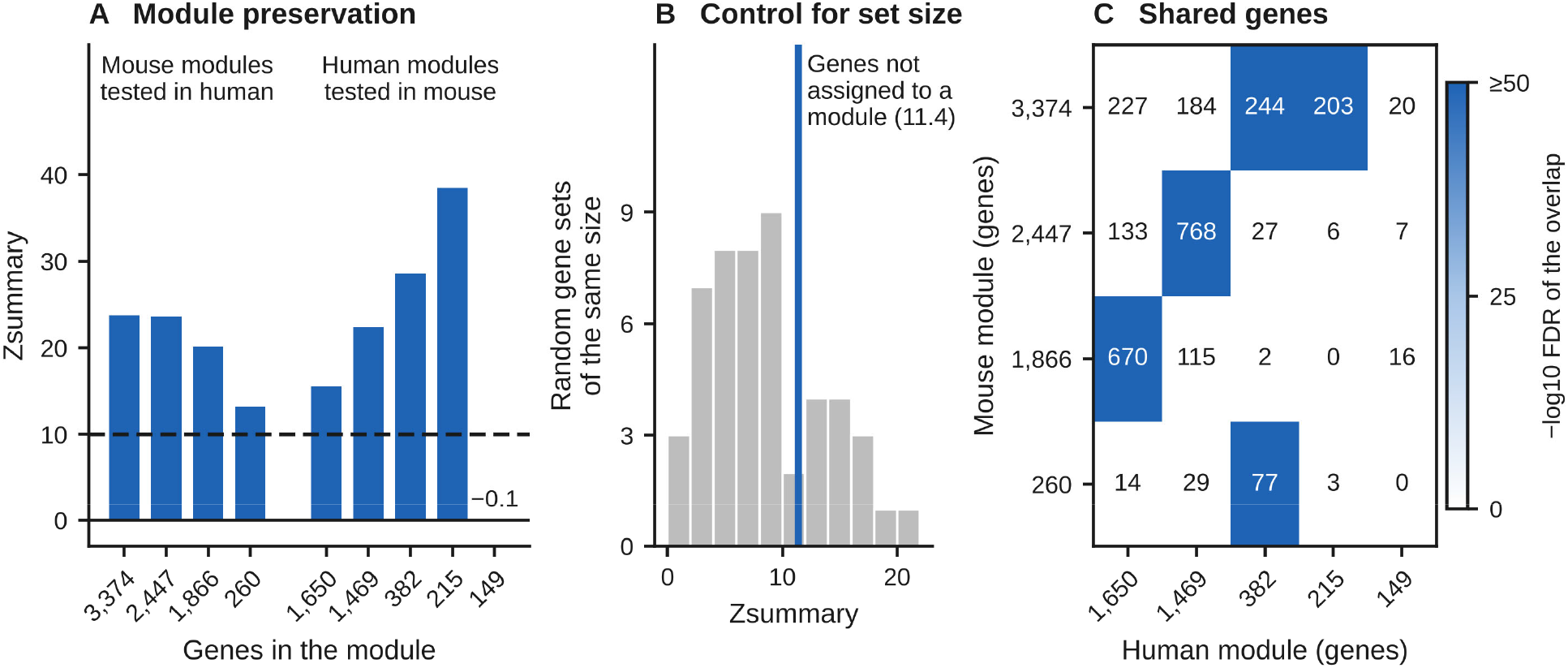
Preservation of gene modules. (A) Preservation of the four mouse modules in human and of the five human modules in mouse (Zsummary). Each module is labeled by its number of genes. Blue bars, Zsummary above 10 (dashed line), the value above which a module is taken as strongly preserved. The value of the human module of 149 genes (−0.1) is printed because its bar is too short to be seen. (B) Zsummary of the mouse genes not assigned to a module (blue line), compared with 50 random gene sets of the same size evaluated with the same statistic (grey). (C) Number of genes shared by each mouse module and each human module. The shading gives the significance of the overlap as −log10 of the false discovery rate (FDR; hypergeometric test); values of 50 or more have the darkest shade, and overlaps no larger than expected by chance are white. The modules are those of the 13,214-gene panel.

One control is needed to interpret this statistic. The 5,267 mouse genes that were not assigned to any module also reached a Zsummary above 10 (11.4). Because Zsummary increases with the number of genes in a set, we evaluated 50 random gene sets of the same size with the same statistic, and they scored between 0.9 and 20.1 (Fig. 3B). The value for the unassigned genes is therefore an effect of the size of the set, and a Zsummary should be interpreted against random gene sets of the same size.

### Disease genes are more conserved than other genes, but not more than matched control genes

Mouse models of brain disorders are often built by manipulating a gene that carries risk for the disorder. Such a model is easier to interpret if the gene has the same co-expression partners in mouse and human, so we asked whether disease genes keep their partners more often than other genes. We tested 11 curated disease gene sets (Methods): seven sets for neuropsychiatric disorders (autism spectrum disorder, schizophrenia, bipolar disorder, attention-deficit/hyperactivity disorder, major depressive disorder, intellectual disability and epilepsy) and four sets for neurological diseases. Together the 11 sets contain 539 genes of the panel. Each set was compared with all other genes of the panel by a rank-based test of the conservation scores. The size of the difference is given as the rank-biserial correlation (r), which is 0 when the genes of the set and the other genes have similar scores and 1 when every gene of the set scores higher than every other gene.

In all neurons, the 539 disease genes were modestly more conserved than other genes (r = 0.102; false discovery rate [FDR] 1.7 × 10⁻⁴; Table 3). Among the single sets, the difference was largest for intellectual disability (r = 0.310), epilepsy (r = 0.250) and autism spectrum disorder (r = 0.234; Fig. 4A). Disease genes, however, differ from other genes in general properties that may themselves be related to conservation. We considered five such properties: the gene’s tolerance of loss-of-function mutations, its mean expression level, its expression variance, its length and its proportion of guanine and cytosine bases (GC content). Tolerance of mutations was measured by the loss-of-function observed/expected upper bound fraction (LOEUF) [26]; a low LOEUF means that fewer loss-of-function variants are observed in the population than expected, that is, the gene is under strong mutational constraint. We took the five properties into account in two ways, and Table 3 shows the results before adjustment, after adjustment and against matched control genes side by side.

**Figure 4.**
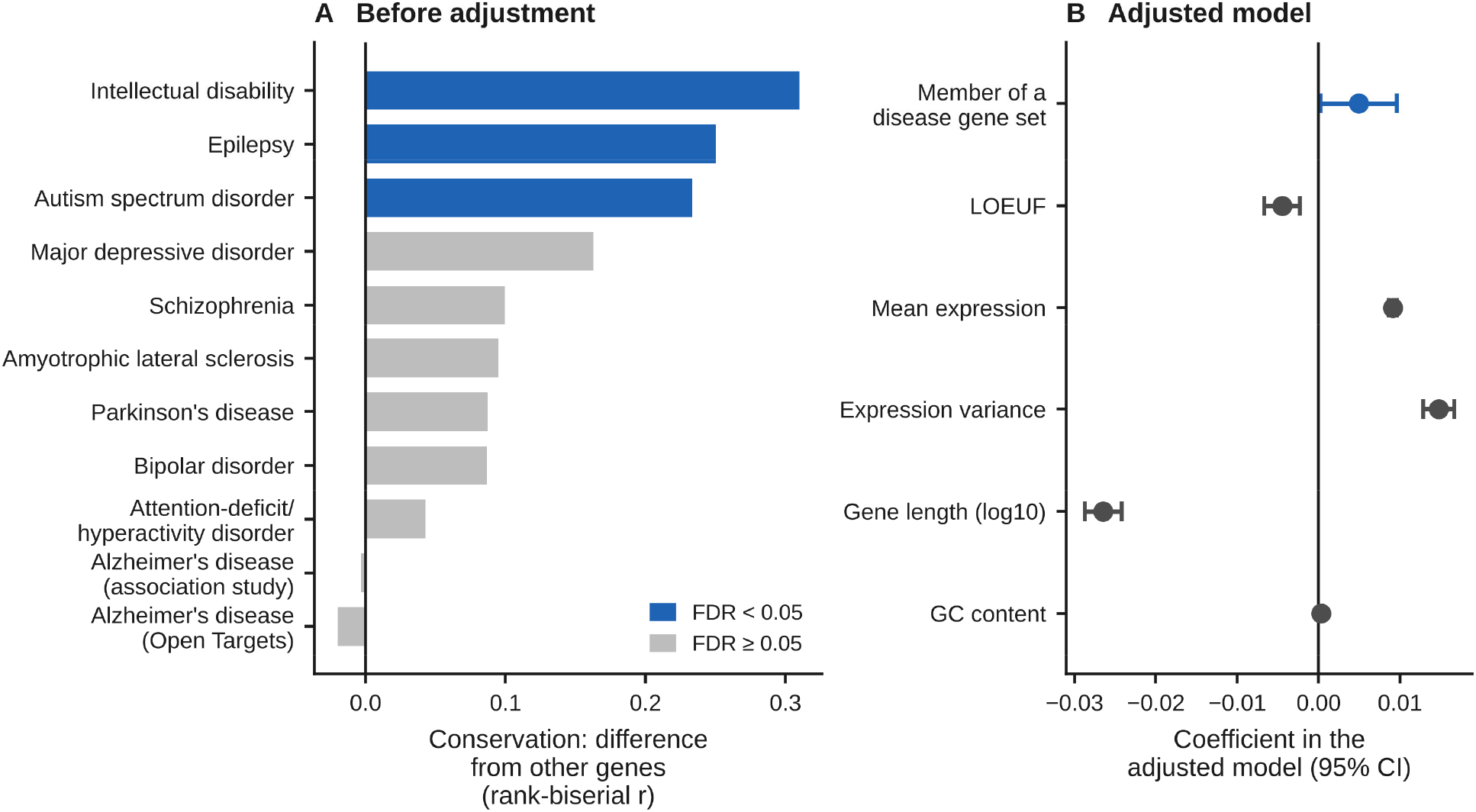
Conservation of disease gene sets before and after adjustment for general gene properties. (A) Difference in the conservation score between each disease gene set and all other genes before adjustment (rank-biserial correlation, r; all neurons). Blue, false discovery rate (FDR) below 0.05. (B) Coefficients of the regression model of the conservation score, with 95% confidence intervals (CI): the coefficient for membership in a disease set (blue) is small (P = 0.039 in a single test that is not corrected for multiple tests), whereas tolerance of loss-of-function mutations (measured by the loss-of-function observed/expected upper bound fraction, LOEUF), mean expression, expression variance, gene length and the proportion of guanine and cytosine bases (GC content) are each associated with conservation. The confidence interval of GC content is narrower than the symbol.

**Table 3.**
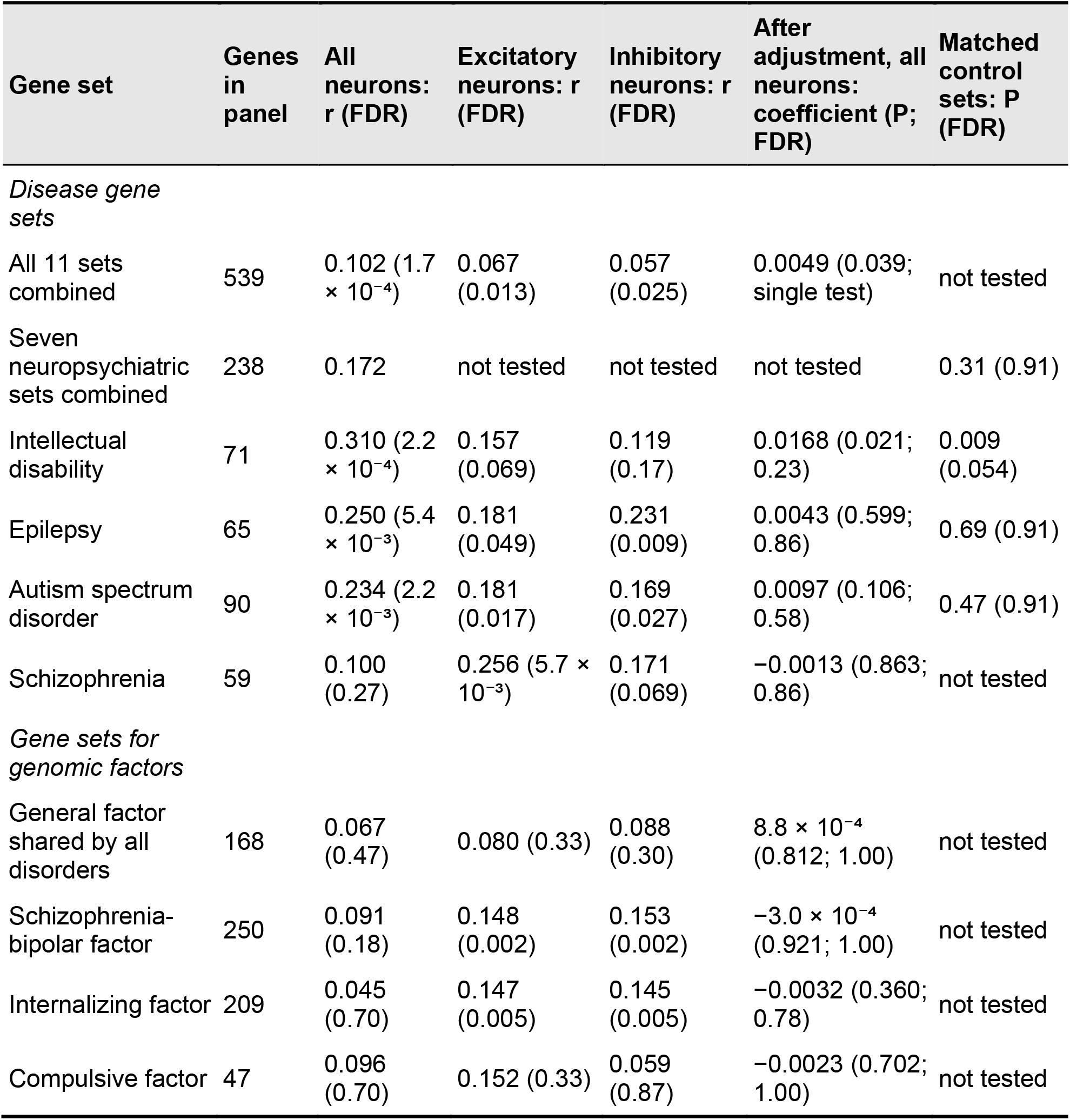
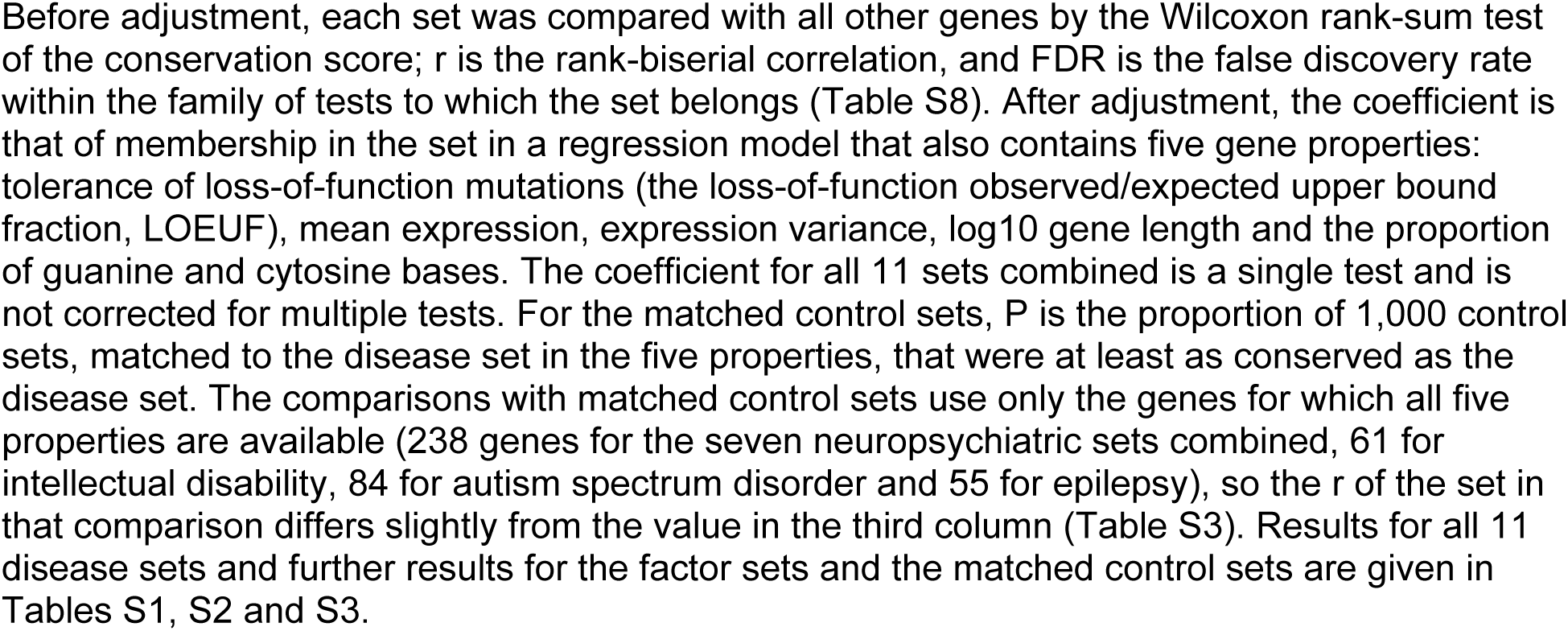
Conservation of disease gene sets and of gene sets for genomic factors: before adjustment, after adjustment and against matched control genes.

The primary analysis compared each disease set with matched control genes. We drew 1,000 control gene sets of the same size, in which every disease gene was replaced by another gene with similar LOEUF, expression level, expression variance, length and GC content, and we asked how often a control set was at least as conserved as the disease set (Fig. 5B). The seven neuropsychiatric sets combined were no more conserved than their matched control sets (r = 0.172, against a mean of 0.159 for the control sets; P = 0.31), and the same was true of the autism set (P = 0.47) and the epilepsy set (P = 0.69). For the combined sets, the difference between the two values (0.013) is small in comparison with the variation among the control sets, 95% of which had values between 0.103 and 0.213 (Fig. 5B). Only the intellectual disability set exceeded its control sets before correction for multiple tests (r = 0.312 against 0.179; P = 0.009). After correction the difference was marginal (FDR = 0.054), and the control sets themselves varied widely (95% interval 0.079 to 0.282). We therefore regard it as an exploratory signal that needs confirmation.

**Figure 5.**
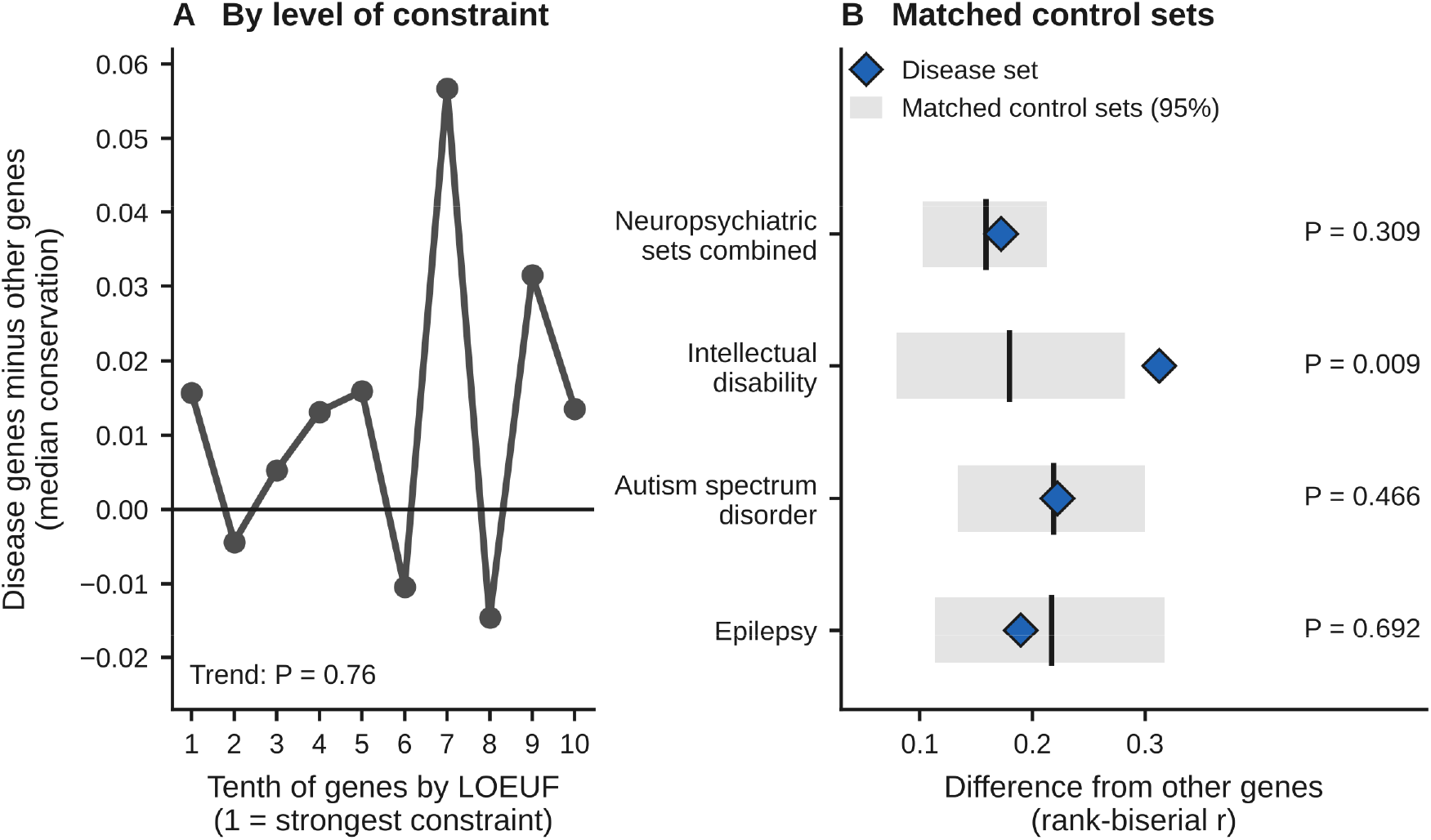
Disease gene sets, mutational constraint and matched control genes. (A) Difference in the median conservation score between the genes of the seven neuropsychiatric sets combined (238 genes) and other genes, within tenths of all genes ordered by the loss-of-function observed/expected upper bound fraction (LOEUF; 1 is the tenth under the strongest constraint); the test for trend has a permutation P of 0.76. (B) Observed difference in the conservation score for each disease set (blue diamonds) compared with the distribution of matched control gene sets (grey bars, 95% interval; black line, mean), with the one-sided P value. The control genes are matched to the disease genes in LOEUF, mean expression, expression variance, gene length and GC content.

A regression model supported this result. In addition to membership in a disease set, the model contained the five gene properties listed above (LOEUF, mean expression level, expression variance, gene length and GC content). In the model, the coefficient for membership of the 539 disease genes in a disease set was 0.0049 (P = 0.039 in this single test, which is not corrected for multiple tests; Fig. 4B). No single set was significant after correction for multiple tests (Table S1; Fig. S3); the smallest FDR was 0.23, for intellectual disability, whose coefficient had a wide confidence interval (0.0168; 95% confidence interval 0.0026 to 0.0310). Mutational constraint itself was associated with conservation (LOEUF coefficient −0.0045; P = 6.8 × 10⁻⁵): genes under strong constraint are more conserved whether or not they are disease genes. The difference between disease genes and other genes did not depend on the level of constraint (interaction P = 0.997), and for the seven neuropsychiatric sets combined it showed no trend across tenths of the genes ordered by LOEUF (P = 0.76; Fig. 5A). Within the tenth under the strongest constraint, which contains 110 of the 238 genes of these sets, the disease genes were nevertheless more conserved than the other genes of that tenth (r = 0.151; P = 0.009). This comparison is a single test that is not corrected for multiple tests, and it holds constraint fixed but not the other gene properties, so we regard its result as exploratory.

The higher conservation of disease genes is therefore real, but it is largely accounted for by the measured gene properties: genes that tolerate mutations poorly and are highly expressed are more conserved for general reasons. This is consistent with the route by which many disease genes are discovered, for example the search for rare damaging variants in patients, which favors genes under strong constraint [18, 19, 20, 21]. The four neurological sets were not more conserved than other genes, but this result is uninformative, because the cell types and regions most affected in these diseases are not those tested here (Supplementary Note 2). The conclusion that the measured gene properties account for the higher conservation of disease genes has three limitations. LOEUF is estimated with error, so adjusting for it removes the influence of constraint only in part, and the small difference that remains after adjustment may still reflect constraint and cannot be assigned to disease genes specifically. LOEUF was also the only measure of constraint that we used. Finally, the analyses show that the five properties together account for the difference, but they do not show how much each property contributes or whether any of them is the cause.

The disease analysis also revealed an analysis choice that can create a false negative result. Disease genes varied less in expression across pseudobulk samples than other genes (median variance 0.294 against 0.690; P = 2.7 × 10⁻²⁴; Fig. S4). A common first step in network analysis is to keep only the most variable genes. When we kept the 5,000 most variable genes, only 46 of the 258 expressed neuropsychiatric disease genes remained (17.8%), and the same test of the conservation of disease genes was no longer significant (r = 0.105; FDR = 0.72), although the size of the difference was similar to that found with all genes of the panel. The negative result was therefore produced entirely by the selection of genes. Network analyses that concern disease genes should therefore test their hypotheses on all expressed genes.

### Gene sets grouped by shared genetic risk are more conserved in excitatory and in inhibitory neurons

Gene sets defined by diagnosis mix genes that are specific to one disorder with genes that affect several disorders. The cross-disorder analysis of the Psychiatric Genomics Consortium [24] provides, in its supplementary tables, lists of the genes linked to each genomic factor and to a general factor that is shared by all disorders. The consortium itself analyzed only the genes of the general factor, the schizophrenia-bipolar factor and the internalizing factor, because few variants were associated with the other factors. Four of these sets contained enough genes of our panel to be tested (Table 3; Fig. 6): the general factor (168 genes), the schizophrenia-bipolar factor (250 genes), the internalizing factor, which groups major depression, anxiety disorders and post-traumatic stress disorder (209 genes), and the compulsive factor, which groups anorexia nervosa and obsessive-compulsive disorder, with weaker contributions of Tourette syndrome and anxiety disorders (47 genes, a small set for which the test is weak).

**Figure 6.**
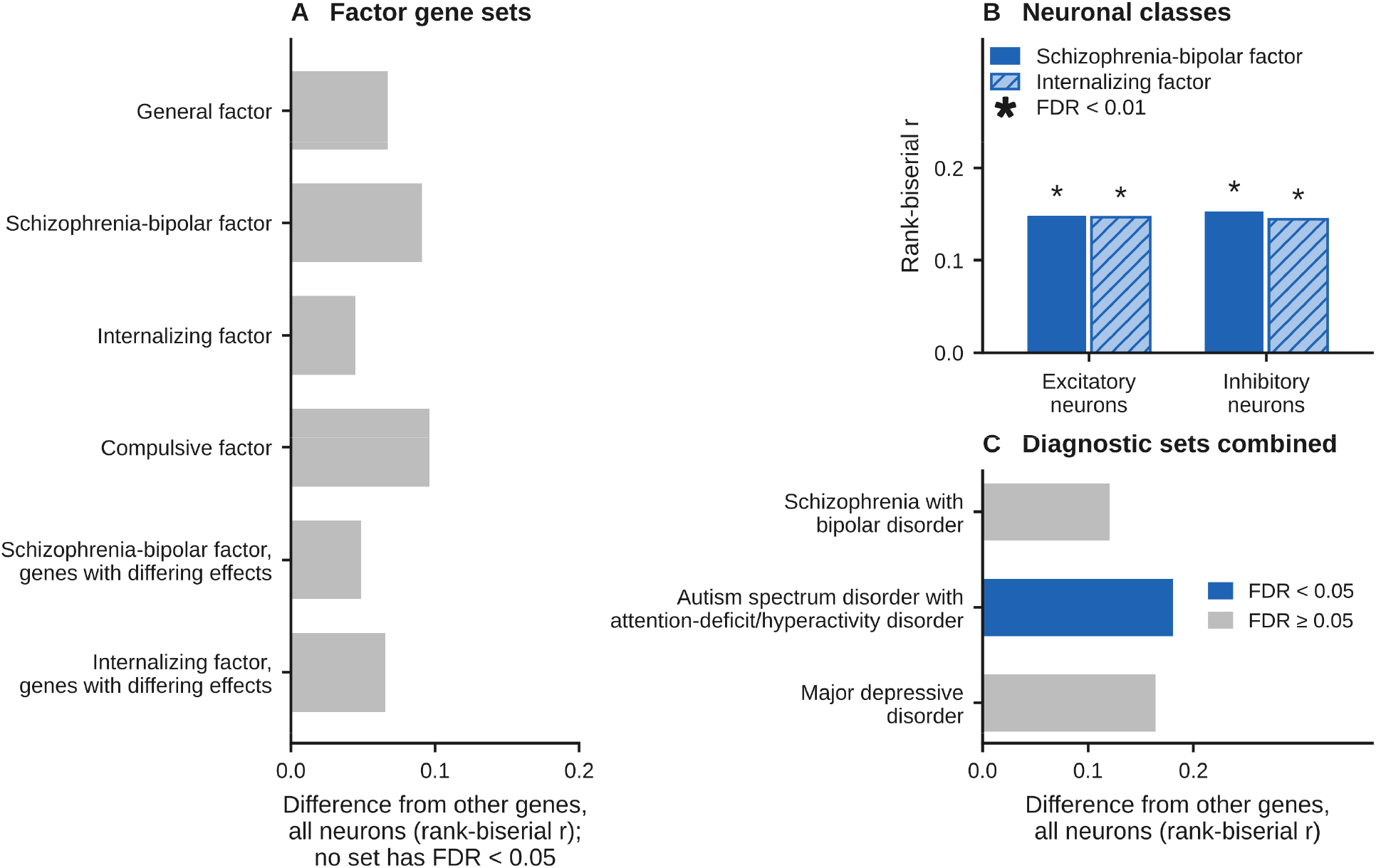
Gene sets grouped by genomic factors of psychiatric risk. (A) Difference in the conservation score between each of six published gene sets [24] and all other genes in all neurons (rank-biserial correlation, r): the sets for the general, schizophrenia-bipolar, internalizing and compulsive factors, and, for the schizophrenia-bipolar and internalizing factors, the genes whose effects differ among the disorders of the factor. No set has a false discovery rate (FDR) below 0.05. The comparison of the genes with differing effects with the genes with shared effects is shown in Fig. S5. (B) The same difference in excitatory and in inhibitory neurons for the schizophrenia-bipolar factor (solid bars) and the internalizing factor (hatched bars); asterisks mark an FDR below 0.01. Both factors are more conserved in excitatory and in inhibitory neurons. (C) Sets built from diagnostic gene sets (exploratory), analyzed because the published gene set for the neurodevelopmental factor is too small to test. Blue, FDR below 0.05.

Two factor sets were more conserved than other genes, and in both cases the difference was present in excitatory and inhibitory neurons. For the schizophrenia-bipolar factor, r was 0.148 in excitatory and 0.153 in inhibitory neurons (FDR = 0.002 for both). For the internalizing factor, r was 0.147 and 0.145 (FDR = 0.005 for both; Fig. 6B). In the analysis of all neurons together, neither factor differed significantly from other genes (Fig. 6A), and sets built by combining diagnostic sets gave results in the same direction (Fig. 6C). The general factor and the compulsive factor were not more conserved than other genes in any analysis, and no factor set remained different from other genes once the five gene properties were included in the model (FDR of 0.29 or higher). The diagnostic schizophrenia set gave a less consistent result: it was more conserved than other genes in excitatory neurons (r = 0.256; FDR = 5.7 × 10⁻³), whereas in inhibitory neurons the difference had the same direction but was not significant after correction for multiple tests (r = 0.171; FDR = 0.069). The result for the diagnostic set should therefore not be interpreted as a difference between excitatory and inhibitory neurons. Further analyses of the factor sets, and the analysis of the 36 genes linked to OCD and CTD by exome sequencing [25], for which the data were inconclusive, are described in Supplementary Notes 3 and 4 (Tables S2 and S3; Figs. S5, S6 and S7).

### No disease gene was shown to have different partners in mouse and human

For the use of mouse models, the genes of greatest concern are disease genes whose partners differ between the species. We screened the disease genes with two criteria: a significant result in the test of the local score (the hypergeometric test of the overlap of partners; FDR below 0.05) together with a conservation score below the median of all genes. The two criteria were fixed before the screen was run on the gene panel; the analysis plan was not registered in a public registry (Methods). Fourteen disease genes met both criteria. For each of them, we then asked whether the gene shares fewer partners between the species than comparison genes with similar connectivity, that is, with a similar number and strength of connections in the network. This was not the case for any of the 14 genes (P = 0.56 to 0.96).

This negative result is the main finding of this part of the study. The 14 genes were selected because a standard test of overlap (the hypergeometric test) was significant for them, and this test assumes that all genes are equally likely to be partners. In a co-expression network this assumption does not hold. Highly connected genes are among the strongest partners of many genes, so the lists of partners of a gene in the two species can share such genes more often than the test assumes, and the test is too liberal. Any study that tests the overlap of partners in correlation networks should therefore make the test against comparison genes of similar connectivity. Apparent partner differences for the 14 genes could also partly reflect the fact that mouse mPFC and human DLPFC are not homologous, and none of them should be given priority for the development of mouse models on the basis of this evidence alone. The 14 genes, the further checks and the tests in networks of protein interactions are given in Supplementary Notes 5 and 6 (Tables S4 and S5; Figs. S8, S9 and S10); the list is provided only for hypothesis generation.

### Effect of sample number, stability of the results and an independent human dataset

The number of samples had a strong effect on the global score (Supplementary Note 7; Figs. S11 and S12; Table S6). In glial cells the median global score was close to zero (0.004 to 0.016 for single glial types), but the glial networks were built from only about 15 pseudobulk samples in mouse and 7 in human for each glial type. When the neuronal data were reduced at random to these sample numbers, their median global score for the 5,000 genes analyzed in glial cells fell from 0.190 to 0.031, so most of the apparent difference between neurons and glial cells is accounted for by sample number. A simulation showed that these sample numbers are sufficient to detect that some conservation is present but not to measure how strong it is. We therefore draw no biological conclusion about the degree of conservation of glial networks or about a difference between glial cells and neurons. In neurons, the score was still rising at the largest sample number, and we estimate that a comparison of co-expression between species for one cell type needs on the order of 50 or more pseudobulk samples per species.

The estimates were stable under alternative analysis choices (Supplementary Note 1; Figs. S13, S14 and S15). When any one of the seven datasets was removed, the median global score stayed between 0.134 and 0.166. The scores of single genes changed little when the networks were built with the Pearson correlation or with mutual rank in place of the Spearman correlation, and the higher conservation of the combined disease genes was found with and without the removal of differences between datasets.

To test whether the human network is specific to the three datasets from which it was built, we used the seven control donors of an independent dataset of human PFC (GSE174367 [27]; 3,450 neurons; Fig. S16). Of five human modules, defined for this test on the 5,000 most variable genes, four were found again: the genes of each were far more strongly co-expressed with one another in the independent dataset than random gene sets of the same size (Z = 20.0 to 31.4); one module was not found again (Z = 1.7). For single genes the agreement was weak (median 0.128), as expected from the analysis of sample number above, so a test of single genes requires an independent dataset with at least about 50 control donors.

## Discussion

We compared the co-expression partners of 13,214 genes between mouse mPFC and human DLPFC and added two reference measurements, comparisons within a species and a correction for measurement noise, that change how the resulting scores should be interpreted (Fig. 2B, C; Table 2). The similarity of partners for single genes is modest (Fig. 2A; Table 2), gene modules are strongly preserved (Fig. 3A), disease genes are no more conserved than matched control genes (Fig. 5B; Table 3), and two analysis choices distort results (the selection of the most variable genes, Fig. S4; small numbers of samples, Fig. S11). Several of these conclusions confirm what earlier studies established, and we do not claim them as new: for example, similar overall structure of human and mouse co-expression networks, together with differences for many single genes and preservation of modules, was shown in bulk tissue and across many species [11, 12, 13, 15]; the major cortical cell classes are conserved while their molecular features differ [7, 9]; and conservation of co-expression was reported to go together with mutational constraint in a comparison of human and mouse brain [4]. Two other earlier findings were not tested here: that glial cells are more divergent than neurons [4] (our glial data are underpowered, as discussed below), and that gene expression in the excitatory neurons of layers 2 and 3 that project within the telencephalon changed exceptionally fast on the human lineage [28]. The lower scores that we found in inhibitory neurons than in excitatory neurons (median global score 0.057 against 0.108; Fig. 2A; Table 2) were not reported in these studies and run in the opposite direction to a comparison of motor cortex, which found fewer conserved marker genes for excitatory than for inhibitory neuron types [9]. The two studies differ in the quantity compared (the co-expression partners of all genes of the panel in our study, the marker genes of cell types in the study of motor cortex) and in the cortical region (prefrontal and primary motor cortex). Within our study, the score also depends on the number of pseudobulk samples (Fig. S11), and the analyses of excitatory and of inhibitory neurons were not matched in this number, so a difference in sample number between the two classes could contribute to the difference in their scores. The difference between excitatory and inhibitory neurons in our data therefore needs replication in other datasets before it is interpreted. Risk genes for neuropsychiatric disorders are known to be under strong mutational constraint and to be concentrated in cortical co-expression modules [18, 22]; that their higher conservation is accounted for by general gene properties (Figs. 4B and 5B) is expected from these observations and from the relation between constraint and conservation of co-expression [4]. The present study extends this work in four ways: it gives reference values from comparisons within a species (Fig. 2B), it corrects for measurement noise (Fig. 2B, C; Table 2), it controls for the number of samples (Fig. S11), and it tests single genes against comparison genes of similar connectivity (Table S4). Its main contribution is a set of reference measurements for interpreting comparisons of networks between species.

The same-species comparisons change how earlier comparisons of co-expression between species should be interpreted. The variation between datasets of one species is similar in size to the difference between species (Fig. 2B; Table 2), so a low score between species cannot be attributed to divergence unless same-species scores are reported with it; we are not aware of an earlier single-cell comparison of networks between species that does so. The regions compared are a second reason for caution. Every score between the species in this study is the score of an mPFC-DLPFC comparison, and the two regions are not homologous: the granular areas of the lateral PFC of primates, which include the DLPFC, do not exist in rodents, and the areas of the rodent mPFC may instead be homologous to cingulate areas of primates [3]. Gene expression differs between areas of the neocortex within a species [29], so a gene could have different partners in mPFC and DLPFC even within one species. Our datasets contain one region per species, so we cannot separate a difference of region from a difference of species, either for single genes or for the median scores. Data from two or more frontal regions in each species are needed to do so.

For disease genes, the primary evidence is the comparison with matched control sets: the seven neuropsychiatric sets combined were no more conserved than control genes of similar constraint, expression level, expression variance, length and GC content (Fig. 5B; Table 3), and the regression model agreed (Fig. 4B). The higher conservation of disease genes is therefore consistent with general gene properties and with the route by which the genes were discovered, which favors genes under strong constraint. Only the intellectual disability set exceeded its control sets, an exploratory signal that was marginal after correction for multiple tests (FDR = 0.054; Fig. 5B; Table 3). For the use of mouse models, the practical conclusion is that high-confidence risk genes for neurodevelopmental disorders are more conserved than the average gene (Fig. 4A); this is consistent with their strong constraint and high expression and does not require a property that is specific to disease genes. Grouping the genes by genomic factor instead of diagnosis showed that the schizophrenia-bipolar factor and the internalizing factor are more conserved than other genes in excitatory and in inhibitory neurons alike before adjustment for gene properties (Fig. 6B; Table 3); after adjustment, which was made in the analysis of all neurons, no factor set differed from other genes (Table 3). For the schizophrenia-bipolar factor, the finding in excitatory and in inhibitory neurons is compatible with the report of the consortium that the genes of this factor are enriched, in data from fetal brain, both in interneurons (which are inhibitory) and in seven subtypes of excitatory neurons (cell-type enrichment analysis in the Results of [24]; in data from adult brain, the same factor was enriched in excitatory neurons of the deep layers). The diagnostic schizophrenia set was significantly more conserved in excitatory neurons only (Table 3), but this result does not show a difference between excitatory and inhibitory neurons, because the difference in inhibitory neurons had the same direction and was close to significance. For claims about mechanism across species, gene sets defined from genetic data are therefore preferable, whereas diagnostic sets remain useful where the clinical category is the question. None of the 14 disease genes selected by the screen showed fewer cross-species partners than connectivity-matched comparison genes (Table S4, Check 4; all P ≥ 0.56). We therefore do not interpret any of them as showing reduced partner sharing between species. Table S4 lists the 14 disease genes with the outcome of four pre-registered checks. The first asked whether each gene also met the two selection criteria when differences between datasets were not removed. The second asked whether selection could be due to differences between species in cell-subtype composition. The third asked whether each gene’s 50 strongest human partners were recovered in an independent human dataset; this check was not evaluable for any of the 14 genes because [state reason, e.g., the independent dataset did not cover these genes at sufficient depth]. The fourth asked whether each gene shares fewer partners between species than comparison genes of similar connectivity, defined as a similar number and strength of connections in the network.

Two analysis choices are of general relevance. Because disease genes vary less in expression than other genes (Fig. S4), any selection of the most variable genes discards most of them (18% were retained among the 5,000 most variable genes) and produces a false negative result; to our knowledge, this effect has not been quantified before in this context. The effect of sample number is related to the known problem of treating cells from one individual as independent observations [30, 31]: with 7 to 15 pseudobulk samples per glial type, the scores fall toward zero, so apparent divergence is expected even when none exists (Fig. S11; Supplementary Note 7). The glial analysis of this study detects that some conservation is present but is underpowered for measuring its strength (Fig. S12; Table S6), and no biological conclusion about the degree of glial conservation can be drawn from these data; in particular, they neither support nor contradict the earlier report that glial cells are more divergent than neurons between human and mouse [4]. The statistical power of a comparison at the level of cell types should be estimated before its result is interpreted, for example with methods developed for the design of single-cell studies of differential expression, such as scPower [32], or with a simulation of the kind used here, and a plateau estimated from a rising curve should be reported together with its dependence on the curve chosen (Fig. S11A).

The Supplement also contains an exploratory analysis of gene expression before and after birth in human cortex, based on the BrainSpan atlas of the developing human brain (Supplementary Note 8; Fig. S17), and an untested proposal for rating how faithfully a mouse model reproduces human gene networks (Supplementary Note 9; Fig. S18). Neither is used for any conclusion of this study.

### What this study does and does not claim

The study provides similarity scores for 13,214 genes in an mPFC-DLPFC comparison, together with same-species reference values and a measure of reproducibility for each gene (Fig. 2; Table 2). The study does not claim that single genes are highly conserved, that disease genes have a network biology of their own, that any gene has been shown to have different partners in the two species, that the glial data show how strongly glial networks are conserved, that differences between the species can be separated from differences between the two regions, or that the relation between disease genes and conservation runs causally through mutational constraint.

Six limitations are most important for these conclusions; the complete list is given in Supplementary Note 10. First, the comparison is between two regions that are not homologous, which is expected to lower the scores and could by itself produce partner differences for single genes. Second, the glial analysis is underpowered for measuring the strength of conservation (Supplementary Note 7). Third, LOEUF, the only measure of constraint used, is estimated with error, so the adjustment for constraint is incomplete, and the small difference that remains in the regression model (coefficient 0.0049; Fig. 4B) cannot be assigned to disease genes specifically. Fourth, the assignment of mouse subtypes to human subtypes agreed between two methods for only 10 of 21 subtypes (Fig. S1); assignments of subtypes, including those of excitatory subtypes, are exploratory, and all primary conclusions are based on broad classes. Fifth, the noise-corrected median (0.349; Fig. 2B; Table 2) is an upper bound, and the independent human dataset, with seven control donors, can test modules but not single genes (Fig. S16). Sixth, the documentation of one input and the public availability of another are incomplete. The origin of the table of orthologs could not be reconstructed, and the analyses were not repeated on the 12,698 genes of the panel that are one-to-one orthologs in a table retrieved from Ensembl (Table S7). The raw data of the in-house mouse dataset will be deposited when this article is published after peer review; until then, the analysis can be repeated from the pseudobulk expression profiles in the analysis package but not from the raw counts of that dataset. Future comparisons of networks between species should report same-species reference values, test claims about gene sets against matched control genes and claims about single genes against comparison genes of similar connectivity, and estimate the number of samples needed in advance. Co-expression does not show which gene regulates which; the next layer of analysis is the comparison of gene-regulatory networks built from single-cell data on RNA and chromatin accessibility [33, 34], preferably measured in the same nuclei [33] (Fig. S19). The scores, the modules and the list of 14 genes selected by the screen are released so that others can use and test them.

## Methods

Details of parameters and of secondary analyses are given in the Supplementary Methods, under the same headings as below.

### Datasets

Mouse mPFC data comprised one in-house dataset, obtained by single-cell RNA sequencing of whole cells, and three public snRNA-seq datasets (Gene Expression Omnibus [GEO] accessions GSE248200, GSE291951 and GSE233763). Human DLPFC data comprised three public snRNA-seq datasets (GSE243292, GSE207334 and GSE261983; PsychENCODE Consortium [35]). An independent human dataset (GSE174367 [27]) was used to test the human network; only the RNA data of its control donors were used. The numbers of cells or nuclei, cell types and pseudobulk samples are given in Table 1. The difference between whole cells and nuclei is one of the differences between datasets. It is contained in the dataset term of the model that removed these differences within each species (see below) and in the same-species comparison of the in-house dataset with the other three mouse datasets (Table 2). The median global score between the species changed little when any one dataset, including the in-house dataset, was removed (Fig. S13B). The mouse glial cells come from the in-house dataset only, so for glial cells this difference cannot be separated from the difference between the species.

The in-house dataset consists of mPFC samples from control mice (Ai9-TRAP2 mice bred on a C57BL/6J background; 20 female and 11 male mice, 3 months of age). The use of these mice was approved by the Institutional Animal Care and Use Committee (IACUC) and the Institutional Biosafety Committee (IBC) of the University of Wyoming. The single-cell RNA sequencing data of these mice were collected for an earlier study, in which the dataset was first reported [36].

The assignment of each cell or nucleus to a cell type was taken from the source study. To relate mouse subtypes to human subtypes, the expression profile of each mouse subtype was correlated with the profile of each human subtype, and each mouse subtype was assigned to the human subtype with the highest correlation. A second assignment, based on shared marker genes, agreed with the first for only 10 of 21 subtypes (6 of 11 inhibitory subtypes, 4 of 9 excitatory subtypes, and not the one subtype that the source study assigned to neither class). Assignments of subtypes, including those of excitatory subtypes, are therefore exploratory and depend on the method. The analyses of broad classes (all neurons, excitatory neurons, inhibitory neurons and glial cells), on which all primary conclusions are based, do not depend on the assignment.

### Pseudobulk samples and removal of differences between datasets

Cells or nuclei from one individual are not independent observations, and treating them as independent overstates the evidence [30, 31, 37]. For each sample of a source dataset and each cell subtype, we therefore summed the raw counts of all cells or nuclei into one expression profile (a pseudobulk sample), provided that at least 10 cells or nuclei were available. Counts were scaled to counts per million (CPM) and transformed as log2(CPM + 1). Differences between datasets were removed within each species with the function removeBatchEffect of the R package limma [38], which fits a linear model with one term per dataset and subtracts the fitted dataset effects. The choice of this method was checked against uncorrected data and against ComBat-seq [39] (Supplementary Methods).

### Genes analyzed and provenance of the list of orthologs

Mouse genes were matched to human genes with a table of one-to-one orthologs, and mouse gene names were converted to the human gene symbols before the networks were built. The primary analysis used the 13,214 one-to-one orthologs that were expressed (log2 CPM above 0) in at least 10% of the pseudobulk samples of each species.

The table of orthologs used for the analysis contains 17,392 gene pairs. Its source, its version and the date of its retrieval were not recorded and could not be reconstructed; the table itself is part of the analysis package, so the gene panel can be reproduced exactly. To document the panel against a source that can be cited, mouse-to-human orthologs were retrieved from Ensembl BioMart (www.ensembl.org; Ensembl release 116, June 2026; *Mus musculus* genes (GRCm39) dataset; attributes: gene name, name of the human ortholog, type of orthology, and percentage of sequence identity; no filters; accessed September 2026), and the pairs classified as one-to-one orthologs were kept (17,067 pairs). The two tables share 15,720 pairs (Jaccard index 0.84; Table S7). We also built a panel from the Ensembl table under the same expression filter; in this panel, a gene with several counterparts in the other species was represented by the counterpart with the highest mean expression. This panel contains 12,880 genes, all of which are in the primary panel (97.5% of its 13,214 genes); 12,698 of them are one-to-one orthologs in the Ensembl table, and 182 have several counterparts. The retrieved Ensembl table is archived in the analysis package. The analyses were not repeated on the genes confirmed by Ensembl, so we cannot state whether every conclusion holds on that subset; we regard this as a limitation.

### Co-expression networks and the scores

For each species and each cell class, we computed the Spearman correlation between every pair of the 13,214 genes across the pseudobulk samples. For each gene, this gives a list of correlations with all other genes, which we call its co-expression profile. The global score of a gene is the Spearman correlation between its co-expression profile in mouse and in human; its significance was assessed by permutation. The local score is the Jaccard index of the gene’s 50 most strongly correlated genes (its 50 strongest partners) in the two species; its significance was assessed with the hypergeometric test, which gives the probability of the observed number of shared partners if the partners were drawn at random from all genes. This test does not take into account that highly connected genes are partners of many genes, and the comparison with genes of similar connectivity showed that it is too liberal (see the screen below). In this study it serves only to select genes for the screen and is not used as evidence about a single gene.

The global score is the score reported for single genes, for the same-species comparisons, for the correction for measurement noise and for the analyses of sample number. For the tests of gene sets in neurons and for the screen for disease genes with different partners, the two scores of each gene were combined into one conservation score: the mean of the local score and of the global score rescaled to the range 0 to 1, that is, (global score + 1)/2. The median conservation score was 0.296 in all neurons.

### Comparisons within a species and correction for measurement noise

For the same-species comparisons, the pseudobulk samples of one dataset formed one set and the samples of the remaining datasets of that species formed the other set. A network was built in each set, and the global score of every gene was computed between the two networks exactly as between the species. This was done once for each dataset, which gave three comparisons in human and four in mouse.

To measure reproducibility, the pseudobulk samples of a species were divided at random into two halves, a network was built in each half, and the global score of each gene was computed between the two halves. This was repeated 20 times, and the mean over the repetitions is the split-half reliability of the gene in that species. The noise-corrected global score of a gene is its observed score divided by the geometric mean of its reliabilities in the two species (the classical correction for attenuation). Because influences shared by the two halves would inflate the correction, we treat the corrected value as an upper bound.

### Gene modules and their preservation

Modules were defined in each species with WGCNA [40], using the pseudobulk samples of all neurons and the 13,214 genes; the settings are given in the Supplementary Methods. Whether a module defined in one species is preserved in the other species was tested in both directions with the module preservation statistics of WGCNA [41]. The summary statistic, Zsummary, combines measures of how densely the genes of the module are connected in the other species and of how similar their pattern of connections is, each expressed relative to random gene sets. A Zsummary above 10 was taken as strong preservation. The overlap between each mouse module and each human module was tested with the hypergeometric test. Because Zsummary increases with the size of a gene set, the 5,267 unassigned mouse genes were compared with 50 random gene sets of the same size, evaluated with the same statistic.

### Gene sets

The seven neuropsychiatric gene sets were: autism spectrum disorder (93 genes; the SFARI Gene database of the Simons Foundation Autism Research Initiative [42]), schizophrenia (63 genes; the SCHEMA exome study [19] and a genome-wide association study [43]), bipolar disorder (47 genes) and attention-deficit/hyperactivity disorder (28 genes), both from genome-wide association studies of the Psychiatric Genomics Consortium (the work of the consortium is reviewed in [44]), major depressive disorder (40 genes; genome-wide association studies [45, 46]), intellectual disability (73 genes; a curated clinical gene panel) and epilepsy (65 genes; a curated clinical gene panel). The neurological gene sets were: Alzheimer’s disease (80 genes from a genome-wide association study [47], and a second set of 174 genes from the Open Targets platform [48]), Parkinson’s disease (102 genes; Open Targets [48]; see also the genome-wide association study [49]) and amyotrophic lateral sclerosis (67 genes; Open Targets [48]; see also the genome-wide association study [50]). The 11 sets together contain 539 genes of the panel. A set had to contain at least 10 genes of the panel to be tested; this rule was set in advance.

The gene sets for the genomic factors were taken from Supplementary Tables 46 and 47 of the cross-disorder analysis [24]; genes identified in fetal brain and in adult brain were combined. The high-confidence gene sets for OCD and CTD were taken from Supplementary Table 16 of the exome study [25]. The identifiers of the Open Targets diseases and the definitions of the factor sets and of the OCD and CTD sets are given in the Supplementary Methods; all external files are stored in the analysis package with their source address, version, date of access and checksum.

### Tests of gene sets

Each set was compared with all other genes of the panel by the Wilcoxon rank-sum test of the conservation scores, and the size of the difference is reported as the rank-biserial correlation (r). The tests were made in all neurons, in excitatory neurons and in inhibitory neurons.

Two analyses took general gene properties into account. The primary analysis used matched control sets. Each gene of a disease set was replaced by a gene that was not in the set and was similar in five properties: LOEUF (a measure introduced in [26]; the values were taken from the Genome Aggregation Database, gnomAD, version 4.1; the database is described in [51]), mean expression, expression variance, log10 gene length and GC content. For each disease set, 1,000 control sets were drawn, the rank-biserial correlation was computed for each, and the P value is the proportion of control sets with a value at least as high as that of the disease set. A disease set is called more conserved than its matched controls if this one-sided P value, corrected for multiple tests, is below 0.05. In the supporting analysis, the conservation score of each gene was regressed on membership in the disease set together with the five gene properties, with standard errors that are robust to unequal variance among genes (HC3). The model used the 12,636 genes for which all five properties were available. LOEUF was the only measure of mutational constraint in the model; other measures (for example the probability of intolerance to loss-of-function variants, or constraint against missense variants) were not tested. Whether the difference between disease genes and other genes depends on constraint was tested by the interaction between membership and LOEUF in the regression model and by a test for trend across tenths of the genes ordered by LOEUF.

### Screen for disease genes with different partners

A disease gene was selected for closer examination if the test of its local score had an FDR below 0.05 and its conservation score was below the median of all genes; we refer to these two criteria as the screen and to the genes that met them as the selected genes. The two criteria and four checks of the selected genes were defined before the analysis (Supplementary Methods). The decisive check asked whether a selected gene shares fewer partners between the species than comparison genes of similar connectivity: its overlap of partners was compared with the overlap of 1,000 comparison genes that were matched to it in the number of connections (degree) and in connection strength, and the P value is the share of comparison genes whose overlap is at most as large as that of the selected gene.

### Effect of sample number, and statistical power at glial sample numbers

Neuronal pseudobulk samples were selected at random to match the sample numbers of one glial type (15 in mouse and 7 in human), of all glial types together (74 and 34), and a series of increasing sample numbers (Supplementary Methods); 50 random selections were made for each. The global score was recomputed for each selection on the genes analyzed in glial cells.

To determine which level of conservation the glial sample numbers can detect, we ran a simulation with the neuronal data. The conservation was lowered stepwise by exchanging the gene labels of a growing fraction of the genes in the human data, and the data of each step were reduced to the glial sample numbers in 200 random selections of samples. A level of conservation counts as detected in a selection when the median global score of the selection exceeds the 95th percentile of the medians obtained by chance (all gene labels exchanged), and the statistical power is the share of selections in which the level is detected. The smallest detectable level is the lowest level with a power of at least 0.80 (Supplementary Methods; Supplementary Note 7).

### Independent human dataset

From GSE174367, control donors with at least 10 excitatory or inhibitory neurons were used (7 donors; 3,450 neurons). Counts were summed per donor, transformed as log2(CPM + 1), and used to build one Spearman network of the 13,214 genes. For single genes, agreement was measured as the Spearman correlation between the gene’s co-expression profile in the main human data and in the independent dataset. For modules, the mean correlation among the genes of a module in the independent network was compared with that of 50 random gene sets of the same size and expressed as a Z score. This test was applied to the five human modules defined on the 5,000 most variable genes (1,060, 779, 459, 159 and 96 genes).

### Statistics and software

Tests were two-sided, except for the comparisons with matched control sets and the tests in the protein-interaction networks, which ask whether an observed value is higher than in comparison sets and are therefore one-sided. The false discovery rate (FDR) was controlled by the Benjamini-Hochberg method within families of tests that were defined before the analysis, and a result was called significant at an FDR below 0.05; all families, with the tests in each, are listed in Table S8. The rules that are described in this article as defined or set in advance (the criteria of the screen and the four checks, the minimum number of genes in a tested set, the families of tests, the three tests in networks of protein interactions and the rules for calling a test uninformative) were fixed in the analysis plan before the corresponding results were computed. The analysis plan was not registered in a public registry. We give more weight to the size of a difference than to its P value, because with 13,214 genes even very small differences reach nominal significance. All random procedures were started from a fixed value (seed 42). A repetition of the complete analysis from the analysis package is described in Supplementary Note 11. The analyses were run in R 4.4.3 (packages WGCNA 1.74 and limma) and Python 3 (numpy, pandas, scipy, statsmodels, anndata and networkx). The complete software environments are provided with the analysis package.

## Declarations

### Ethics statement

All data used in this study come from publicly available datasets, except one in-house single-cell RNA sequencing dataset that was collected for an earlier study [36]. The animal procedures of the in-house dataset were approved by the Institutional Animal Care and Use Committee (IACUC) of the University of Wyoming under protocol number 2023-0056. The human data are public datasets from the cited studies; no new human samples were collected for this study.

### Author contributions

Q.Q. S designed the study, performed the network analyses, and wrote the manuscript. T.C. and CZ help set-up the public access data, curated the data and performed quality checks, and performed statistical analysis and provided inputs to the manuscript.

### Funding

This work was supported by grants R21MH131363, R21MH141703 and P20GM121310.

### Competing interests

The author declares no competing interests.

## Acknowledgements

We thank the investigators who made the public datasets available, the PsychENCODE Consortium [35], and Yihan Wang, who collected the mouse single-cell RNA sequencing data reported in an earlier study [36].

## Supplementary information

### Supplementary Methods

The headings follow those of the Methods. Each paragraph gives the parameters and the secondary analyses that the Methods refer to.

#### Datasets

Together the four mouse datasets contain 107,471 neurons, assigned to 21 subtypes, and 14,024 glial cells of five types; the glial cells come from the in-house dataset. The three human datasets contain 93,433 neurons, assigned to 18 subtypes, and 31,283 glial cells of five types; the glial cells come from the control donors of GSE243292. In the independent human dataset (GSE174367 [27]), RNA and chromatin accessibility were measured in separate nuclei from the same prefrontal cortex tissue samples of donors with Alzheimer’s disease and of control donors; only the RNA data of the control donors were used.

#### Pseudobulk samples and removal of differences between datasets

The summing of counts gave 180 neuronal and 74 glial pseudobulk samples in mouse and 320 neuronal and 34 glial pseudobulk samples in human. The linear model that removed the differences between datasets contained the dataset as its only term. A model that also contained the cell subtype could not be fitted in mouse, because 13 of the 21 mouse subtypes occur in a single dataset, so that subtype and dataset cannot be separated. To check the choice of method, we compared the corrected data with uncorrected data and with data corrected by ComBat-seq [39], a method that works on counts. The check used the first three principal components of the pseudobulk samples, computed separately for each species and for each version of the data, and measured, for each component, the proportion of its variance that is explained by the dataset (η²). Before correction these proportions were 0.82, 0.18 and 0.004 in human and 0.75, 0.81 and 0.86 in mouse. After the limma correction they were 0 in both species. After ComBat-seq they were 0.03, 0.005 and 0.80 in human and 0.31, 0.63 and 0.11 in mouse. We therefore used the limma correction for all network analyses.

#### Genes analyzed

A second panel, the 5,000 genes with the highest mean variance across pseudobulk samples, was used for sensitivity analyses and to demonstrate the effect of selecting variable genes.

#### Co-expression networks and the scores

The significance of the global score was assessed with 200 permutations per gene, and with 2,000 permutations in a check of resolution. To test whether the scores depend on the measure of association, the networks were rebuilt with the Pearson correlation, with the mutual rank of the Spearman correlations, and with proportionality (φ [52]). Proportionality was computed on centered log-ratio values, which are appropriate for relative abundances, and not on log2(CPM + 1) values (CPM, counts per million).

#### Chance values for the scores

To obtain the global score expected by chance, the co-expression profile of each gene in mouse was correlated with the profile of a randomly chosen gene in human (1,000 randomizations). Observed and chance values were compared within tenths of the genes ordered by connection strength (the sum of a gene’s absolute correlations with all other genes). To test whether the higher conservation of the combined disease genes could arise from the size of the gene set alone, the test was repeated on 1,000 random gene sets of the same size.

#### Comparisons within a species and correction for measurement noise

Besides the median of the corrected scores of single genes, we report, for comparison, the observed median divided by the geometric mean of the two median reliabilities.

#### Gene modules and their preservation

The module analysis has three steps. First, a correlation that is insensitive to outlying samples (the biweight midcorrelation) is computed for every pair of genes and raised to a power, which emphasizes strong correlations; the sign of the correlation is kept, so that only positively correlated genes are grouped. The power was chosen as the lowest value at which the network approximates the form typical of biological networks, with few highly connected genes and many weakly connected ones (scale-free fit); it was 7 in mouse and 11 in human. Second, genes are grouped by hierarchical clustering. Third, groups of at least 30 genes are taken as modules, and modules with similar expression patterns are merged (merging height 0.25). Genes that fall into no module are left unassigned. The preservation statistics used 200 permutations. For the overlap of modules, the hypergeometric test gives the probability of at least the observed number of shared genes among the 13,214 genes if the two modules were unrelated; the P values were adjusted by the Benjamini-Hochberg method, which controls the false discovery rate (FDR). The 50 random gene sets that were compared with the unassigned genes were each evaluated with 30 permutations.

#### Gene sets

From the Open Targets platform, genes with an association score of at least 0.5 were included; the disease identifiers were MONDO_0004975 (Alzheimer’s disease), MONDO_0005180 (Parkinson’s disease) and MONDO_0004976 (amyotrophic lateral sclerosis). A set for Huntington’s disease (MONDO_0007739) contained only two genes and was excluded. From the cross-disorder analysis [24] we took the genes targeted by the general factor, by the compulsive, schizophrenia-bipolar and internalizing factors, and, for the last two factors, the genes whose effects differ among the disorders of the factor. Sets with fewer than 10 genes in the panel (the neurodevelopmental factor, 2 genes; the substance-use factor, 7 genes) were not tested. The high-confidence gene sets for obsessive-compulsive disorder (OCD) and chronic tic disorders (CTD) were defined with the significance threshold of the exome study [25] (a false discovery rate, q, below 0.1 in its transmission and de novo association test): 12 genes for OCD, 10 for CTD and 34 for the two disorders combined. The three sets together contain 36 genes, of which 34 are in the panel (*CARMIL2* and *CCDC168* are not). In a sensitivity analysis, four genes in which damaging variants were more frequent in control individuals than in patients (*ABCD2*, *DNAH9*, *DNAH10* and *TEP1*) were removed. Sets built by combining diagnostic sets (schizophrenia with bipolar disorder; autism spectrum disorder with attention-deficit/hyperactivity disorder; major depressive disorder alone) were analyzed as an exploratory family, because the published gene set for the neurodevelopmental factor contains only 2 genes of the panel.

### Tests of gene sets: matched control sets

The replacement for a disease gene was drawn at random from the genes in the same tenth of LOEUF (loss-of-function observed/expected upper bound fraction) that lay within 0.2 standard deviations of the disease gene in each property (log LOEUF, mean expression, expression variance, log10 gene length and GC content, the proportion of guanine and cytosine bases; 0.1 standard deviations in a sensitivity analysis). If no gene met this condition, the replacement was drawn at random from the five most similar genes of the same tenth. The P value was computed so that it cannot be zero (the method of Phipson and Smyth). After matching, the disease genes and their controls differed by about 0.04 standard deviations in each property. Gene length and GC content were taken from Ensembl BioMart. The test for trend across tenths of LOEUF used Kendall’s τ with a permutation P value.

### Genes selected by the screen: the four checks

In the first check, the gene had to meet the two criteria of the screen both in the data from which differences between datasets had been removed and in the uncorrected data. In the second check, the gene had to meet the two criteria when the contributions of the cell subtypes were balanced between the species. In the third check, the 50 strongest human partners of the gene had to be found again in the control donors of GSE174367; because that dataset has seven donors, the check was recorded as not evaluable for all genes. In the fourth check, the overlap of partners of the selected gene was compared with the overlap of 1,000 comparison genes that were matched to it in the number of connections (degree) and in connection strength; the P value is the share of comparison genes whose overlap is at most as large as that of the selected gene. For each of the 14 selected genes, the partners found in human only and the partners found in mouse only were each tested for enrichment of functional categories (Gene Ontology biological process, the Kyoto Encyclopedia of Genes and Genomes and Reactome) with g:Profiler [53].

#### Effect of sample number

The series of increasing sample numbers consisted of seven pairs (7, 10, 15, 25, 34, 50 and 320 pseudobulk samples in human, with 15, 20, 30, 50, 74, 100 and 180 in mouse). In this series the mouse number is about twice the human number, except in the last pair, which is the complete neuronal data. Three curves that approach a plateau (Michaelis-Menten, exponential and Hill) were fitted to the series by non-linear least squares, with confidence intervals from 1,000 resamplings of the residuals. Because the score was still rising at the largest sample number, the plateaus are extrapolations that depend on the curve. The difference between glial cells and neurons at equal sample numbers is the observed glial score minus the neuronal score at the glial sample number, with the standard deviation across the random selections.

#### Statistical power at glial sample numbers

To determine which level of conservation the glial sample numbers can detect, we ran a simulation with the neuronal data, restricted to the genes analyzed in glial cells. The conservation was lowered stepwise by exchanging the gene labels of a growing fraction of the genes in the human data (0%, 25%, 50%, 62.5%, 75% and 87.5%); an exchange of all labels gave the values expected by chance. For each step, the median global score at full sample number defines the level of conservation of that step. The data of each step were then reduced to the glial sample numbers in 200 random selections of samples. A level counts as detected in a selection when the median global score of the selection exceeds the 95th percentile of the chance medians, and the statistical power is the share of selections in which the level is detected. The smallest detectable level is the lowest level with a power of at least 0.80, obtained by linear interpolation between the steps. The simulation was repeated with selections from excitatory neurons only, and the glial data themselves were compared with 200 permutations of their gene labels.

#### The internalizing factor in glial cells

The difference between the genes of the internalizing factor and other genes in the glial networks (global score) was compared with the differences obtained for the same gene set in neuronal data reduced to the glial sample numbers. We decided in advance to call this test uninformative if the smallest difference detectable with 80% power was larger than the observed difference.

#### Independent human dataset

Of the 13,214 genes of the panel, 13,213 were detected in the independent dataset.

#### Further analyses of stability

The scores were recomputed with the alternative measures of association, with panels of 5,000 and 10,000 genes, on data without removal of dataset differences, and after removal of each of the seven datasets in turn (with the correction for dataset differences fitted again each time). To check the assignment of mouse subtypes to human subtypes, a second assignment was built from the overlap of the 30 genes most specific to each subtype in each species (Jaccard index) and compared with the assignment based on correlation. The global score was also correlated with the identity of the protein sequences of the two orthologs, and the difference in expression level between the species (absolute log2 fold change) was correlated with the scores in each cell class.

#### Networks of physical protein interactions

Two resources were used. PCNet version 1.4 [54], obtained from the Network Data Exchange (identifier c3554b4e-8c81-11ed-a157-005056ae23aa), combines many kinds of evidence, including co-expression (18,630 genes and 2,687,393 connections). Because co-expression is also the kind of data from which the genes were selected, a result in PCNet could be circular. The second resource, BioGRID version 5.0.262, was therefore restricted to physical interactions (1,041,378 connections in human; 76,422 in mouse, of which 62,621 could be transferred to human gene names through one-to-one orthologs). Three tests, defined in advance, were applied to the 14 selected genes. The first compared their mean number of connections in PCNet with that of 1,000 sets of comparison genes matched in co-expression connectivity, and with comparison sets drawn from disease genes. The second compared, for each selected gene, the overlap between its physical partners in human and in mouse (Jaccard index in BioGRID) with that of matched comparison genes. The third measured the fraction of the co-expression partners that a selected gene has in human but not in mouse that are also its neighbors in the interaction network, in PCNet and in BioGRID, against matched comparison genes. P values are the proportion of 1,000 comparison sets with a value at least as high as the observed one.

#### Expression before and after birth

Gene expression in the developing human brain was taken from BrainSpan (gene-level data, Gencode version 10; 47,807 genes and 524 samples). The analysis was restricted to cortical samples (288 samples from 41 donors; 123 samples before birth and 165 after birth); 94.3% of the panel genes were present. For each gene we computed the mean expression after birth minus the mean expression before birth, on the log2 scale (log2[RPKM + 1]; RPKM, reads per kilobase per million reads). A negative value means higher expression before birth. Gene sets were compared with other genes on this difference by the rank-based test. Across genes, the difference was correlated with the conservation score (Spearman correlation). Two further analyses were made. The difference was added to the regression model with the five gene properties, and the correlation was repeated within the genes with higher expression in neurons than in glial cells. These were defined as the genes for which the mean expression in the human neuronal samples minus the mean expression in the human glial samples was above the median of all genes. No donor contributes samples from both periods, so an analysis within donors was not possible.

#### Statistics

For tests based on randomization, the smallest attainable P value is limited by the number of randomizations. The fixed starting value of the random procedures (seed 42) is recorded in each output file. The analysis was run a second time from the analysis package, independently of the first run; because the second run used the same data and the same code, it is a check of reproducibility and not an independent replication (Supplementary Note 11).

### Supplementary Note 1. Chance values, stability of the scores, and the assignment of subtypes

#### Chance values

To confirm that the modest median global score is above the value expected by chance, we compared each gene’s correlations in mouse with the correlations of a different, randomly chosen gene in human. Observed and chance values were compared within tenths of the genes ordered by connection strength, which is the sum of a gene’s absolute correlations with all other genes (Fig. S2B). In every tenth, the observed median exceeded the median of the chance values, and in the four tenths with the strongest connections it also exceeded the 90th percentile of the chance values. For the tenth of genes with the weakest connections, the observed median was 0.091 and the chance value 0.012; for the tenth with the strongest connections, the values were 0.385 and 0.030.

#### Measures of association

The networks were rebuilt with the Pearson correlation and with the mutual rank of the Spearman correlations. The scores of single genes from these networks correlated with the primary scores at 0.87 and 0.99 (Fig. S13A; Fig. S2A). A third measure, proportionality, which is designed for relative abundances, gave scores that were unrelated to the primary scores (correlation about −0.04). It measures a different quantity, and we report it separately.

#### List of orthologs

The list of one-to-one orthologs used for the analysis and a list newly retrieved from Ensembl BioMart largely overlapped (Jaccard index 0.84; Table S7; Fig. S14). Admitting genes with more than one counterpart in the other species added only 182 genes. The source of a list of orthologs is therefore more important than the rule for genes with several counterparts. The origin of the list used for the analysis could not be reconstructed, which we report because it limits the reproducibility of the analysis.

#### Removal of differences between datasets

The linear correction that we used (limma) left no detectable association between the first principal components and the dataset, whereas an alternative method that works on counts (ComBat-seq) left such associations (Supplementary Methods). The samples are shown on the first two principal components in Fig. S15 and, for the limma procedure, in Fig. S13C. In human, the association that remained after ComBat-seq is on the third component, which is not shown. The higher conservation of the combined disease genes was found with and without the correction.

#### Removal of single datasets

When each of the seven datasets was removed in turn, the median global score was between 0.134 and 0.166 (Fig. S13B).

#### Assignment of subtypes

The assignment of each mouse neuronal subtype to a human subtype, which we derived from the correlation of expression, agreed with an assignment based on shared marker genes for only 10 of 21 subtypes (6 of 11 inhibitory subtypes and 4 of 9 excitatory subtypes; the one remaining subtype, which the source study assigned to neither class, did not agree; Fig. S1). Assignments of subtypes, including those of excitatory subtypes, are therefore exploratory and depend on the method. All primary conclusions of this study are based on the broad classes (all neurons, excitatory neurons and inhibitory neurons), which do not depend on these assignments.

### Supplementary Note 2. The disease gene sets one by one, and the neurological sets

Table S1 gives the conservation of each of the 11 disease gene sets before and after adjustment for the five gene properties, and Fig. S3 shows the adjusted coefficients and the expression of each set in excitatory and in inhibitory neurons. After adjustment, no single set was significant after correction for multiple tests; the smallest false discovery rate was 0.23, for intellectual disability. Taken one by one, 7 of the 11 sets had a significantly lower expression variance than other genes (Fig. S4), which is the reason why a selection of the most variable genes removes most disease genes.

The four neurological sets (two for Alzheimer’s disease, one for Parkinson’s disease and one for amyotrophic lateral sclerosis) were not more conserved than other genes, but two caveats limit what this result means. Many genes of the Alzheimer’s disease sets are expressed mainly in microglia, whereas the networks tested here are neuronal. Parkinson’s disease primarily affects the substantia nigra, and amyotrophic lateral sclerosis affects the motor cortex and spinal cord, so the prefrontal cortex may not be the relevant region for these sets. The results for the neurological sets are therefore uninformative for these diseases; they are not evidence that conservation is absent.

### Supplementary Note 3. Further analyses of the gene sets for genomic factors

#### Genes whose effects differ among the disorders of a factor

For the schizophrenia-bipolar factor and the internalizing factor, the cross-disorder analysis [24] also lists genes whose effects differ among the disorders of the factor, in contrast to genes whose effects are shared by these disorders. We asked whether the two kinds of genes differ in conservation. They did not (Fig. S5). The sets of genes with differing effects are small (34 and 76 genes; Table S2). We had decided in advance that a result that is not significant in sets of this size would be reported as not distinguishing between no difference and a difference too small to detect.

#### Sets built from diagnostic gene sets

The published gene set for the neurodevelopmental factor contains only 2 genes of our panel and could not be tested. As an exploratory analysis we therefore combined diagnostic gene sets to approximate three factors: schizophrenia with bipolar disorder for the schizophrenia-bipolar factor, autism spectrum disorder with attention-deficit/hyperactivity disorder for the neurodevelopmental factor, and major depressive disorder for the internalizing factor. Where a published factor set exists, the combined diagnostic set gave results in the same direction (Table S2; Fig. 6C). In all neurons, only the set built from autism spectrum disorder and attention-deficit/hyperactivity disorder was more conserved than other genes (r = 0.181; false discovery rate 0.010). We did not build a combined diagnostic set for the compulsive factor: none of the 11 diagnostic sets belongs to a disorder of this factor, and the published gene set for the factor could be tested directly.

The values of all factor sets in each neuronal class are shown in Fig. S6A.

### Supplementary Note 4. Details of the analysis of the gene sets for obsessive-compulsive disorder (OCD) and chronic tic disorders (CTD)

The 36 genes linked to OCD and CTD by exome sequencing [25] are a set discovered entirely through rare variants. Because these genes were identified by tests that are sensitive to mutational constraint, higher conservation was expected, and we treated the set as a further example and not as an independent confirmation. The main set, the genes linked to the two disorders combined, was more conserved than other genes in all neurons before correction for multiple tests (r = 0.224; P = 0.028), but the difference was not significant after correction (false discovery rate 0.266) and was absent within excitatory and inhibitory neurons (Table S3, part B; Fig. S7). The data are therefore inconclusive for this set. The study that identified these genes reported increased expression of the risk genes in postnatal cerebellum and in prenatal and postnatal cortex and striatum [25], so the adult prefrontal cortex may not be the only informative region; a comparison of mouse and human cerebellum or striatum was not made in this study.

The exome study [25] provides three high-confidence gene sets: 12 genes for OCD, 10 genes for CTD and 34 genes for the two disorders combined (Table S3). We designated the set for the two disorders combined, tested in all neurons, as the main test. The difference from other genes before correction for multiple tests (r = 0.224; P = 0.028) was similar when four genes were removed in which damaging variants were more frequent in control individuals than in patients (r = 0.217; P = 0.040). The separate sets for OCD and for CTD contain 11 and 10 genes of the panel, and neither differed from other genes.

To ask how much of the difference is accounted for by the five gene properties, we divided the coefficient of the gene set in the adjusted model by its coefficient in the unadjusted model. A ratio of 1 means that the properties account for none of the difference, and a ratio of 0 that they account for all of it. The ratio was 0.90, with a 95% confidence interval from 0.30 to 1.55 (Fig. S7B). The data are therefore compatible with a large reduction of the difference and with no reduction, and the result can only generate hypotheses. Twelve of the 34 genes in the panel belong to the tenth of all genes with the lowest values of LOEUF, the measure of tolerance of loss-of-function mutations (the distribution of the 34 genes is shown in Fig. S7A). This excess is expected and is not independent evidence, because the statistical tests that identified the genes favor genes under strong constraint.

### Supplementary Note 5. The genes selected by the screen: the four checks, the genes that passed two checks, and *FOXP2*

Fourteen disease genes met the two criteria of the screen: a significant result in the test of the local score (false discovery rate, FDR, below 0.05) and a conservation score below the median of all genes. We refer to them as the selected genes. In the whole panel, 156 genes (1.2%) met the two criteria in all neurons, 91 (0.7%) in excitatory neurons and 55 (0.4%) in inhibitory neurons. Table S4 lists the 14 disease genes among them with the outcome of four checks that had been defined in advance. The first check asked whether the gene also met the two criteria when differences between datasets were not removed. The second asked whether the selection could be due to differences between the species in the composition of cell subtypes. The third asked whether the gene’s 50 strongest human partners were found again in an independent human dataset. The fourth asked whether the gene shares fewer partners between the species than comparison genes with similar connectivity, that is, with a similar number and strength of connections in the network.

Five genes (*BCL11A*, *FOXP2*, *MCTP1*, *NEGR1* and *SRCAP*) passed the first two checks (Fig. S8). The third check could not be evaluated: the independent dataset contains only seven control donors, too few to reproduce the 50 strongest partners of a single gene, so we do not count it as a failure. No gene passed the fourth check (P = 0.56 to 0.96 for the 14 genes).

*FOXP2* is of interest on descriptive grounds only. It is among the most highly connected genes in the human network (96th percentile), and the partners it has in human but not in mouse (Fig. S8) are enriched for genes of system development (the most significant category; Fig. S9), nervous system development and trans-synaptic signaling. It did not pass the fourth check, and its partner differences, like those of the other selected genes, could reflect the fact that mouse medial prefrontal cortex and human dorsolateral prefrontal cortex are not homologous, and not a difference between the species.

Two further observations support the caution. Disease genes were not significantly over-represented among all genes that met the two criteria of the screen (odds ratio 2.4; FDR = 0.22). In addition, when the analysis was restricted to the 5,000 most variable genes, eight different disease genes met the criteria (among them *SATB2*, *CNTNAP2* and *NRGN*), and none of them met the criteria again in the analysis of all 13,214 genes. The 14 selected genes also did not differ from comparison genes in networks of physical interactions between proteins (Supplementary Note 6). The list is provided only for hypothesis generation, and none of the genes should be given priority for the development of mouse models on the basis of this evidence alone.

### Supplementary Note 6. The selected genes in networks of protein interactions

Co-expression is a correlation and does not show that two gene products interact. We therefore asked whether the 14 selected genes also differ from comparison genes in networks of interactions between proteins (Table S5; Fig. S10). Two resources were used: PCNet [54], which combines many kinds of evidence including co-expression, and BioGRID restricted to physical interactions, which contains no co-expression evidence and is the more suitable control. Three tests were defined in advance. First, the selected genes did not have more connections in PCNet than comparison genes matched in co-expression connectivity (P = 0.88). Second, the physical partners of the selected genes in human and in mouse did not overlap less than those of matched comparison genes (P = 0.51). Third, we asked whether the co-expression partners that a selected gene has in human but not in mouse are also its neighbors in the interaction networks. In PCNet the fraction was higher than for comparison genes before correction for multiple tests (P = 0.020; false discovery rate 0.10), but in BioGRID it was not (P = 0.74). This result in PCNet probably arises from the connections in PCNet that are derived from co-expression, which is the same kind of data from which the genes were selected. The networks of protein interactions therefore add no support for any selected gene. A negative result in a database of physical interactions that covers few gene pairs is, however, weak evidence that no interaction exists.

### Supplementary Note 7. Sample number and the underpowered glial comparison: neuronal data at glial sample numbers, the score at increasing sample numbers, statistical power, the difference from neurons at equal sample numbers, and the internalizing factor

#### Low scores in glial cells and the number of samples

In glial cells the median global score was close to zero (0.004 to 0.016 for single glial types). This could be misinterpreted as evidence that the gene networks of glial cells have diverged between mouse and human. The glial networks, however, were built from far fewer pseudobulk samples than the neuronal networks (per glial type, about 15 in mouse and 7 in human). With so few samples the score is strongly lowered and imprecise, so the glial analysis cannot measure how strong the conservation is (for this purpose it is underpowered), and we use the glial data only to show what the number of samples does to the score. We reduced the neuronal data to the same numbers of pseudobulk samples by random selection and recomputed the score for the 5,000 genes analyzed in glial cells (Fig. S11). With all neuronal samples, the median global score of these genes was 0.190. It fell to 0.031 ± 0.009 (mean ± standard deviation over the random selections) when the neuronal data were reduced to the sample numbers of one glial type, and to 0.097 ± 0.013 at the sample numbers of all glial types together (74 in mouse and 34 in human; Fig. S11B). Against glial values of 0.008 for one glial type and 0.064 for all glial types together, sample number alone therefore explains 87% and 74%, respectively, of the apparent difference between neurons and glial cells.

#### The score at increasing sample numbers

When the number of human neuronal samples was increased stepwise from 7 to 320, with the number of mouse samples increased in parallel from 15 to 180, the score rose throughout and was still rising at the largest number (Fig. S11A). Three mathematical curves that level off gave different estimates of the final plateau (0.18 to 0.22), so we report the curve itself and not a single plateau. The practical guidance is that a comparison of co-expression between species for one cell type needs on the order of 50 or more pseudobulk samples per species, and that a score near zero from fewer samples should not be interpreted as biological divergence.

#### What the glial sample numbers can detect

The glial networks are built from 6 or 7 human and 14 to 16 mouse pseudobulk samples for each glial type, and on 34 human and 74 mouse samples for all glial types together. We asked by simulation which level of conservation such sample numbers can detect (Methods and Supplementary Methods; Table S6; Fig. S12). The simulation used the neuronal data, restricted to the 5,000 genes analyzed in glial cells. The conservation was lowered in steps by exchanging the gene labels of 0%, 25%, 50%, 62.5%, 75% and 87.5% of the genes in the human data, which gave median global scores of 0.190, 0.107, 0.058, 0.035, 0.018 and 0.010 at full sample number; we call these values the levels of conservation. An exchange of all gene labels gave the chance values. With 7 human and 15 mouse samples, the statistical power to distinguish a level from chance was 0.79 at a level of 0.018 and 0.93 at a level of 0.035, and the smallest level detected with a power of 0.80 was 0.019, one tenth of the neuronal value. With 34 human and 74 mouse samples, the power was 0.99 at the lowest level tested (0.010). When the samples were selected from excitatory neurons only, which resembles a single glial type more closely than a selection from all neuronal subtypes, the smallest detectable levels were similar (0.020 and 0.008).

The glial data were then compared with chance values of their own, obtained by permutation of the gene labels in the human glial network (Table S6, part B). In each of the five glial types and in all glial types together, the observed median global score exceeded every one of the 200 chance values (P = 0.005, the smallest value attainable with 200 permutations). Conservation is therefore detectable in the glial networks.

The strength of this conservation cannot be measured at these sample numbers. In the simulation, conservation of the size found in neurons (level 0.190) gave scores between 0.011 and 0.059 with 7 human and 15 mouse samples (the range that contains 95% of the selections; mean 0.031), and conservation two to five times weaker (levels 0.107 to 0.035) gave scores between 0.001 and 0.037. The glial values of the single types lie below the first range or in its lower part, and all lie inside the second. With 34 human and 74 mouse samples, conservation as in neurons gave scores between 0.074 and 0.124, and the value of all glial types together lies below this range (0.064, or 0.073 when recomputed; Table S6); it lies inside the range obtained with excitatory neurons only (0.050 to 0.083). The glial values are therefore compatible with conservation as strong as in neurons and with conservation several times weaker. The comparison is underpowered for the question of how strongly glial networks are conserved, and no biological conclusion about the degree of glial conservation, or about a difference between glial cells and neurons, can be drawn from these data. Two properties of the simulation limit it further: a selection of neuronal samples mixes many subtypes, whereas a glial type is one cell type, and the exchange of gene labels is only one of many ways in which conservation could be lower.

#### Difference between glial cells and neurons at equal sample numbers

The glial value was lower than the value of neuronal data reduced to the same sample numbers by 0.023 for one glial type and by 0.033 for all glial types together (Fig. S11B). We do not interpret this difference, for the reasons given above and because the proportions of glial subtypes differ between the species.

#### The internalizing factor in glial cells

The cross-disorder analysis reported that, in data from adult brain, the genes of the internalizing factor are enriched in oligodendrocytes and in Bergmann glia, a glial cell type of the cerebellum (cell-type enrichment analysis in the Results of [24]). In our glial networks, the genes of the internalizing factor had a higher global score than other genes (r = 0.123). To judge this value, we compared it with the values obtained for the same gene set in neuronal data reduced to the glial sample numbers. The glial value did not exceed these values (z = 0.67; P = 0.31; Fig. S6B). With the present number of glial samples, the smallest difference that could be detected with 80% power is r = 0.185, which is larger than the observed value. By the rule set in advance, the test is therefore uninformative and is not evidence against a difference in glial cells. The smallest detectable difference of a gene set depends on the number of genes in the set (78 genes of the internalizing factor are among the genes analyzed in glial cells) and is not improved by the simulation above, which concerns the median of all genes.

### Supplementary Note 8. Expression before and after birth

As a first analysis of development, we asked whether the gene sets studied here are expressed more strongly before or after birth in human cortex, using the BrainSpan data (288 cortical samples from 41 donors; Supplementary Methods; Fig. S17). For each gene, the quantity analyzed is the mean expression after birth minus the mean expression before birth, so a negative value means higher expression before birth. All gene sets tested tended toward higher expression before birth than other genes (Fig. S17A). The tendency was largest for the compulsive factor (r = −0.19; false discovery rate [FDR] 0.099, which is marginal) and close to zero for the schizophrenia-bipolar factor; no set had an FDR below 0.05. Across all genes of the panel, the difference (after birth minus before birth) was weakly and positively correlated with the conservation score (Spearman correlation 0.020; P = 0.027; Fig. S17B). In the regression model with the five gene properties, the coefficient of this difference had an FDR of 0.075. The correlation was stronger among the genes with higher expression in neurons than in glial cells (0.080; P = 1.7 × 10⁻⁴). Genes that are expressed more strongly after birth therefore tend to be slightly more conserved between mouse and human. The correlation is weak, and we do not interpret it further. Two cautions are needed in interpreting this result. The cortical samples of BrainSpan come from several regions, and the cell composition of cortex changes during development, so the difference between the two periods also reflects cell composition. In addition, the cerebellum could not be examined, because no suitable pair of mouse and human cerebellar snRNA-seq datasets was available.

### Supplementary Note 9. A proposal for rating how faithfully a mouse model reproduces human gene networks

This note outlines a proposal. It is not a result of this study, and it has not been tested. We suggest that the fidelity of a mouse model could be rated from five components (Fig. S18). The first is the similarity of the transcriptome, measured with scores of co-expression conservation of the kind developed here. The second is the similarity of cell-type composition. The third is the agreement of disease genetics, with disease genes grouped by genomic factor [24] and compared with matched control gene sets. The fourth is a search for genes with changed partners, in which each gene selected by a screen is checked against comparison genes of similar connectivity and in networks of protein interactions. The fifth is a single summary rating that weights the first four; the idea of one rating of fidelity for each model follows the classification scores used to rate how closely cancer models resemble tumors [55]. In this proposal the factor gene sets serve as examples of how the third component would be applied and not as evidence for it. Each component can be measured separately today. The weights and thresholds of the summary rating, however, would have to be set with human datasets that were not used to build it and with data on the response to gene perturbation, and such data do not yet exist for the PFC. We present the outline to organize future work, and we caution against using any single component, including the scores of this study, as a measure of the validity of a mouse model.

### Supplementary Note 10. Complete list of limitations

Abbreviations used in this note: PFC, prefrontal cortex; mPFC, medial PFC; DLPFC, dorsolateral PFC; snRNA-seq, single-nucleus RNA sequencing; LOEUF, loss-of-function observed/expected upper bound fraction; OCD, obsessive-compulsive disorder; CTD, chronic tic disorders.

1. *Regions.* The comparison is between mouse mPFC and human DLPFC, two regions that are not homologous [3]. All scores between the species are scores of an mPFC-DLPFC comparison. The mismatch is expected to lower the scores, and it is most important for the 14 genes selected by the screen, whose partner differences could reflect the region and not the species. The datasets contain one region per species, so region and species cannot be separated.
2. *Glial cells.* The glial comparison is underpowered: the human glial cells come from one study with seven control donors, and the analyses of glial types have very few samples. Conservation is detectable in the glial networks, but its strength cannot be measured, and no biological conclusion about the degree of glial conservation can be drawn from these data (Supplementary Note 7). The mouse glial cells are whole cells from the in-house dataset and the human glial cells are nuclei, so the glial comparison between the species is also a comparison of whole cells with nuclei.
3. *Neuronal subtypes.* Networks of single neuronal subtypes are built from 3 to 13 pseudobulk samples and are similarly noisy. We emphasize the analyses of broad classes and of all neurons.
4. *Significance of the global score.* With 13,214 tests, the permutation test of the global score cannot reach small P values, so the screen relies on the test of the local score. That test proved too liberal when compared with genes of similar connectivity. We therefore report the selected genes for hypothesis generation and not as established differences.
5. *Assignment of subtypes between species.* The assignment is based on correlation of expression. A second assignment based on shared marker genes agreed for only 10 of 21 subtypes (6 of 11 inhibitory subtypes, 4 of 9 excitatory subtypes, and not the one subtype assigned to neither class). Assignments of subtypes, including those of excitatory subtypes, are exploratory and depend on the method, and all primary conclusions are based on broad classes.
6. *Dataset and subtype in mouse.* In mouse, 13 of 21 subtypes occur in a single dataset, so differences between datasets and between subtypes cannot be fully separated. The correction model contains the dataset only, and the changes it makes are small.
7. *Protein sequence.* The relation between sequence conservation and network conservation is weak (squared correlation 0.033).
8. *Kinds of networks.* We analyzed co-expression of genes only. Regulatory elements and their target genes [56], networks of regulatory genes [16], networks at the level of transcript isoforms and communication between cells are further layers that were not examined.
9. *Independent dataset.* The independent dataset contains samples from donors with Alzheimer’s disease and from seven control donors, of which only the controls were used. Seven donors are sufficient to test modules but not single genes. The modules tested were those defined on the 5,000 most variable genes, not the modules of the full gene panel. The dataset is human only and does not test the comparison between species.
10. *List of orthologs.* The origin of the list used for the analysis could not be reconstructed; the list itself is part of the analysis package. Two sources shared a Jaccard index of 0.84, and 12,698 of the 13,214 panel genes are one-to-one orthologs in a table retrieved from Ensembl. The analyses were not repeated on that subset.
11. *Disease gene sets.* The sets come from sources of different kinds (SFARI Gene, exome studies, genome-wide association studies and Open Targets), and thresholds such as the score of at least 0.5 in Open Targets are arbitrary. The set for Huntington’s disease was too small to test.
12. *Correction for measurement noise.* The correction assumes that the noise in the two halves of the samples is unrelated. Shared influences would make the corrected value too high.
13. *Post-mortem tissue.* The post-mortem interval, the state before death and the cause of death affect human tissue, have no counterpart in mouse and differ among human datasets. The same-species comparisons capture them only as far as they differ between datasets.
14. *Factor gene sets.* The factor gene sets are themselves derived from genome-wide association studies by assigning variants to genes, and the distinction between genes identified in fetal and in adult brain is imperfect. The set for the compulsive factor is small (47 genes), and its tests are correspondingly weak.
15. *OCD and CTD gene sets.* These sets were discovered through rare variants and are tested here with measures developed on other kinds of sets. The confidence interval of the ratio of adjusted to unadjusted coefficients is wide (0.30 to 1.55), and the excess of genes with low LOEUF is expected because constraint contributed to their discovery.
16. *Adjustment for gene properties.* LOEUF was the only measure of mutational constraint used, and it is estimated with error. Because of this error, adjusting for LOEUF removes the influence of constraint only in part, so the small difference that remains after adjustment may still reflect constraint and cannot be assigned to disease genes specifically. The analyses show that the measured properties together account for the difference; they do not show how much each contributes or that any of them is the cause.
17. *Protein-interaction networks.* PCNet includes connections derived from co-expression, the kind of data analyzed here, and BioGRID restricted to physical interactions covers few gene pairs. A negative result there is weak evidence that no interaction exists.
18. *Cerebellum.* The cerebellum was not examined, because no suitable pair of mouse and human cerebellar snRNA-seq datasets was available. Whether the increased expression of the OCD and CTD genes in postnatal cerebellum [25] and the enrichment of the internalizing factor in Bergmann glia [24] are accompanied by conserved co-expression therefore remains untested.
19. *Ancestry.* The genetic sources (the cross-disorder analysis, the exome study and the population data from which LOEUF is derived) come predominantly from individuals of European ancestry. The conclusions about discovery may not hold for other ancestries.
20. *Matched control sets.* The control sets are matched on measured gene properties only. Disease genes are also studied and annotated more thoroughly than other genes, and this difference is not matched; it could raise or lower the remaining difference.
21. *Developmental data.* The cortical samples of BrainSpan come from several regions and change in cell composition during development, so the difference in expression between the two periods also reflects composition. No donor contributes samples from both periods, so an analysis within donors was not possible.
22. *Whole cells and nuclei.* The in-house mouse dataset consists of whole cells, and the six public datasets consist of nuclei. In mouse, this difference is contained in the dataset term of the model that removed the differences between datasets and in the same-species comparison of the in-house dataset with the other three mouse datasets (Table 2). The median global score between the species changed little when any one dataset, including the in-house dataset, was removed (Fig. S13B). For glial cells, this difference cannot be separated from the difference between the species (see Glial cells above).
23. *Raw data of the in-house dataset.* The raw data of the in-house mouse dataset will be deposited when this article is published after peer review. Until then, the analysis can be repeated from the pseudobulk expression profiles of this dataset in the analysis package, but not from its raw counts.

### Supplementary Note 11. Repetition of the analysis

The analysis was run a second time from the analysis package, independently of the first run. All values of the second run refer to the panel of the 5,000 most variable genes. The median local score was 0.0101 in both runs. The odds ratio for the overlap of mouse and human modules was 10.9. The correlation of the scores of single genes between data with and without removal of dataset differences was 0.577 in the second run and 0.518 in the first. The difference in conservation for the combined disease genes was r = 0.182 in the second run and 0.190 in the first. Sample number explained 91% to 101% of the difference between neurons and glial cells in the second run and 74% to 87% in the first. An independent matching of orthologs recovered 4,812 of the 5,000 genes. The second run led to four recommendations, each of which motivated an analysis reported here: the disease analysis on all expressed genes, the control for sample number in glial cells, the documentation of the settings of the module analysis, and the check of the model used to remove differences between datasets. Because the second run used the same data and the same code, it is a check of reproducibility and not an independent replication.

### Supplementary tables

**Table S1.** Conservation of each of the 11 disease gene sets.

| Gene set | Genes in panel | All neurons: r (FDR) | Excitatory neurons: r (FDR) | Inhibitory neurons: r (FDR) | After adjustment, all neurons: coefficient (P; FDR) |
| --- | --- | --- | --- | --- | --- |
| All 11 sets combined | 539 | 0.102 ($1.7 \times 10^{-4}$) | 0.067 (0.013) | 0.057 (0.025) | 0.0049 (0.039; single test) |
| Intellectual disability | 71 | 0.310 ($2.2 \times 10^{-4}$) | 0.157 (0.069) | 0.119 (0.17) | 0.0168 (0.021; 0.23) |
| Autism spectrum disorder | 90 | 0.234 ($2.2 \times 10^{-3}$) | 0.181 (0.017) | 0.169 (0.027) | 0.0097 (0.106; 0.58) |
| Epilepsy | 65 | 0.250 ($5.4 \times 10^{-3}$) | 0.181 (0.049) | 0.231 (0.009) | 0.0043 (0.599; 0.86) |
| Major depressive disorder | 39 | 0.163 (0.17) | 0.027 (0.79) | 0.163 (0.17) | 0.0108 (0.211; 0.78) |
| Schizophrenia | 59 | 0.100 (0.27) | 0.256 ($5.7 \times 10^{-3}$) | 0.171 (0.069) | -0.0013 (0.863; 0.86) |
| Bipolar disorder | 43 | 0.087 (0.42) | 0.129 (0.24) | 0.162 (0.17) | -0.0017 (0.835; 0.86) |
| Attention-deficit/hyperactivity disorder | 26 | 0.043 (0.78) | -0.057 (0.73) | -0.081 (0.58) | -0.0038 (0.711; 0.86) |
| Alzheimer's disease (genome-wide association study) | 65 | -0.003 (0.96) | 0.022 (0.79) | -0.069 (0.42) | 0.0024 (0.703; 0.86) |
| Alzheimer's disease (Open Targets) | 135 | -0.020 (0.78) | -0.071 (0.25) | -0.114 (0.069) | 0.0036 (0.398; 0.86) |
| Parkinson's disease (Open Targets) | 82 | 0.087 (0.26) | 0.093 (0.24) | 0.121 (0.16) | 0.0058 (0.369; 0.86) |
| Amyotrophic lateral sclerosis (Open Targets) | 60 | 0.095 (0.28) | 0.125 (0.18) | 0.120 (0.20) | 0.0038 (0.586; 0.86) |

**Table S2.**
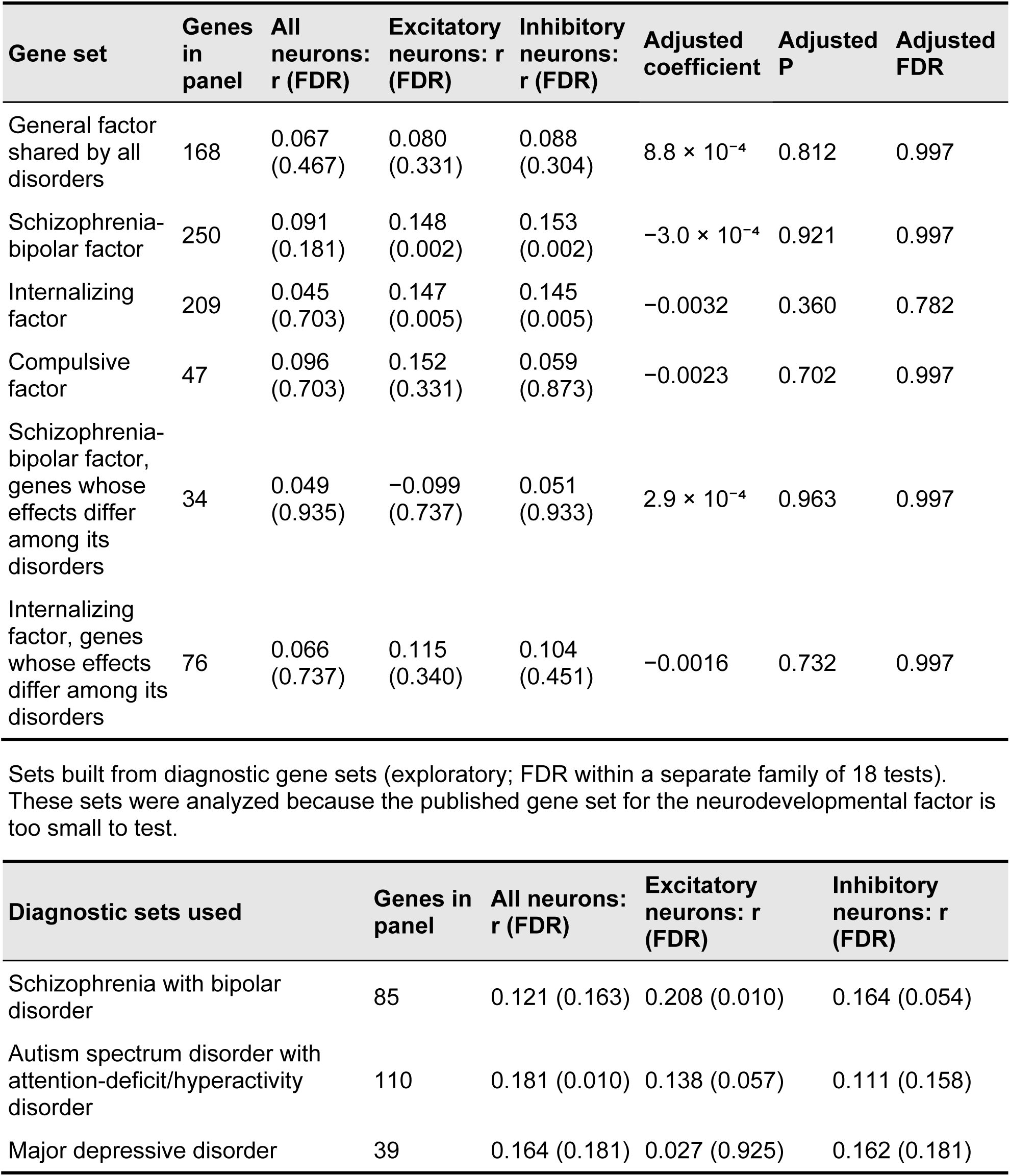
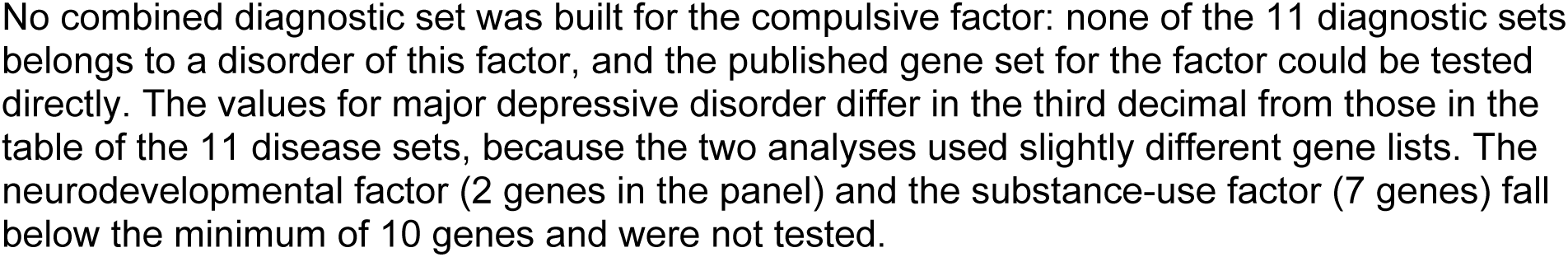
Conservation of gene sets for genomic factors of psychiatric risk: all sets. Each cell gives the rank-biserial correlation r of the conservation score for the set compared with all other genes, with the false discovery rate (FDR) in parentheses. The adjusted model contains membership in the set and five gene properties (LOEUF, mean expression, expression variance, log10 gene length and GC content; all neurons; standard errors robust to unequal variance). The FDR was computed within the family of 66 tests of factor sets and of gene sets for obsessive-compulsive disorder and chronic tic disorders. LOEUF, loss-of-function observed/expected upper bound fraction; GC content, proportion of guanine and cytosine bases.

**Table S3.** Matched control gene sets, and the gene sets for obsessive-compulsive disorder (OCD) and chronic tic disorders (CTD). *A. Disease gene sets compared with matched control gene sets.* For each disease set, 1,000 control sets were drawn in which every gene was replaced by a gene similar in LOEUF, mean expression, expression variance, gene length and GC content. The P value is the proportion of control sets with an r at least as high as that of the disease set (one-sided). r, rank-biserial correlation of the conservation score compared with all other genes; FDR, false discovery rate within the family of six tests on constraint and matched controls. Only the genes for which all five gene properties are available were used. *B. High-confidence gene sets for OCD and CTD.* The sets are those of the exome study [25] (false discovery rate, q, below 0.1 in the transmission and de novo association test of that study): 12 genes for OCD, 10 for CTD and 34 for the two disorders combined; the three sets together contain 36 genes, 34 of which are in the panel. The set for the two disorders combined, in all neurons, was designated the main test. r, rank-biserial correlation of the conservation score compared with all other genes; FDR, false discovery rate within the family of 66 tests of factor sets and of OCD and CTD sets. The adjusted model contains the five gene properties.

| Gene set | Genes | Observed $r$ | Control sets: mean [95% interval] | $z$ | P | FDR |
| --- | --- | --- | --- | --- | --- | --- |
| Seven neuropsychiatric sets combined | 238 | 0.172 | 0.159 [0.103, 0.213] | 0.48 | 0.309 | 0.907 |
| Intellectual disability | 61 | 0.312 | 0.179 [0.079, 0.282] | 2.55 | 0.009 | 0.054 |
| Autism spectrum disorder | 84 | 0.222 | 0.218 [0.133, 0.300] | 0.09 | 0.466 | 0.907 |
| Epilepsy | 55 | 0.190 | 0.217 [0.114, 0.318] | -0.51 | 0.692 | 0.907 |

| Gene set | Gene<br>s in<br>panel | All<br>neurons<br>: $r$ | P | FDR | Excitatory<br>: $r$ | Inhibitory<br>: $r$ | Adjusted<br>coefficient<br>t | Adjusted<br>P | Adjusted<br>FDR |
| --- | --- | --- | --- | --- | --- | --- | --- | --- | --- |
| OCD and<br>CTD<br>combined<br>(main<br>test) | 32 | 0.224 | 0.028 | 0.266 | 0.017 | 0.015 | 0.0129 | 0.050 | 0.304 |
| All 36<br>genes | 34 | 0.218 | 0.028 | 0.266 | 0.027 | 0.006 | 0.0126 | 0.062 | 0.331 |
| All 36<br>genes<br>without<br>four<br>genes<br>with more<br>variants<br>in control<br>individuals | 30 | 0.217 | 0.040 | 0.293 | n.c. | n.c. | 0.0102 | 0.159 | 0.523 |
| OCD | 11 | -0.105 | 0.548 | 0.904 | -0.215 | -0.216 | -0.0074 | 0.588 | 0.925 |
| CTD | 10 | 0.127 | 0.486 | 0.873 | 0.195 | 0.001 | 0.0055 | 0.757 | 0.997 |

The ratio of the adjusted coefficient to the unadjusted coefficient was 0.90 (95% confidence interval 0.30 to 1.55; 5,000 resamplings of genes). A ratio of 1 means that the gene properties account for none of the difference; the interval is too wide to decide. Twelve of the 34 genes in the panel are among the tenth of genes with the lowest LOEUF; this is expected, because the tests that identified these genes favor genes under strong constraint. n.c., not computed. Excitatory and Inhibitory refer to excitatory and inhibitory neurons. The four genes removed in the third row are *ABCD2*, *DNAH9*, *DNAH10* and *TEP1*. LOEUF, loss-of-function observed/expected upper bound fraction; GC content, proportion of guanine and cytosine bases.

**Table S4.** The 14 disease genes selected by the screen and the outcome of the four checks (13,214 genes).

| Gene | Disease set | Selected in | Check 1 | Check 2 | Check 3 | Check 4 (P) | Connectivity in human |
| --- | --- | --- | --- | --- | --- | --- | --- |
| *BCL11A* | Autism spectrum disorder; intellectual disability | All neurons | Passed | Passed | Not evaluable | 0.96 | 0.34 |
| *FOXP2* | Attention-deficit/hyperactivity disorder (see note) | All neurons; inhibitory neurons | Passed | Passed | Not evaluable | 0.81 | 0.96 |
| *MCTP1* | Bipolar disorder | Excitatory neurons | Passed | Passed | Not evaluable | 0.96 | 0.70 |
| *NEGR1* | Major depressive disorder; also Alzheimer's disease (Open Targets) | Excitatory neurons | Passed | Passed | Not evaluable | 0.69 | 0.94 |
| *SRCAP* | Autism spectrum disorder; intellectual disability | Inhibitory neurons | Passed | Passed | Not evaluable | 0.95 | 0.74 |
| *CTNND2* | Autism spectrum disorder | All neurons | Failed | Failed | Not evaluable | 0.70 | 0.82 |
| *DRD4* | Attention-deficit/hyperactivity disorder | Inhibitory neurons | Failed | Passed | Not evaluable | 0.61 | 0.16 |
| *FHIT* | Major depressive disorder | All neurons | Failed | Passed | Not evaluable | 0.94 | 0.57 |
| *GABRB2* | Schizophrenia | All neurons | Failed | Passed | Not evaluable | 0.86 | 0.88 |
| *KCNJ6* | Attention-deficit/hyperactivity disorder | All neurons | Failed | Passed | Not evaluable | 0.70 | 0.19 |
| *NFIA* | Autism spectrum disorder; intellectual disability | Excitatory neurons | Failed | Passed | Not evaluable | 0.76 | 0.48 |
| *PRKD1* | Schizophrenia | Excitatory neurons | Failed | Passed | Not evaluable | 0.88 | 0.22 |
| *SLC4A7* | Major depressive disorder | Inhibitory neurons | Failed | Passed | Not evaluable | 0.56 | 0.94 |
| *TLK2* | Autism spectrum disorder; intellectual disability | All neurons | Failed | Passed | Not evaluable | 0.91 | 0.48 |

Check 1, the gene meets the two criteria of the screen with and without removal of differences between datasets. Check 2, the selection is not explained by differences in the composition of cell subtypes. Check 3, the 50 strongest human partners are found again in the independent human dataset. Check 4, P value of the comparison with genes of similar connectivity. Connectivity in human, rank of the gene's connectivity in the human network as a fraction (1 is the most connected gene). Check 3 was not evaluable for any gene, because the independent dataset has seven donors, too few to reproduce the 50 strongest partners of a single gene (the overlap of partners between the main data and the independent dataset was 0.000 to 0.020 for these genes). No gene passed check 4 (P from 0.56 to 0.96), so the list serves only to generate hypotheses. *FOXP2* is known as a gene for an inherited disorder of speech and language [57]; it is in the set for attention-deficit/hyperactivity disorder because it was named in a genome-wide association study, which does not establish a role in that disorder.

**Table S5.** The 14 selected genes in networks of protein interactions. PCNet version 1.4 combines many kinds of evidence, including co-expression. BioGRID version 5.0.262 was restricted to physical interactions; mouse interactions were transferred to human gene names through one-to-one orthologs. Each observed value was compared with 1,000 sets of comparison genes matched in co-expression connectivity; the P value is the proportion of comparison sets with a value at least as high (one-sided). FDR, false discovery rate within the family of five tests.

| Test | Observed | Comparison sets:<br>mean ± SD | P | FDR |
| --- | --- | --- | --- | --- |
| Number of connections in PCNet,<br>against matched genes | 301 | 424 ± 112 | 0.877 | 0.877 |
| Number of connections in PCNet,<br>against matched disease genes | 301 | 418 ± 109 | 0.865 | 0.877 |
| Overlap of physical partners<br>between human and mouse (Jaccard<br>index, BioGRID) | 0.0147 | 0.0159 ± 0.0082 | 0.510 | 0.877 |
| Fraction of the co-expression<br>partners present in human only that<br>are neighbors in PCNet | 0.0456 | 0.0212 (SD not<br>available) | 0.020 | 0.100 |
| Fraction of the co-expression<br>partners present in human only that<br>are neighbors in BioGRID | 0.0077 | 0.0094 (SD not<br>available) | 0.737 | 0.877 |
| Retention of PCNet after matching<br>through orthologs (descriptive) | 0.826 | not applicable | not<br>applicable | not<br>applicable |

No test was significant after correction for multiple tests. The result in PCNet that was significant before correction ( $P = 0.020$ ) was not found in BioGRID ( $P = 0.74$ ), and it probably arises from the connections in PCNet that are themselves derived from co-expression, the kind of data from which the genes were selected. SD, standard deviation.

**Table S6.** Simulation of statistical power at the glial sample numbers, and chance values for the glial data. *A. Neuronal data with conservation lowered stepwise and reduced to the glial sample numbers.* The level of conservation is the median global score at full sample number (180 mouse and 320 human samples) after the gene labels of the stated share of genes were exchanged in the human data. For each level, the table gives the median global score in 200 random selections of samples (mean, with the range that contains 95% of the selections) and the statistical power, which is the share of selections whose median exceeds the 95th percentile of the chance medians. Genes: the 5,000 genes analyzed in glial cells. *B. The glial data: observed median global score and chance values.* The chance values are the medians obtained after the gene labels of the human glial network were permuted (200 permutations). P is the share of permutations with a median at least as high as the observed one; 0.005 is the smallest value attainable with 200 permutations. The observed values were recomputed from the stored glial expression profiles with the code of the simulation. They differ slightly from the values of the original analysis that are quoted in the text (0.004 to 0.016 for single types and 0.064 for all types together), whose computation could not be reconstructed in every detail.

| Level of conservation at full sample number | Share of genes with exchanged labels | 7 human and 15 mouse samples: score [95% range] | Power | 34 human and 74 mouse samples: score [95% range] | Power |
| --- | --- | --- | --- | --- | --- |
| 0.190 | 0% | 0.031 [0.011, 0.059] | 1.00 | 0.094 [0.074, 0.124] | 1.00 |
| 0.107 | 25% | 0.017 [0.006, 0.037] | 1.00 | 0.054 [0.041, 0.072] | 1.00 |
| 0.058 | 50% | 0.009 [0.002, 0.020] | 0.98 | 0.028 [0.021, 0.039] | 1.00 |
| 0.035 | 62.5% | 0.005 [0.001, 0.012] | 0.93 | 0.017 [0.011, 0.024] | 1.00 |
| 0.018 | 75% | 0.003 [0.000, 0.008] | 0.79 | 0.009 [0.005, 0.013] | 1.00 |
| 0.010 | 87.5% | 0.002 [-0.001, 0.004] | 0.45 | 0.005 [0.002, 0.008] | 0.99 |
| chance | 100% | 0.000 [-0.002, 0.002] | 0.05 | 0.000 [-0.001, 0.002] | 0.05 |

| Glial cells | Human samples | Mouse samples | Observed median global score | 95th percentile of chance | P |
| --- | --- | --- | --- | --- | --- |
| Astrocyte | 7 | 15 | 0.006 | 0.001 | 0.005 |
| Oligodendrocyte | 7 | 16 | 0.018 | 0.001 | 0.005 |
| Oligodendrocyte precursor cells | 7 | 14 | 0.003 | 0.001 | 0.005 |
| Microglia | 7 | 15 | 0.013 | 0.001 | 0.005 |
| Endothelial | 6 | 14 | 0.007 | 0.001 | 0.005 |
| All glial types together | 34 | 74 | 0.073 | 0.001 | 0.005 |

**Table S7.** Influence of the list of orthologs.

| Comparison | Genes or gene pairs in the first list | In the second list | In both | Jaccard index |
| --- | --- | --- | --- | --- |
| List of one-to-one orthologs used for the analysis, compared with a list retrieved from Ensembl BioMart | 17,392 | 17,067 | 15,720 | 0.839 |
| Primary panel of expressed genes, compared with a panel built from the Ensembl list that also admits genes with several counterparts | 13,214 | 12,880 | 12,880 | 0.975 |

Of the 12,880 genes of the panel built from the Ensembl list, 12,698 are one-to-one orthologs in that list and 182 have several counterparts (167 in one species and 15 in both). The source of a list of orthologs is therefore more important than the rule for genes with several counterparts. The origin of the list used for the analysis could not be reconstructed; the list itself is part of the analysis package.

**Table S8.** Families of tests within which the false discovery rate was controlled. Within each family, the Benjamini-Hochberg method was applied to the P values of the tests listed, and a result was called significant at a false discovery rate (FDR) below 0.05. The families were defined before the analysis.

| Family | Tests in the family | Number of tests | Where reported |
| --- | --- | --- | --- |
| 1 | The 11 disease gene sets combined, compared with all other genes, in all neurons, in excitatory neurons and in inhibitory neurons | 3 | Table 3, first row |
| 2 | Each of the 11 disease gene sets in the same three analyses | 33 | Tables 3 and S1; Fig. 4A |
| 3 | Coefficient of each of the 11 disease gene sets after adjustment for the five gene properties (all neurons) | 11 | Table S1; Fig. S3A |
| 4 | Expression variance of each of the 11 disease gene sets compared with other genes | 11 | Fig. S4 |
| 5 | Overlap of each of the four mouse modules with each of the five human modules | 20 | Fig. 3C |
| 6 | Factor gene sets (six sets in three analyses, before and after adjustment; 36 tests), gene sets for obsessive-compulsive disorder and chronic tic disorders (26 tests), and genes with differing effects compared with genes with shared effects (4 tests) | 66 | Tables 3, S2 and S3; Fig. S5 |
| 7 | Sets built from diagnostic gene sets (three sets in three analyses, before and after adjustment) | 18 | Table S2 |
| 8 | Constraint and matched controls: four comparisons with matched control sets and two tests across levels of constraint (interaction and trend) | 6 | Tables 3 and S3 |
| 9 | Tests of the selected genes in networks of protein interactions | 5 | Table S5 |
| 10 | Developmental tests (gene sets and correlation with the conservation score) | 8 | Supplementary Note 8 |
| 11 | Test of the local score of every gene of the panel, separately in each cell-class analysis | 13,214 in each analysis | Supplementary Note 5 |
| 12 and 13 | The 14 selected genes tested again with 30 strongest partners (family 12) and with 100 strongest partners (family 13) in place of 50 | 14 in each family | Analysis package |

The single test of the 11 sets combined after adjustment ( $P = 0.039$ ) belongs to no family and is reported without correction. Three further sets of tests of the original analysis are not listed, because their tests are not recorded in one result table of the analysis package: groupings of

### Legends of supplementary figures

**Figure S1.**
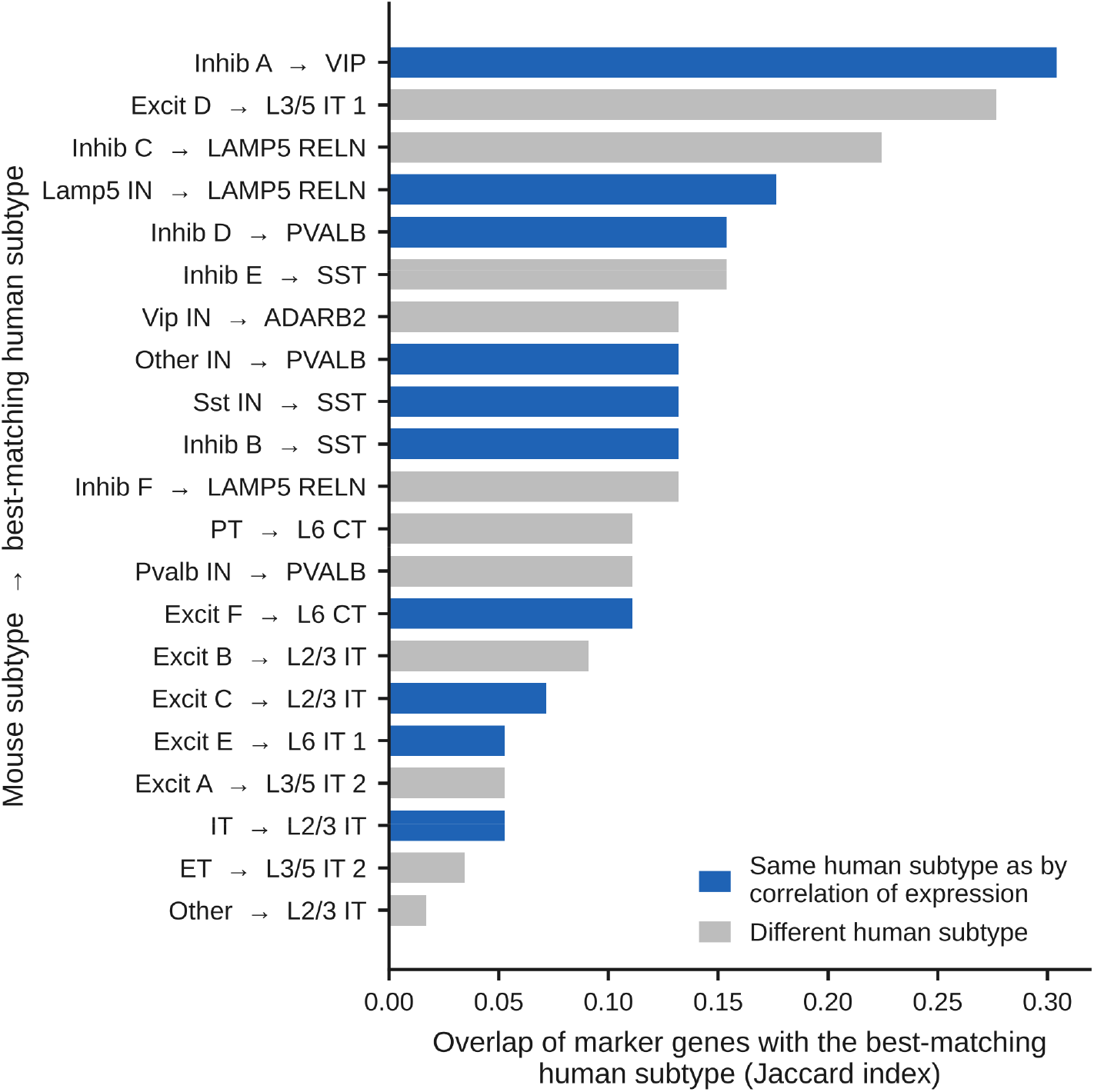
Assignment of mouse subtypes to human subtypes by shared marker genes. For each mouse subtype, the overlap of marker genes (Jaccard index) with the best-matching human subtype, which is named after the arrow; blue, the same human subtype as in the assignment based on correlation of expression (10 of 21 subtypes). Subtype names are those of the source studies. L followed by numbers, cortical layers; IT, intratelencephalic; ET, extratelencephalic; PT, pyramidal tract; CT, corticothalamic; IN, interneuron; Excit and Inhib followed by a letter, excitatory and inhibitory subtypes that the source study distinguishes by letters. VIP, SST, PVALB, LAMP5, RELN and ADARB2 are marker genes of inhibitory subtypes, written in the mouse form (for example Vip) in the names of mouse subtypes. A number at the end of a name distinguishes subtypes that share the name; Other and Other IN are subtype names of the source study.

**Figure S2.**
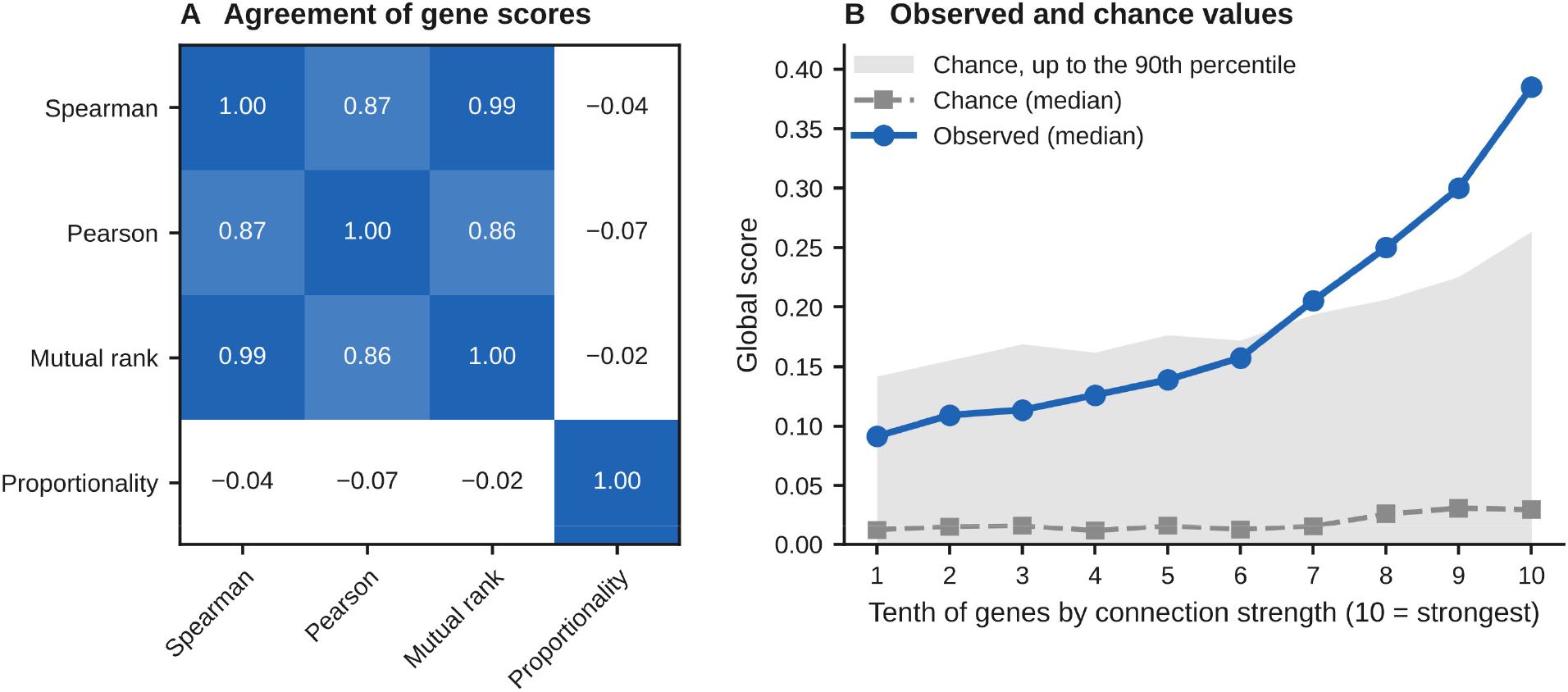
Scores obtained with different measures of association, and chance values. (A) Agreement of the scores of single genes among networks based on the Spearman correlation, the Pearson correlation, mutual rank and proportionality (correlation of the global scores; values below 0 are drawn in white). (B) Median of the observed global score (blue) compared with the score expected by chance (the profile of each gene in mouse compared with the profile of a randomly chosen gene in human; grey line, median; grey band, values up to the 90th percentile), within tenths of the genes ordered by connection strength.

**Figure S3.**
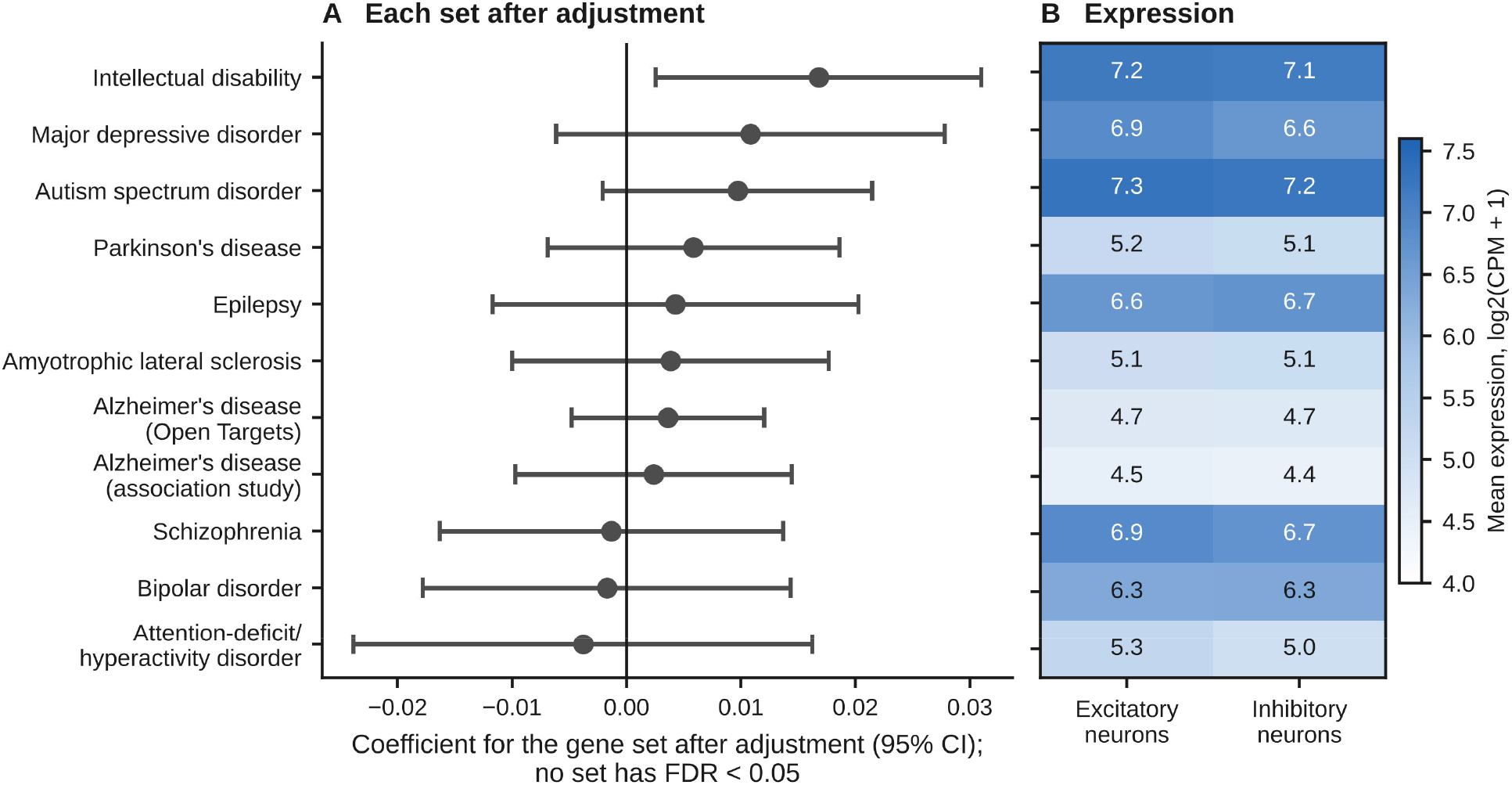
Disease gene sets one by one. (A) Coefficient for membership in each disease gene set after adjustment for the five gene properties, with 95% confidence intervals (CI); no set has a false discovery rate (FDR) below 0.05. (B) Mean expression of the genes of each disease set in excitatory and inhibitory neurons; rows as in A. CPM, counts per million.

**Figure S4.**
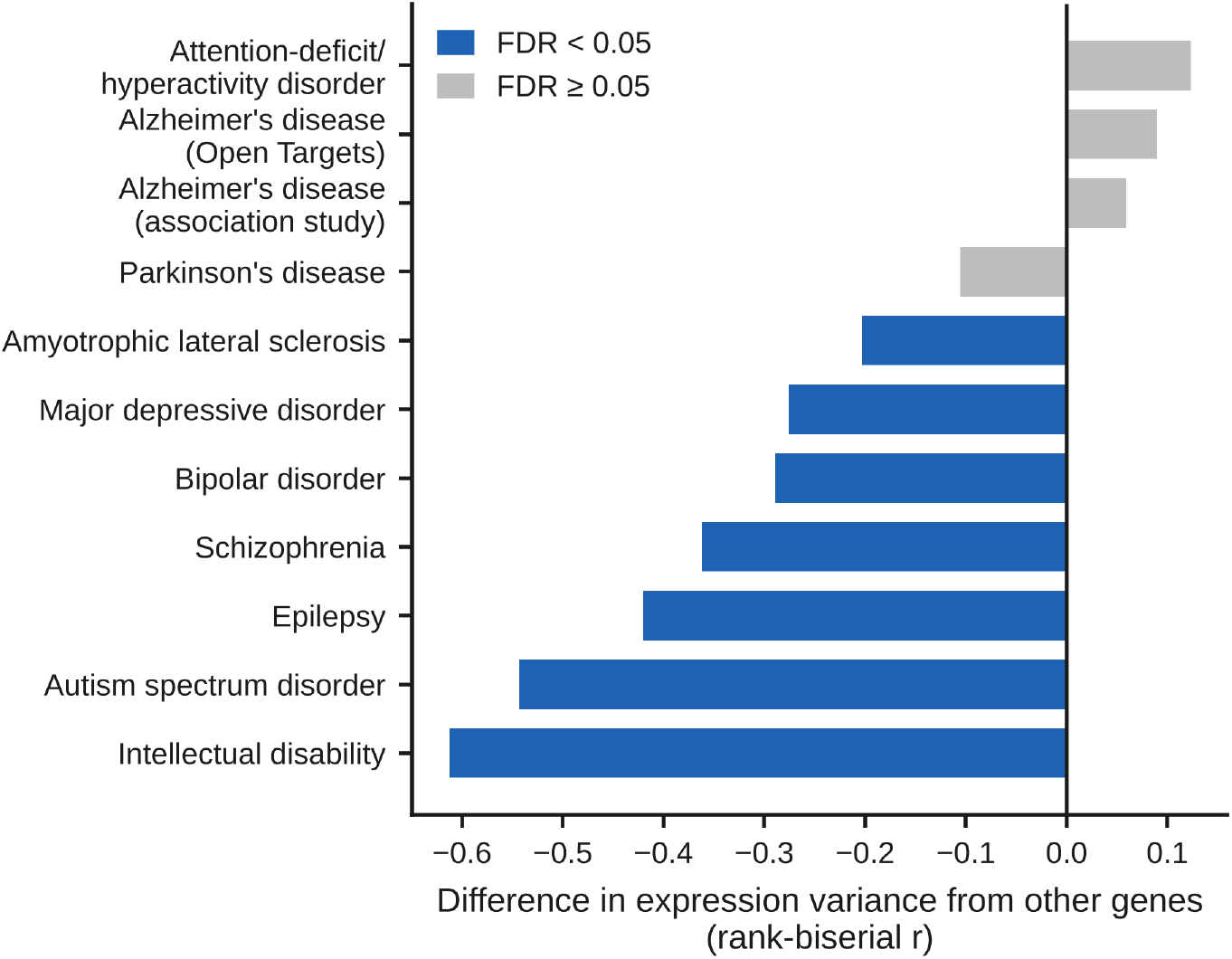
Expression variance of disease gene sets. Difference in expression variance between each disease gene set and all other genes (rank-biserial correlation, r; negative values indicate lower variance in the disease set). Blue, false discovery rate (FDR) below 0.05.

**Figure S5.**
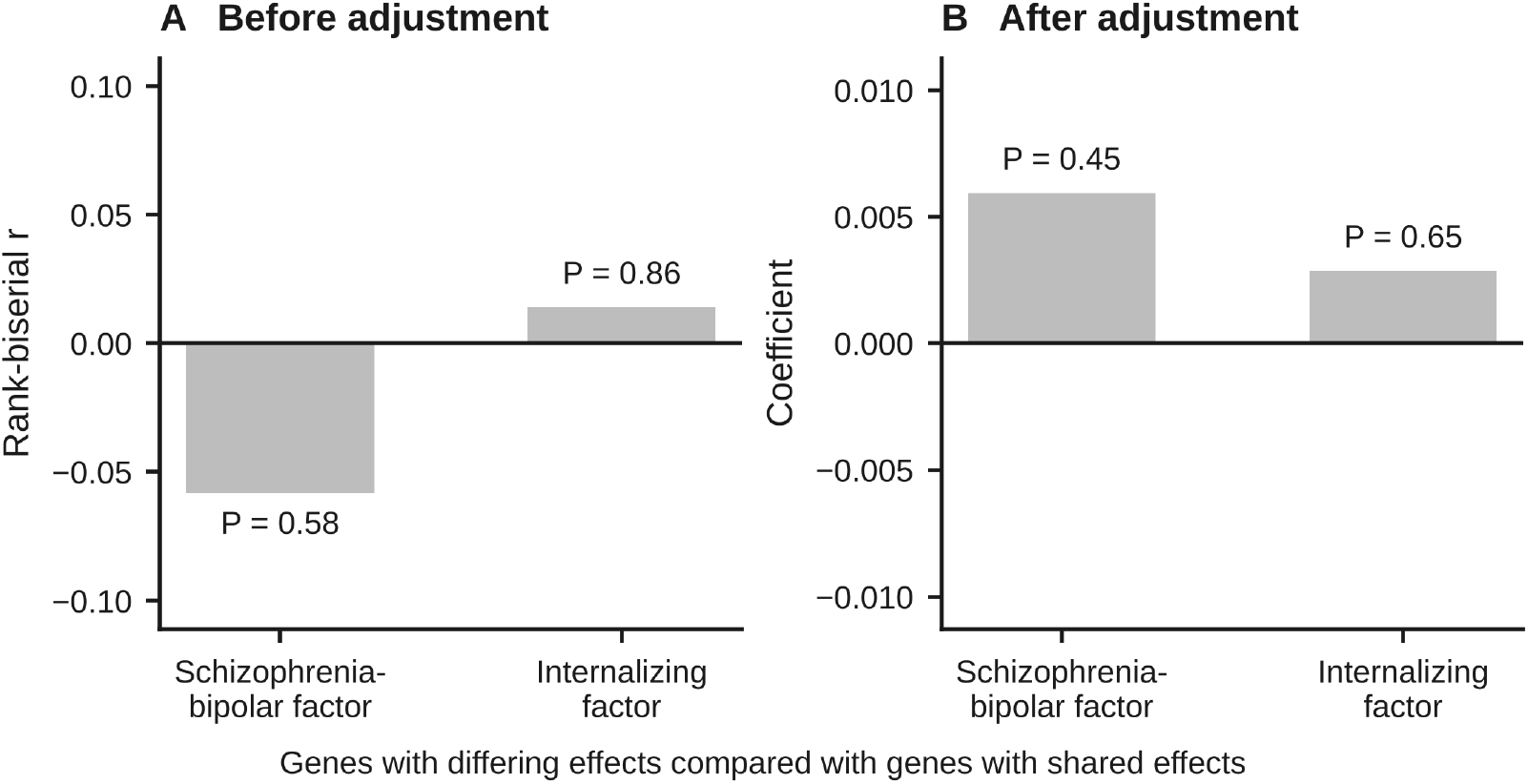
Genes whose effects differ among the disorders of a factor. The genes whose effects differ among the disorders of the schizophrenia-bipolar factor and of the internalizing factor are each compared with the genes shared by the disorders of that factor. (A) Difference in conservation before adjustment (rank-biserial correlation, r; positive values indicate higher conservation of the genes with differing effects). (B) Coefficient after adjustment for the five gene properties. The P values printed in the panels are not corrected for multiple tests. Neither comparison is significant.

**Figure S6.**
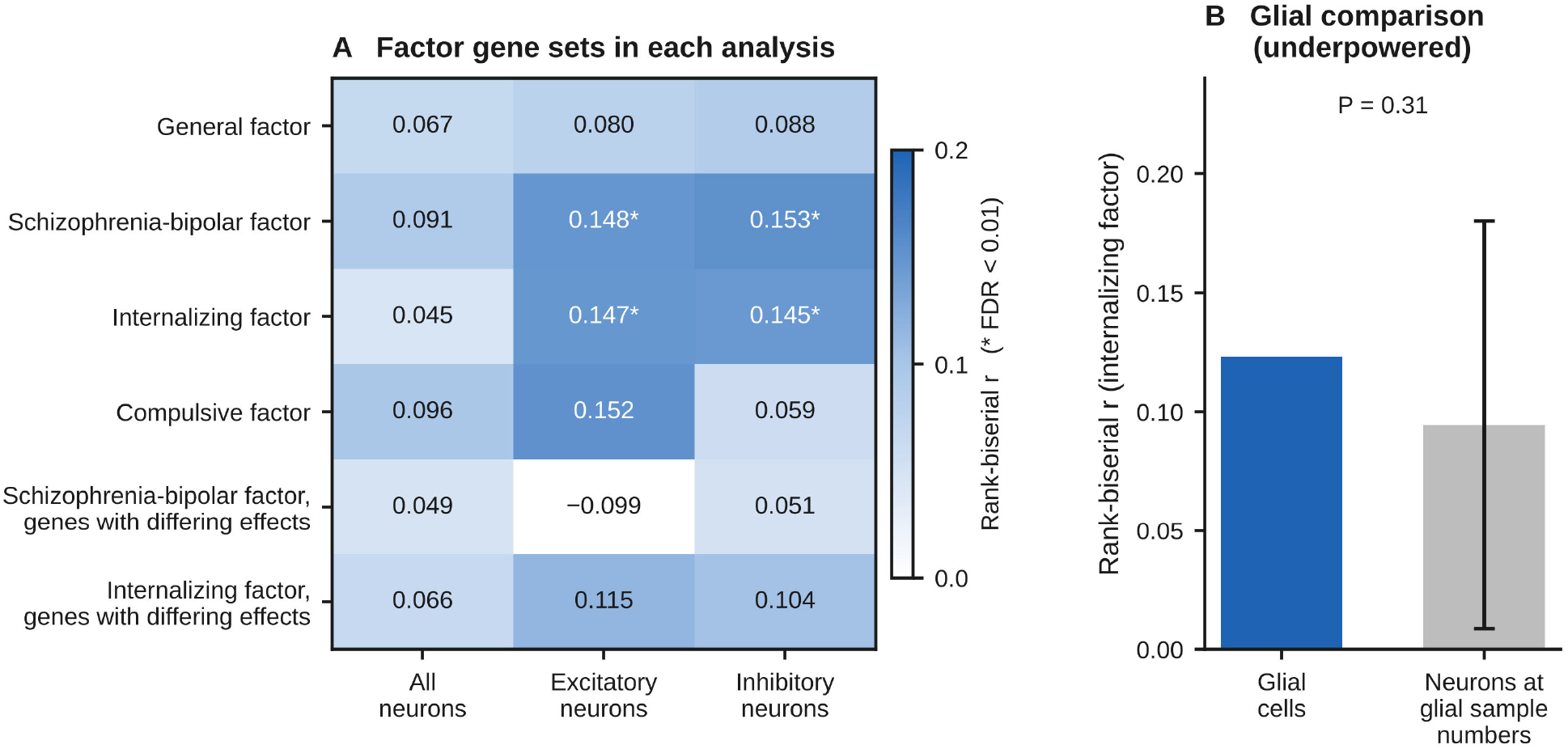
Factor gene sets in each cell class, and the internalizing factor in glial cells. (A) Difference in conservation between each factor gene set and all other genes in all neurons, excitatory neurons and inhibitory neurons (rank-biserial correlation, r; values are printed in the panel, and values below 0 are drawn in white; asterisks, false discovery rate, FDR, below 0.01). The last two rows are the genes whose effects differ among the disorders of a factor. (B) Underpowered glial comparison: difference for the internalizing factor in glial cells (blue), compared with the differences obtained for the same gene set in neuronal data reduced to the glial sample numbers (grey; mean and two standard deviations; P = 0.31); at the present sample number this test is uninformative.

**Figure S7.**
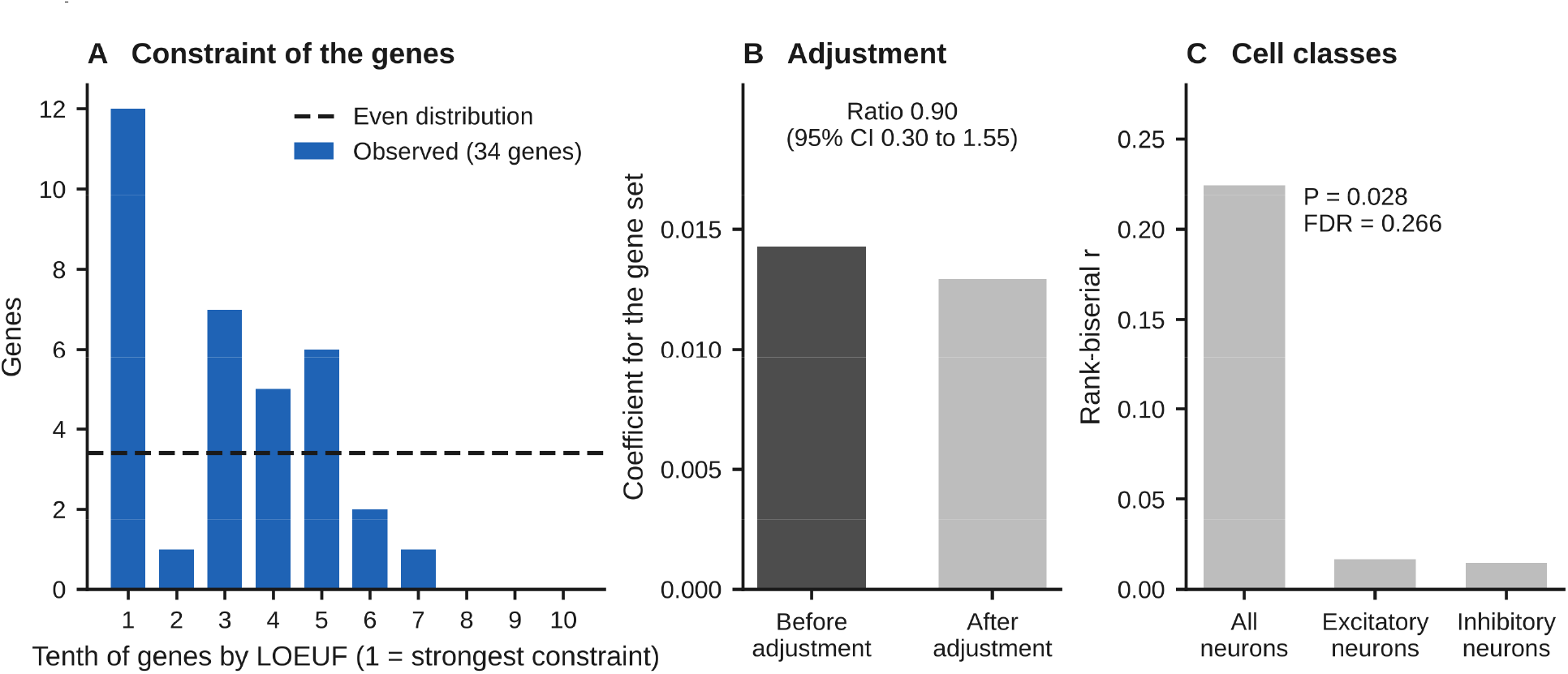
Gene sets for obsessive-compulsive disorder (OCD) and chronic tic disorders (CTD). (A) Distribution of the 34 genes of the three high-confidence sets [25] that are in the panel across tenths of all genes ordered by the loss-of-function observed/expected upper bound fraction (LOEUF; 1 is the tenth under the strongest constraint), compared with an even distribution (dashed line, 3.4 genes in each tenth). The excess of genes under strong constraint is expected, because constraint contributed to the discovery of these genes. (B) Coefficient for the main gene set (the genes linked to the two disorders combined, 32 genes in the panel) in all neurons before and after adjustment for the five gene properties; the ratio of the two is 0.90 (95% confidence interval, CI, 0.30 to 1.55). (C) Difference in the conservation score between the main gene set and all other genes (rank-biserial correlation, r), in all neurons (P = 0.028 before correction for multiple tests; false discovery rate, FDR, 0.266) and in excitatory and inhibitory neurons (no difference).

**Figure S8.**
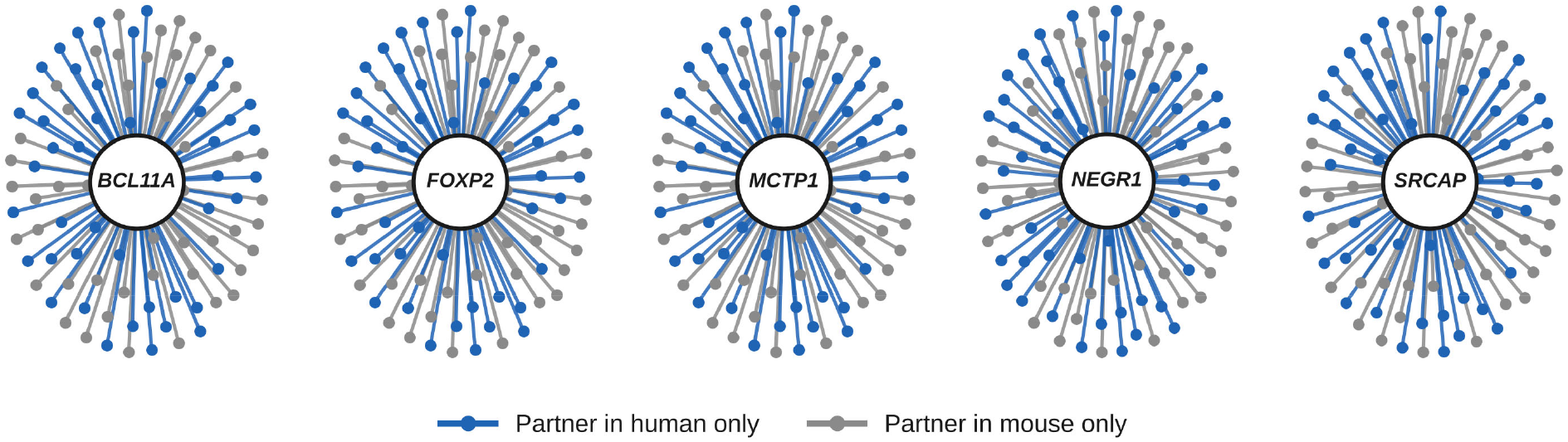
Partners of the five selected genes that passed the first two checks. (*BCL11A*, *FOXP2*, *MCTP1*, *NEGR1* and *SRCAP*). Each line connects the gene to one partner. Blue, those of the gene’s 50 strongest partners in human that are not among its 50 strongest partners in mouse; grey, the reverse. Each gene has 47 to 49 partners of each kind.

**Figure S9.**
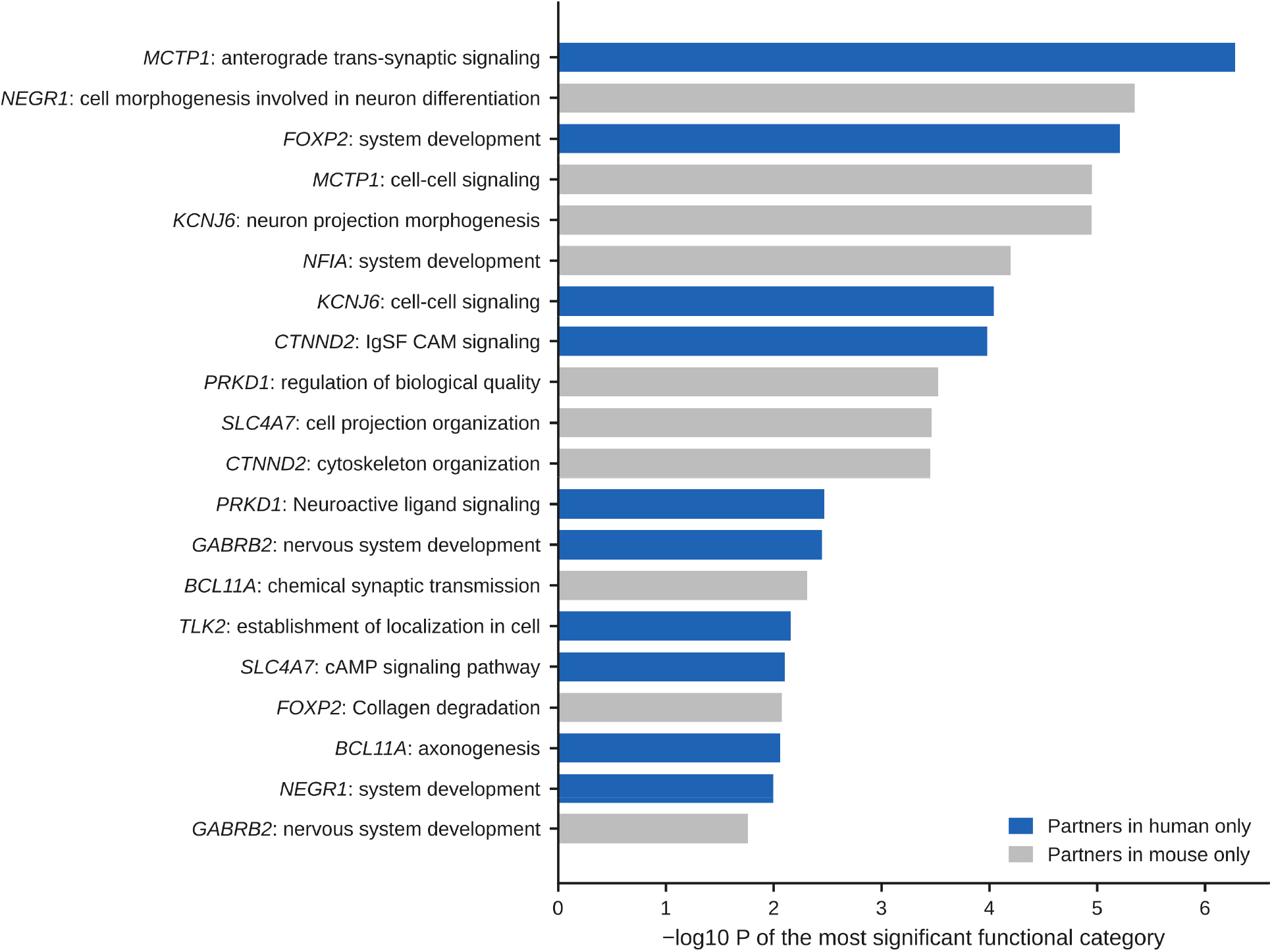
Functional categories enriched among the partners that differ between the species. For each of the 14 selected genes, the partners present in human only (blue) and the partners present in mouse only (grey) form two sets. For each set, the most significant category from g:Profiler was determined (Gene Ontology biological process, the Kyoto Encyclopedia of Genes and Genomes or Reactome), and the 20 sets with the most significant categories are shown. Each row gives the selected gene and the category, named as in the source database (IgSF CAM, cell adhesion molecules of the immunoglobulin superfamily; cAMP, cyclic adenosine monophosphate). The P values are those returned by g:Profiler.

**Figure S10.**
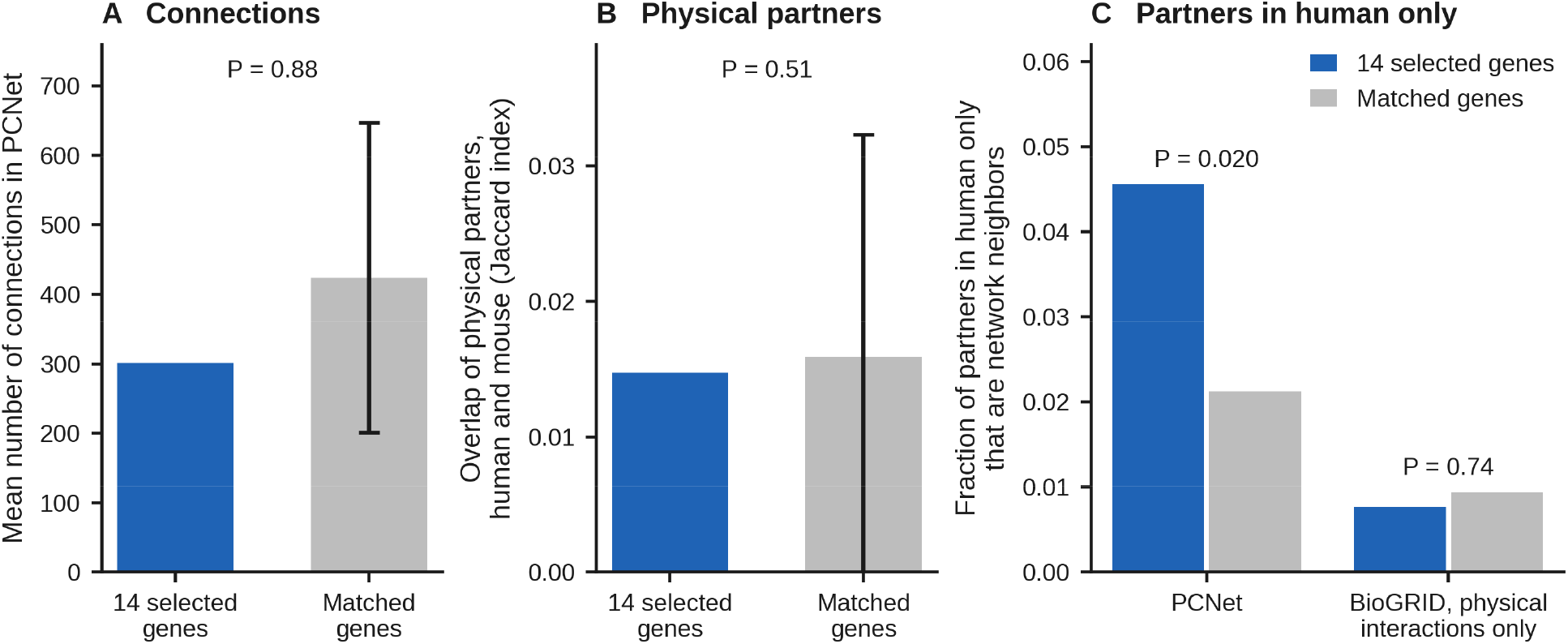
The 14 selected genes in networks of protein interactions. Blue, the selected genes; grey, matched comparison genes (mean; the error bars in A and B are two standard deviations of the comparison sets). (A) Mean number of connections of the selected genes in PCNet (P = 0.88). (B) Overlap of physical partners between human and mouse in BioGRID (P = 0.51). (C) Fraction of the co-expression partners present in human only that are neighbors of the selected gene in PCNet (P = 0.020 before correction for multiple tests; false discovery rate 0.10) and in BioGRID restricted to physical interactions (P = 0.74). No test is significant after correction for multiple tests.

**Figure S11.**
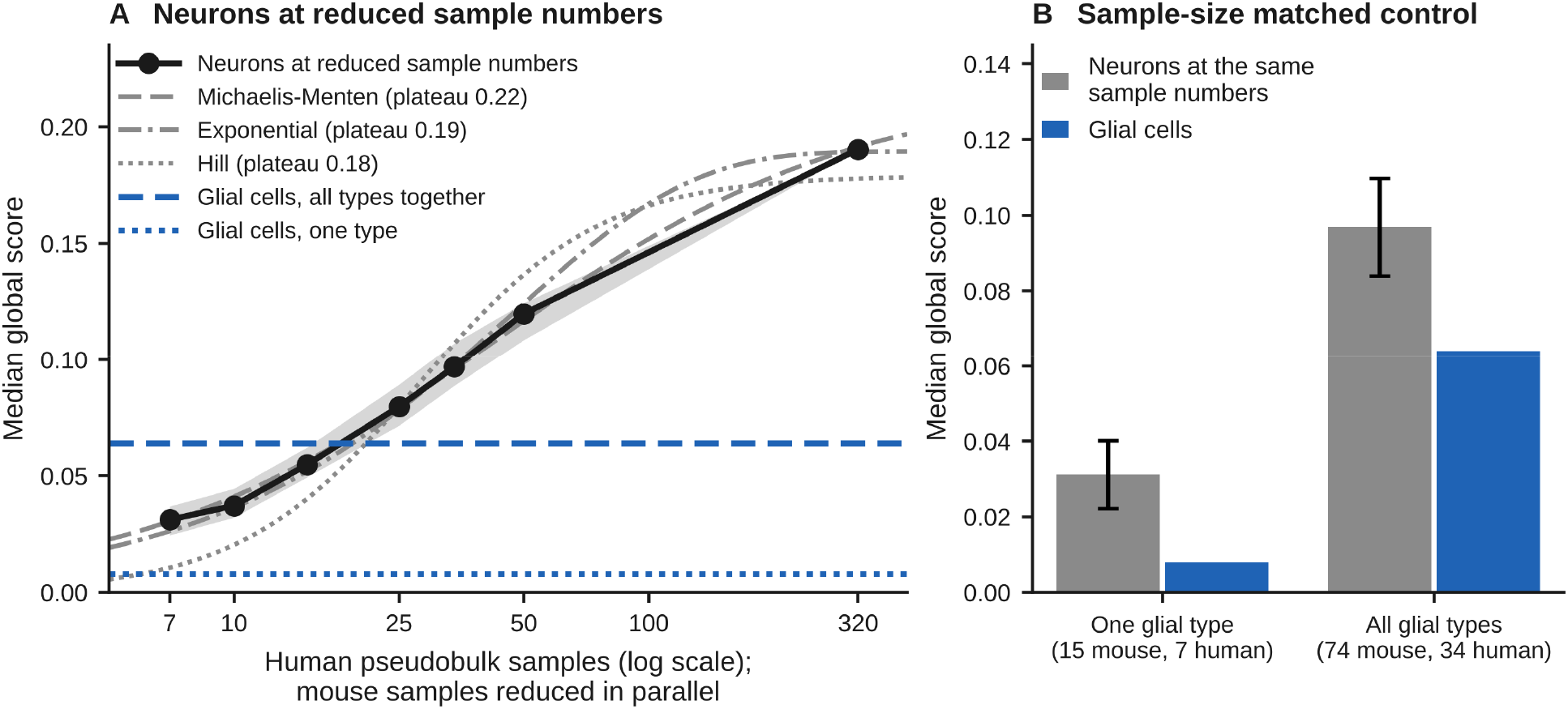
Control for sample number: the global score at reduced numbers of samples. The glial comparison is underpowered for measuring the strength of conservation, and the figure shows what the number of samples does to the score; it does not show a biological difference between glial cells and neurons. (A) Median global score of neuronal data reduced to increasing numbers of pseudobulk samples (black; shading, interquartile range of the random selections). The horizontal axis gives the number of human samples; the number of mouse samples was reduced in parallel (15, 20, 30, 50, 74 and 100 mouse samples for 7, 10, 15, 25, 34 and 50 human samples; the last point is the complete data, 180 mouse and 320 human samples). Three fitted curves (Michaelis-Menten, exponential and Hill) approach a plateau (grey; the plateaus of 0.18 to 0.22 are extrapolations). Blue lines, the observed values of glial cells (0.064 for all glial types together and 0.008 for one glial type; the five glial types range from 0.004 to 0.016). (B) Neurons reduced to the glial sample numbers (grey; error bars, standard deviation across 50 random selections) compared with glial cells (blue).

**Figure S12.**
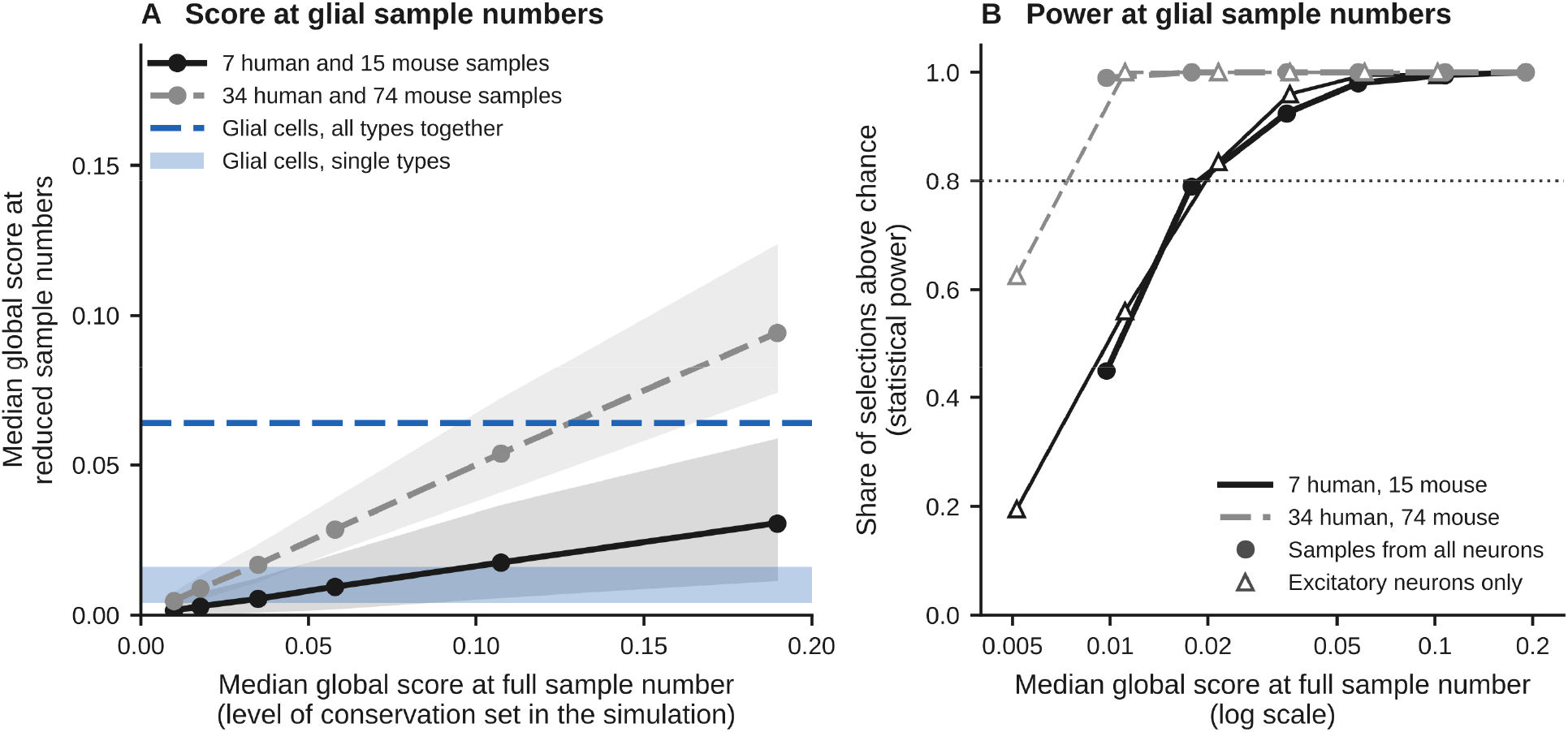
Underpowered glial comparison: what the glial sample numbers can detect. Simulation with neuronal data in which the conservation was lowered stepwise (Supplementary Note 7). (A) Median global score at reduced sample numbers plotted against the level of conservation, which is the median global score of the same data at full sample number. Black, 7 human and 15 mouse samples (one glial type); grey, 34 human and 74 mouse samples (all glial types together). Lines, mean of 200 random selections of samples; shading, range that contains 95% of the selections. Blue, the values of glial cells (band, single glial types, 0.004 to 0.016; dashed line, all glial types together, 0.064). (B) Statistical power, the share of selections whose median exceeds the 95th percentile of the chance medians, plotted against the level of conservation; dotted line, power of 0.80. Filled circles, samples selected from all neurons; open triangles, samples selected from excitatory neurons only.

**Figure S13.**
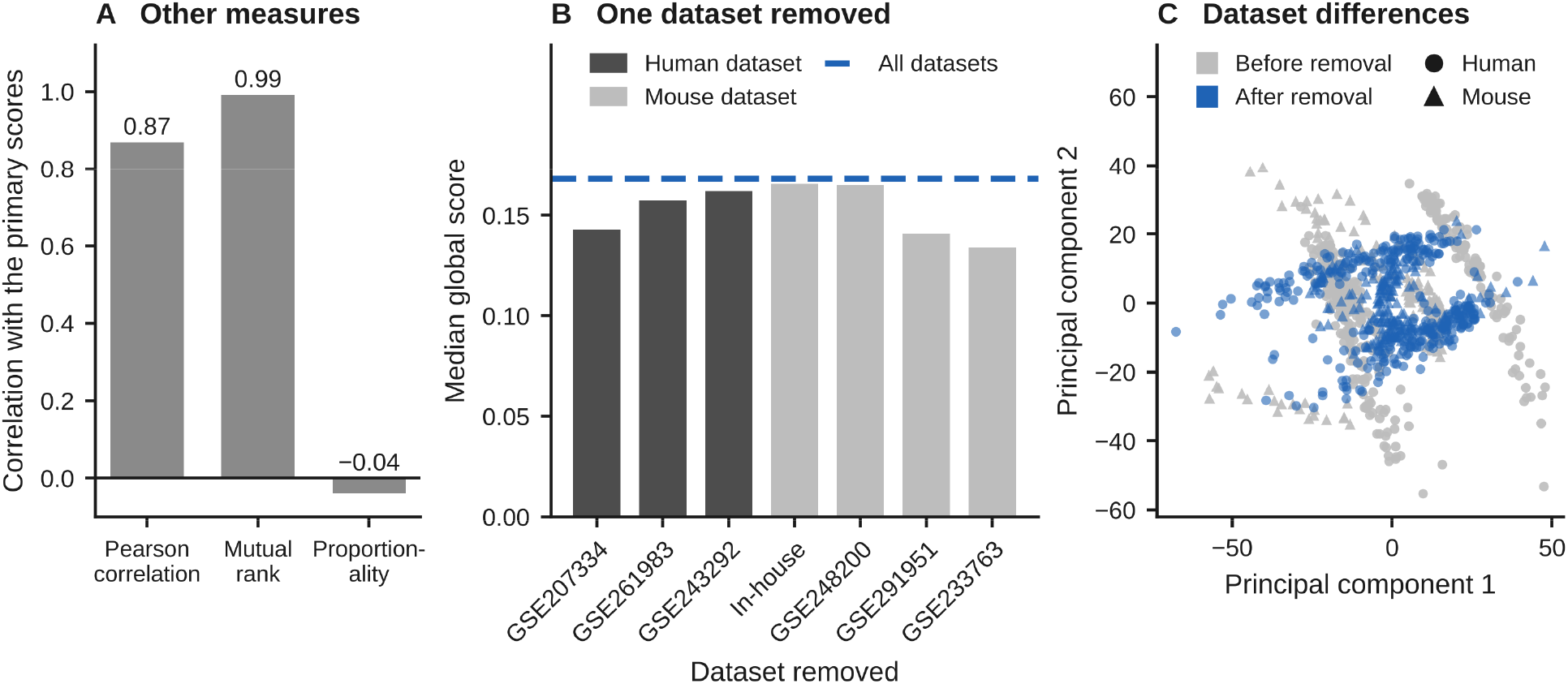
Stability of the scores under alternative analysis choices. (A) Correlation between the global scores of single genes obtained with another measure of association and the primary global scores, which are based on the Spearman correlation. (B) Median global score after removal of each dataset in turn (0.134 to 0.166; dark grey, a human dataset removed; light grey, a mouse dataset removed; dashed blue line, all datasets). (C) Pseudobulk samples plotted on the first two principal components before (grey) and after (blue) removal of differences between datasets; circles, human; triangles, mouse. The components were computed separately for each species and for the data before and after removal, so the four clouds of points share axes only for display.

**Figure S14.**
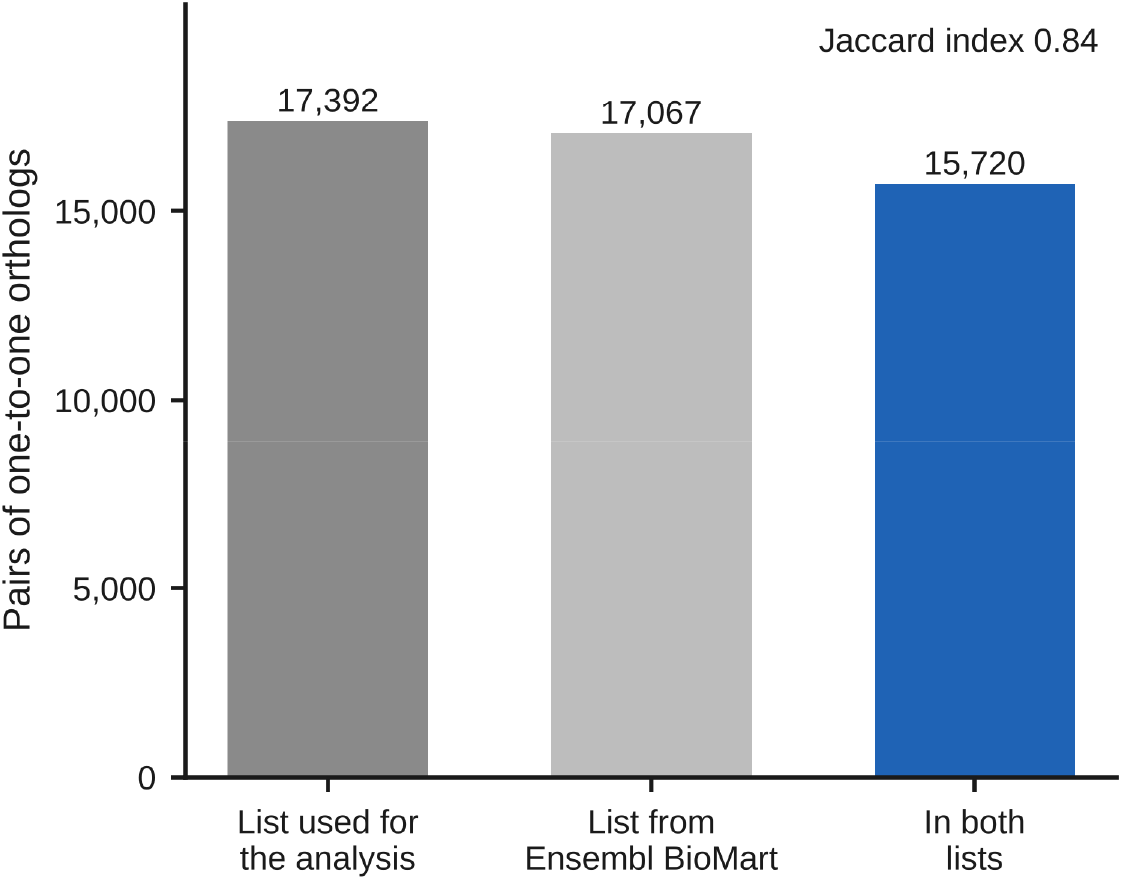
Influence of the list of orthologs. Numbers of gene pairs in the list used for the analysis and in the list newly retrieved from Ensembl BioMart, and the number shared (blue; Jaccard index 0.84).

**Figure S15.**
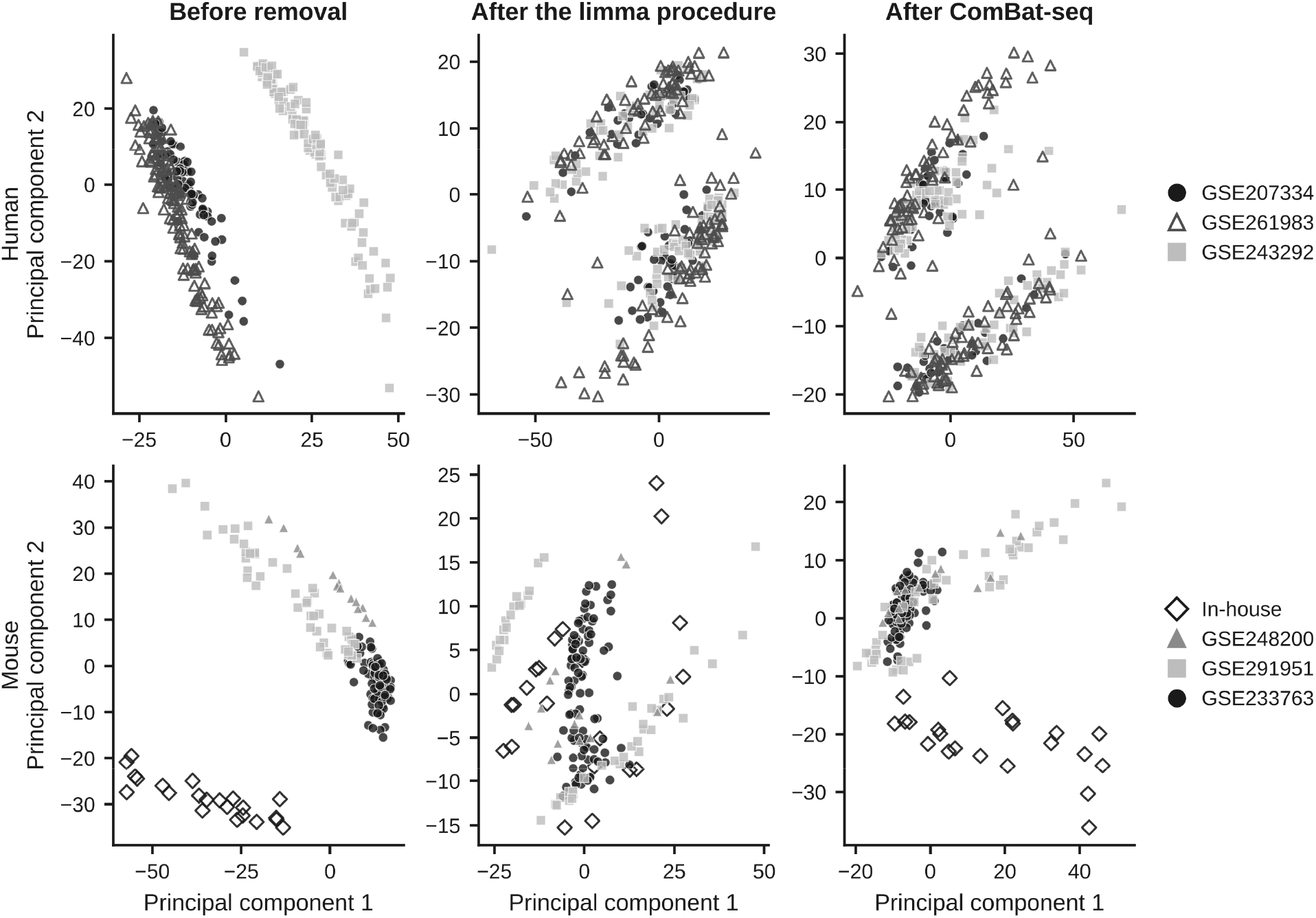
Differences between datasets before and after correction. Human (top) and mouse (bottom) pseudobulk samples plotted on the first two principal components, before removal of differences between datasets, after the limma procedure and after ComBat-seq. Within each species, each dataset has its own symbol (filled or open) and shade of grey. The components were computed separately for each panel.

**Figure S16.**
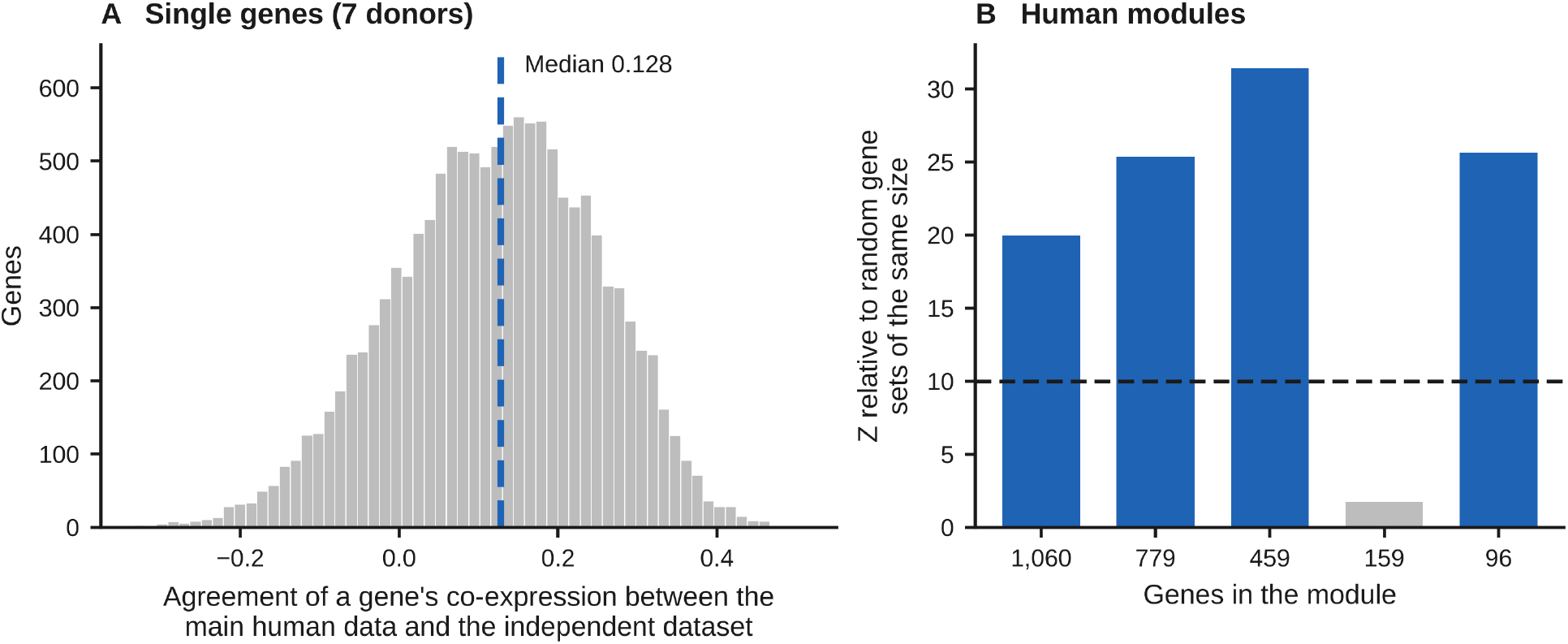
Test of the human network in an independent human dataset (control donors of GSE174367). (A) Agreement between the main human data and the independent dataset for single genes (dashed blue line, median 0.128; seven donors are too few for single genes). (B) Co-expression among the genes of each of the five human modules defined on the 5,000 most variable genes, in the independent dataset, expressed as a Z score relative to random gene sets of the same size; each module is labeled by its number of genes; blue, Z above 10 (dashed line).

**Figure S17.**
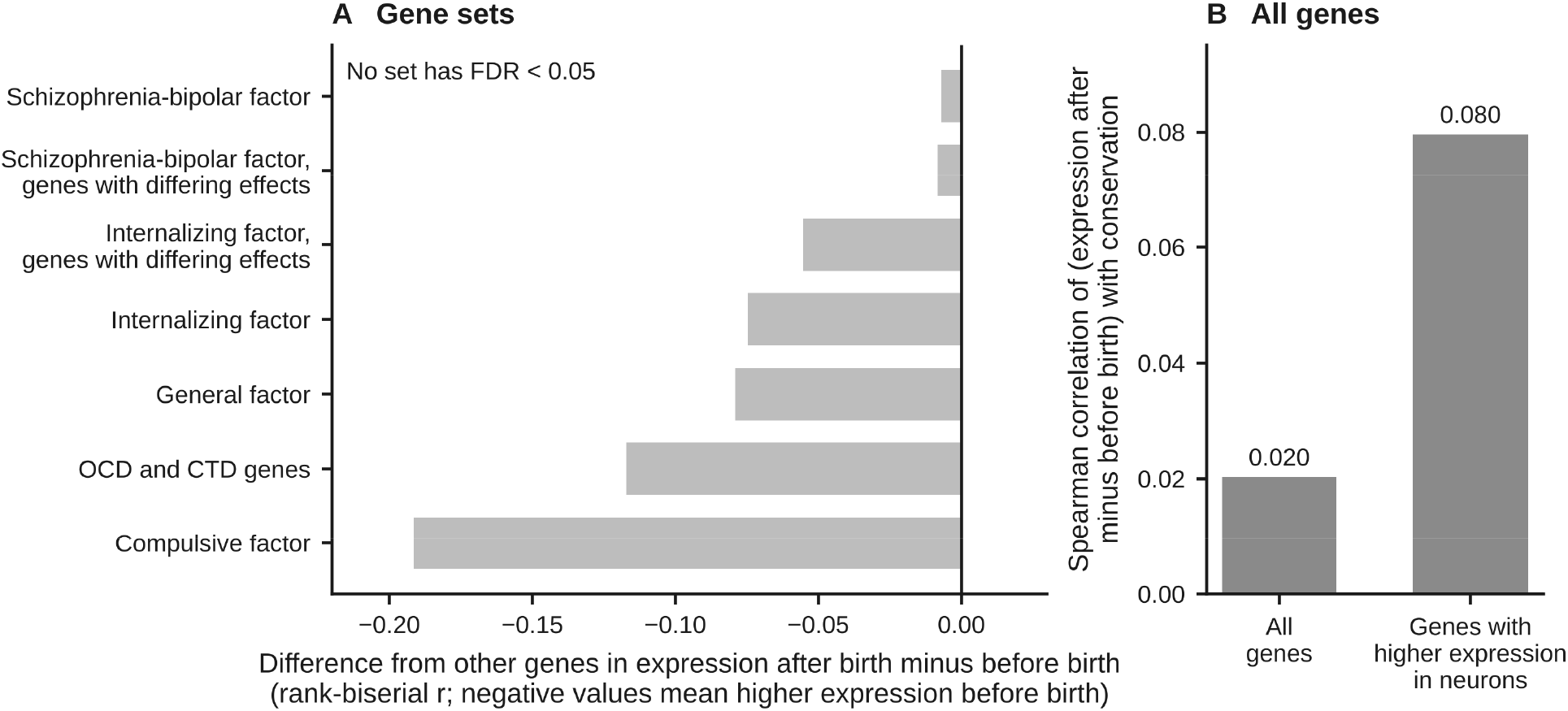
Expression before and after birth. (A) Difference between each gene set and all other genes in expression after birth minus expression before birth in human cortex (rank-biserial correlation, r; negative values indicate higher expression before birth). No set has a false discovery rate (FDR) below 0.05. (B) Spearman correlation between the same quantity (expression after birth minus before birth) and the conservation score, for all genes of the panel and for the genes with higher expression in neurons than in glial cells (Supplementary Methods). OCD, obsessive-compulsive disorder; CTD, chronic tic disorders.

**Figure S18.**
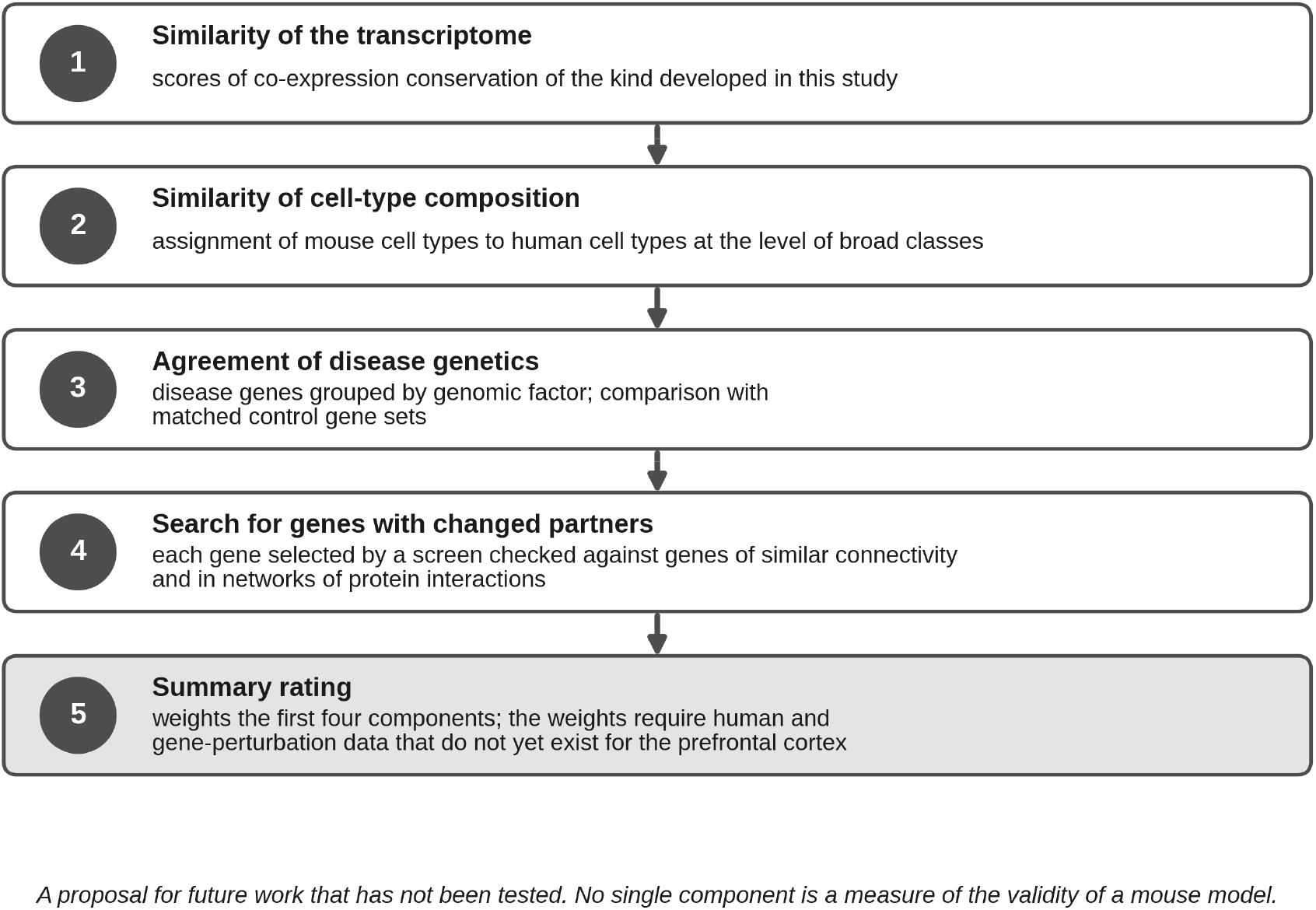
Proposed rating of how faithfully a mouse model reproduces human gene networks. The five components are described in Supplementary Note 9. The rating is a proposal for future work and has not been tested.

**Figure S19.**
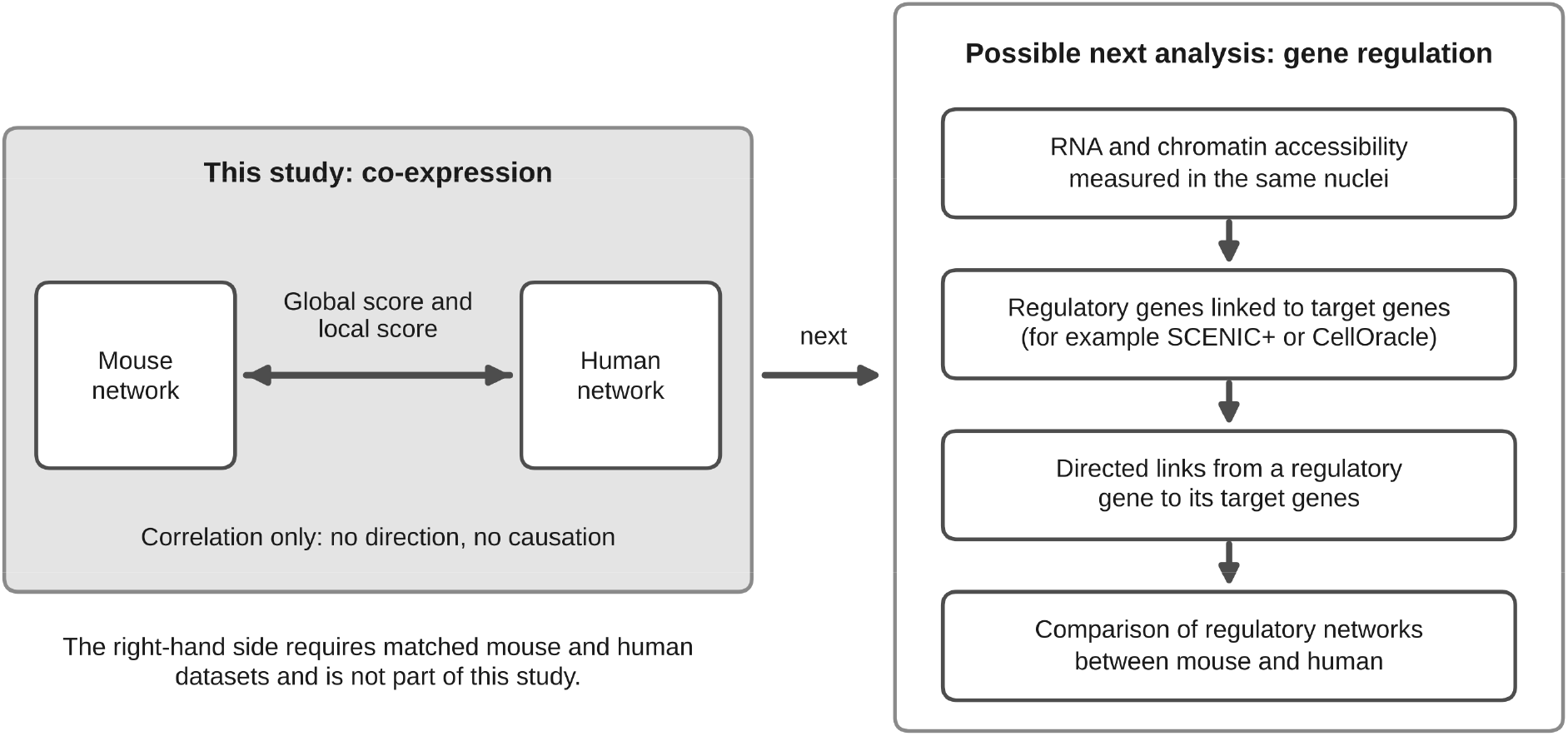
From co-expression to gene regulation. Diagram of a possible next analysis, in which regulatory genes are linked to their target genes using data that measure RNA and chromatin accessibility in the same nuclei, for example with the methods SCENIC+ or CellOracle. The diagram does not show results of this study.

#### Further tables in the analysis package

Scores of all genes in each cell class; comparisons within a species and noise-corrected scores of all genes; tests of disease gene sets in each cell class; the regression model for the combined disease genes and for each set; expression variance of disease gene sets; the genes of each disease set; genes selected by the screen with the four checks and their partners; functional categories of the partners; sensitivity to the thresholds of the screen; principal-component coordinates before and after removal of dataset differences; comparison of ortholog lists and the inclusive panel; assignment of subtypes by marker genes; agreement among measures of association; chance values; network density; curves fitted to the sample-number series and the difference at equal sample numbers; removal of single datasets; scores and module tests in the independent dataset; module preservation in both directions, the control for unassigned genes and the overlap of mouse and human modules. For the genomic factors and the OCD and CTD sets: tests of conservation; overlap between factor sets and combined diagnostic sets; genes with differing effects; ratios of adjusted to unadjusted coefficients and distribution across LOEUF; tests within levels of LOEUF, the interaction test and matched control sets (with the balance of matching and both matching widths); the internalizing factor in glial cells; the simulation of statistical power at glial sample numbers; the tests in protein-interaction networks; the developmental tests; one table with all false discovery rates; the gene properties; and the external gene sets with their sources and versions.

#### Earlier analyses kept in the analysis package

The analyses of disease gene sets and of glial cells that were restricted to the 5,000 most variable genes are kept in the analysis package because they document the two analysis choices that distort results. They are superseded by the analyses reported here and should not be cited as results.

## Notes

### Competing Interest Statement

The authors have declared no competing interest.

